# Molecular elucidation of drug-induced abnormal assemblies of the Hepatitis B Virus capsid protein by solid-state NMR

**DOI:** 10.1101/2022.09.14.507909

**Authors:** Lauriane Lecoq, Louis Brigandat, Rebecca Huber, Marie-Laure Fogeron, Morgane Callon, Alexander Malär, Shishan Wang, Marie Dujardin, Mathilde Briday, Thomas Wiegand, David Durantel, Dara Burdette, Jan Martin Berke, Beat H. Meier, Michael Nassal, Anja Böckmann

## Abstract

Hepatitis B virus (HBV) capsid assembly modulators (CAMs) represent a new class of anti-HBV antivirals. CAMs disturb proper nucleocapsid assembly, by inducing formation of either aberrant assemblies (CAM-A) or of apparently normal but genome-less empty capsids (CAM-E). Classical structural approaches have revealed the CAM binding sites on the capsid protein (Cp), but conformational information on the CAM-induced off-path aberrant assemblies is lacking. We show that solid-state NMR can provide such information, including for wild-type full-length Cp183, and we find that in these assemblies, the asymmetric unit comprises a single Cp molecule rather than the four quasi-equivalent conformers typical for the icosahedral T=4 symmetry of the normal HBV capsids. Furthermore, while in contrast to truncated Cp149, full-length Cp183 assemblies appear, on the mesoscopic level, unaffected by CAM-A, NMR reveals that on the molecular level, Cp183 assemblies are equally aberrant. Finally, we use a eukaryotic cell-free system to reveal how CAMs modulate capsid-RNA interactions and capsid phosphorylation. Our results establish a structural view on assembly modulation of the HBV capsid, and they provide a rationale for recently observed differences between in-cell versus in vitro capsid assembly modulation.

## Introduction

Chronic infection with hepatitis B virus (HBV), a small enveloped retrotranscribing DNA virus, afflicts more than 250 million people. With nearly one million deaths per year, the resulting chronic hepatitis B (CHB) represents one of the most deadly diseases, especially in Africa and the Western Pacific region^1^. While an effective prophylactic vaccine is available, current therapies can rarely cure infection. Owing to severe side-effects, only few patients are eligible for the finite treatment with type-I interferon (IFN), whereas nucleos(t)ide analogs (NAs) inhibiting the viral polymerase usually require life-long administration to keep virus replication suppressed; otherwise, the covalently closed circular (ccc) DNA, the persistent nuclear form of the viral genome, will resume to give rise to progeny virions^2^. Hence multiple efforts are underway to achieve a functional cure, i.e. maintained viral suppression upon therapy withdrawal, or ideally a sterilizing cure, i.e. elimination of the virus^3^. Besides targeting HBV-relevant host factors^4^, novel direct-acting antivirals are being investigated^5^. One of the most advanced new drug classes are capsid assembly modulators (CAMs)^6,7^, sometimes called core protein allosteric modulators (CpAMs). They target the core protein (Cp), the building block of the icosahedral HBV capsid, whose pleiotropic activities are crucial for multiple steps of the viral life-cycle^8,9^, including as a specialized compartment for genome replication^7^. Cp comprises an N-terminal assembly domain (residues 1-140) connected by a nine-residue linker to a highly basic C-terminal domain (CTD; residues 150-183) which binds nucleic acids, notably the viral pregenomic (pg) RNA^10^, the substrate for capsid-internal reverse transcription into the HBV-typical partially double-stranded (ds) relaxed circular (rc) DNA in infectious virions^11^. Cp’s assembly domain features five α-helices, as determined by early cryo-EM and x-ray studies^12,13^ of recombinant capsids from truncated CTD-less Cp. The central helices α3 and α4 from two Cp molecules associate into a four-helix bundle, forming a stable dimer; these dimers then assemble into closed icosahedral shells. The major class comprises 120 Cp dimers arranged with triangulation number T=4 symmetry, a minor class (T=3) consists of only 90 dimers. In the simplest case of icosahedral symmetry, T=1, three structurally identical capsid protein subunits contribute each of the 20 triangular faces of the icosahedron. Larger shells accommodating more than 60 subunits are accessible based on the principle of quasi-equivalence^14^, whereby each face is further subdivided by a certain integer. The respective triangulation (T) number represents the number of similarly but non-identically structured (“quasi-equivalent”) subunits required to adapt to the same number of symmetry-related non-identical environments. Hence for T=4 symmetry there are four different molecules or, for viruses with a single capsid protein like HBV, four conformers in the asymmetric unit, termed A, B, C and D (**Figure 1a**).

**Figure 1:**
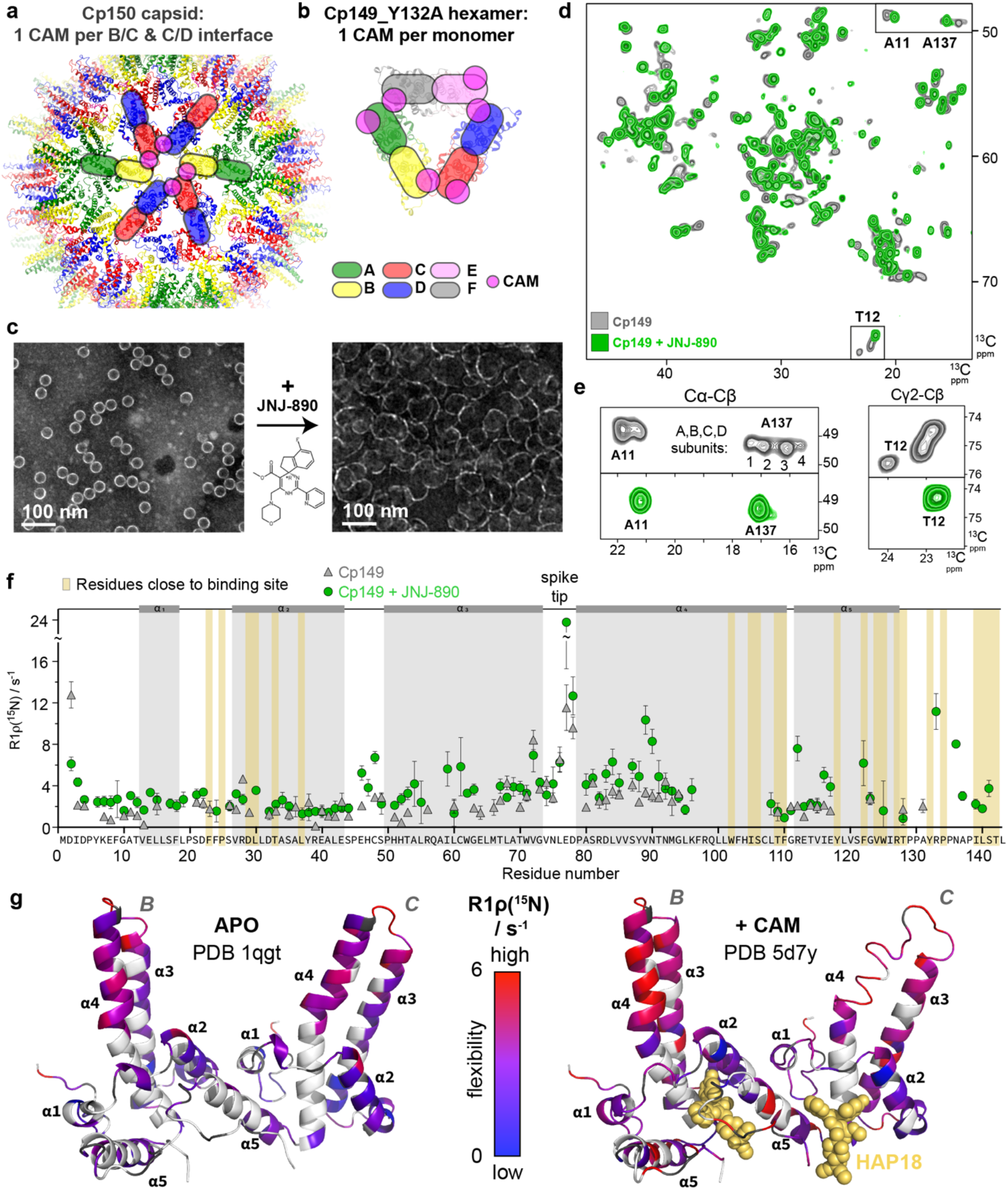
Effect of JNJ-890 binding on Cp149 structure and dynamics. **a)** Representation of the localization of CAM binding sites (pink circles) found in the ABCD subunits of HBV Cp150 capsid stabilized by artificial disulfide bridges involving C150 (*e.g.* PBD 5d7y^32^). **b)** In the Cp149_Y132A crystalline hexamer, each monomer can accommodate one CAM. In addition to the A, B, C and D subunits of T=4 icosahedral capsids, E and F subunits are present in the asymmetric unit of the Y132A hexamer crystal trimer of dimers (*e.g.* PBD 5wre^33^). **c)** Negative-staining EM micrographs of Cp149 NMR samples in the absence and presence of the JNJ-890 CAM-A. **d)** Overlay of aliphatic spectral regions of ^13^C-^13^C DARR spectra of Cp149 in absence (grey, from reference^47^) and in presence of JNJ-890 (green). **e**) Extracts of the peak multiples corresponding to the different conformers in the asymmetric unit of T12 and A137 resonances, and the resulting single resonance in presence of CAM-A. **f)** Rotating-frame relaxation-rate constants R1ρ(^15^N) in the absence and presence of JNJ-890 (for relaxation-rate constant differences see **Figure S3**, and experimental details in **Table S4**). The larger the rate constant, the larger the molecular dynamics on the microsecond time scale. **g)** Experimentally-elucidated dynamics plotted on the apo-capsid structure (PDB 1qgt^13^, left structure) and on a CAM-A-bound structure (PDB 5d7y^32^ with HAP18 in yellow spheres, right structure), where red indicates high amplitude motions on the microsecond timescale and blue low amplitude motions. Prolines are colored in grey and residues with unavailable data in white.

Cp plays important roles in almost every step of the HBV life cycle, including pgRNA co-encapsidation with the viral polymerase, enabling capsid-internal pgRNA reverse transcription into relaxed circular (rc)DNA, signaling for capsid envelopment with the surface proteins, and virion secretion. Also, the capsid carries a continuous flow of rcDNA into the nucleus *via* the core protein’s nuclear localization signals located in the CTD. With few recent exceptions^15,16^ most structural studies were done on CTD-less Cp variants, hence the underlying structural dynamics are not yet understood in detail, but they must be tightly regulated. Interfering with proper nucleocapsid assembly and its regulation is therefore a promising antiviral strategy (see reference^7^ for a recent review).

The first small molecule capsid assembly modulators (CAMs) were phenylpropenamide derivatives such as AT-130^17^ which promote formation of pgRNA-less empty capsids, and heteroaryldihydropyrimidines (HAPs), e.g. BAY 41-4109^18^ which can induce formation of aberrant, nonspherical multimers which are neither functional in virus replication^19,20^. These in vitro phenotypes, though exerting some concentration dependence^21^, are prototypical for the two main categories of CAMs: on one hand, CAM-E compounds, such as phenylpropenamides and sulfamoylbenzamides including JNJ-632 and JNJ-827, that cause assembly of empty capsids of apparently normal shape; on the other hand, CAM-A compounds such as HAPs, including JNJ-890, that lead to aberrant assemblies^22–25^. An earlier classification into class I and class II CAMs was applied in a non-uniform manner and led to confusion in the past^26^.

As neither empty nor aberrant non-spherical particles are compatible with containment of the pgRNA - viral polymerase complex, both CAM classes prevent the formation of rcDNA. Moreover, CAMs have been proposed to decrease the formation of cccDNA and to restore innate signaling, which is not achieved by current NA treatments^22^. CAMs are thus considered as some of the most potent candidates for a curative hepatitis B treatment^6,7^, and several compounds are in clinical trials^26,27^.

Structures of capsids in the presence of CAMs have been determined both by X-ray and cryo-EM^28–33^. Accordingly, despite their different impact on overall particle morphology, both types of CAMs bind to the same hydrophobic cavity at the Cp dimer-dimer interface, called the HAP-binding pocket (**Figure 1a-b**). It is described to consist of a concave interface involving residues F23, P25, D29, L30, T33, L37 (comprising mainly helix 2), W102, I105, S106, T109, F110, Y118, F122, I139, L140 and S141 (comprising helices 4 and 5 and linker region) from one chain, and the region from residue V124 to P134 (helix 5 and linker region) from the other chain^6^.

Notably, however, as both X-ray crystallography and cryo-EM require regularly ordered specimens, the aberrant Cp assemblies induced by CAM-A compounds are intrinsically unsuited for these high-resolution analyses. Hence, in all these studies, recombinant Cp mutants were used which either enabled crystallization in a non-particulate planar hexagonal form *via* the capsid assembly-preventing Y132A mutation^25,30,34^, or artificially forced to maintain near-icosahedral symmetry through intra-capsid cross linking of extra cysteine residues added as residue 150 to the C terminus truncated Cp149 (Cp150)^21,28,29,32^. In the Y132A mutant, each of the three dimer-dimer interfaces in the hexamer accommodates one CAM (**Figure 1b**), while in the Cp150 capsid, only one CAM binds per dimer in the B and C subunits (**Figure 1a**); how this relates to CAM binding to wild-type Cp remains to be determined, because as the aberrant, irregular assemblies observed upon addition of CAM-A to wild-type Cp149 and alike have not been investigated by any direct structural approaches. Furthermore, truncated Cps lacking the entire CTD were used in nearly all studies, hence very little structural information is available for the full-length wild-type Cp183 protein, and whether and how CAM binding is affected by the CTD.

Solid-state NMR can structurally analyze large protein assemblies without the need for crystals or symmetry, as has been shown for protein fibrils (reviewed e.g. in^35^), membrane proteins (reviewed e.g. in^36^), molecular machines (e.g.^37,38^) and viral capsids and envelopes (reviewed e.g. in^39^). NMR chemical shifts are highly sensitive environmental indicators, and can report on conformational variations and molecular binders^39,40^. We here used this approach to investigate the conformational impact of CAMs on the irregular assemblies which form in their presence, including from full-length wild-type Cp183 with RNA, and also the phosphorylated, nucleic acid-free form P7-Cp183 (see **Table S1** for the full list of forms investigated). In addition to incorporation of CAMs into preformed capsids, we also used wheat-germ cell-free protein synthesis (WG-CFPS) to study the effects of CAMs on Cp when present in situ during assembly upon Cp exit from the ribosome. Our study provides thus a molecular-level observation of CAM assembly modulation for full-length Cp carrying a functional CTD. It shows that the data on the impact of CAM on Cp149 capsids do not tell the whole story, and due caution should be exercised in applying them to full-length capsid proteins.

## Results

### CAM-A induce Cp assemblies with a single conformer

Negative staining EM of Cp in the presence of CAM-A compounds has revealed irregularly-shaped structures, for example induced by HAP compounds^41,42^. For illustration, **Figure 1c** (right panel; as opposed to intact capsids in the left panel) shows such micrographs of Cp149 capsids in presence of JNJ-890, a CAM-A compound. The mesoscopic heterogeneity of the observed capsids made us suspect that NMR spectra might show severe heterogenous line broadening, representing a distribution of different protein conformations. In order to assess the structural features at a molecular level, we recorded 2D Dipolar Assisted Rotational Resonance (DARR) spectra (**Figure 1d**, **Table S2**) and hNH (**Figure S1**, **Table S3**) nitrogen-proton correlation spectra of Cp149 capsids in presence of JNJ-890 (in green), and compared them to spectra without CAM-A (in grey). The spectra surprisingly demonstrate that the aberrant objects induced by the CAM-A compound actually yield well-resolved resonance lines in the solid-state NMR spectra, with lines which are even narrower than in the absence of CAMs. Of note, whether CAMs were added to Cp149 preassembled capsids, or to Cp149 dimers before capsid assembly, made no difference for the resulting spectra (**Figure S2**), as also observed by EM^43^. Sequential resonance assignments using 3D NMR following previously described protocols^44,45^ revealed only a single peak per atom for the CAM-A capsid spectra, while we previously described for the apo form that many amino-acid residues displayed resolved peaks for the four different molecules in the asymmetric unit, A, B, C and D of the T=4 icosahedral capsid^46^. We have shown that the resulting peak splitting is strongest for amino-acid residues located at the dimer-dimer interfaces^46^, where pentamers and hexamers have in total four slightly distinguishable monomer conformations^13^. The splitting becomes smaller than the linewidth for residues remote from the dimer-dimer interfaces, as for example the amino acids in the spike, resulting only in a certain line broadening. This pattern is lost in CAM-A capsids, as illustrated in the 2D NMR spectra of **Figure 1e** for the example of T12 and A137 amino-acid residues, both located at the dimer-dimer interface. Quite unexpectedly, the CAM-A Cp spectra are thus simplified with respect to the apo Cp, and the peaks are particularly narrow. This behavior was observed for all 28 residues which showed peak splitting in the apo form. The flattening out observed in the EM micrographs of the assemblies suggests that the lattice is closer to hexagonal, since such types of lattices form rather flat superstructures, whereas the pentameric axes in icosahedral structures introduce curvature. That the observed objects are not fully flat might indicate that a few pentamers are still present, but at an abundance of below approximately 5 % they would be below the NMR detection limit of our NMR experiments. This implies that in the Cp149-JNJ-890 complex, the remaining pentameric structures are present at most at this frequency. However, the observed curvature may also have a different origin.

In order to assess the impact of the CAM-A on capsid dynamics, we measured NMR relaxation-rate constants *R*1ρ(^15^N) of Cp149 in the presence and absence of JNJ-890. These constants report on the internal flexibility of the NH vector of each amino-acid residue on the microsecond time scale. Overall it appears that the relaxation-rate constants, i.e. the flexibility on the microsecond time scale, increase upon drug binding in most parts of the protein, as shown in **Figure 1f** (for a plot of the rate-constant differences see **Figure S3**). In particular, the capsid spike acquires significant new mobility in presence of JNJ-890, as shown on the Cp structures in **Figure 1g**. In addition, smaller effects are detected in the spike base. Some regions remain mostly unaffected in terms of dynamics, especially helix α2, as well as the end of helix α4’ and the beginning of helix α5. Interestingly, residues close to the flexible linker and the CAM binding site (region 130-142) became more intense upon addition of JNJ-890, possibly through collapse of the split peaks, or following a stabilization upon interaction, enabling the measurement of more data points in this region.

In summary, we surprisingly observe, for the abnormal assemblies resulting from addition of a CAM-A compound, very high-quality NMR spectra. They interestingly reveal a collapse of the formerly split peaks in T=4 icosahedral Cp into single peaks, and also show increased dynamic behavior, notably in the spike region.

### NMR fingerprints directly pinpoint CAM-proximal residues

While negative-staining EM can distinguish between aberrant Cp assemblies formed in the presence of CAMs and normal capsids, NMR spectra allow to obtain a detailed fingerprint of the interacting residues, and to detect changes in local symmetry, including a different asymmetric unit. **Figure 2** shows EM micrographs and sections of NMR spectra (for full aliphatic regions see **Figure S4**) for Cp149 with a variety of CAMs for which the chemical structures are given. While in *apo* Cp149 (top of **Figure 2**) capsids, four different A137 and two different A11 peaks are clearly distinguished, the number of peaks decreases when CAMs are added. For GS-942049, GS-832471, and JNJ-827, it can be seen that two out of the previously four peaks disappear. For JNJ-632 and HAP_R10^48^, the two peaks start to coalesce. This points to the dimers, which start out as parts of the hexameric and pentameric units in the context of icosahedral symmetry, becoming more similar to each other. Finally, for JNJ-890, only a single major peak is left, indicating a virtually single conformation. It should be noted that the single peak for the JNJ-890 Cp finally indicates full occupancy with CAM on all sites. **Figure 2c** shows the residue-specific chemical-shift differences between the *apo* and CAM-bound capsids for the different compounds plotted on the Cp dimer 3D structure (PDB 5d7y^32^) (for chemical shift perturbations (CSPs) see **Figure S5a**). CSPs allow to directly identify the residues which are most impacted by the interaction with a CAM. A number of similar CSPs is observed for both CAM-A and CAM-E, highlighted in yellow in **Figure S5**. Interestingly, F24, L30 and P129 show significantly larger CSPs for CAM-A than for CAM-E (green in **Figure S5**), and N136 and P138 show large CSPs only in presence of CAM-E (pink in **Figure S5**). Large CSPs in these resonances can thus be considered as discriminators for CAM-E vs CAM-A compounds. Interactions between the CAMs and these residues might be responsible for the divergent action of the two classes of molecules. The largest number of significant CSPs is observed for the CAM-A compounds, likely since CSPs not only reflect the binding of CAMs, but also changes in symmetry. Notably, the CSPs observed for CAMs differ from those observed for Triton-X100^47^, which binds to the hydrophobic pocket located at the intra-dimer interface, whereas CAMs bind to the hydrophobic cavity located at the inter-dimer interface.

**Figure 2:**
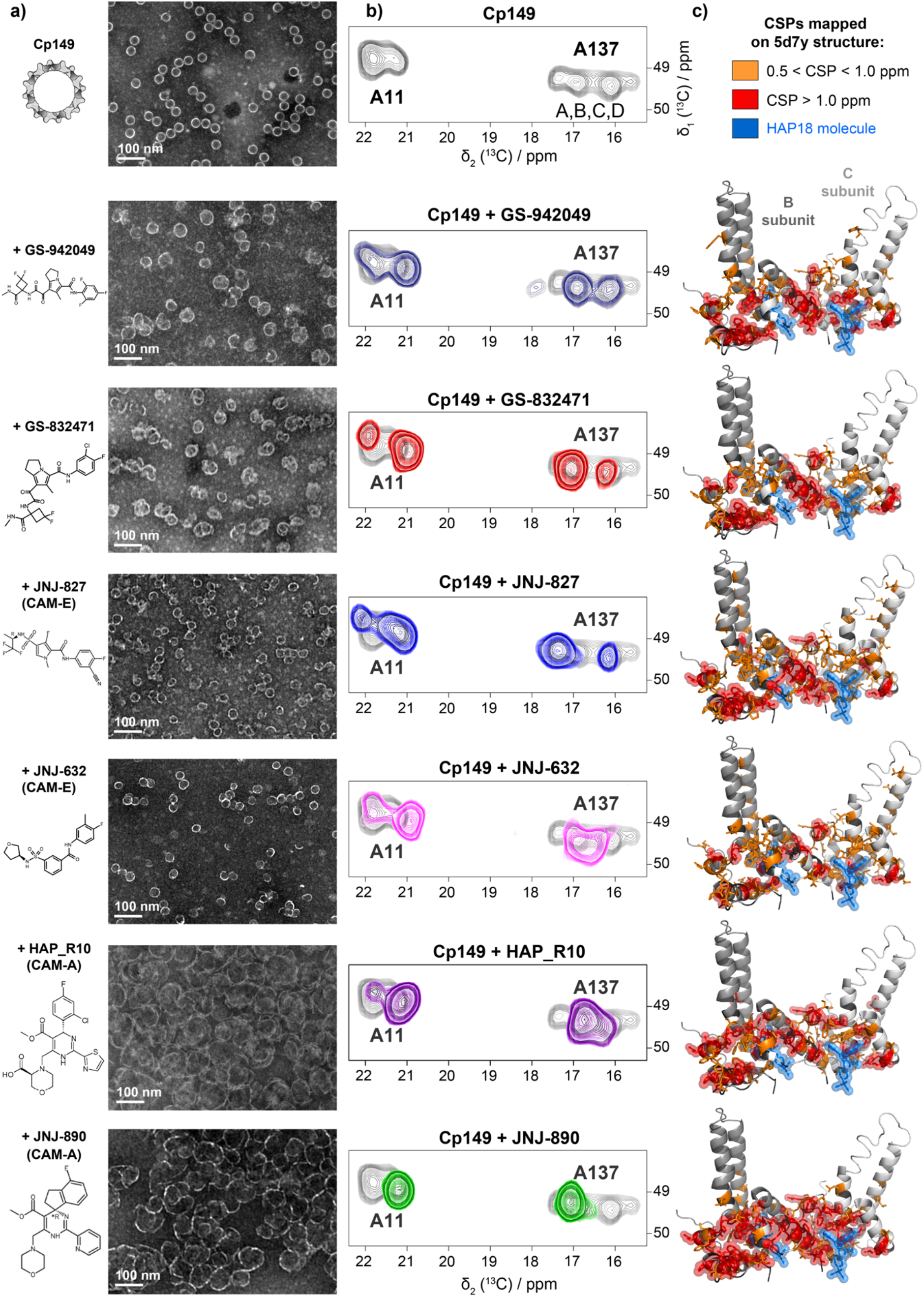
CSPs induced by different CAM-A and CAM-E compounds. **a)** Negative-staining EM micrographs corresponding to the ^13^C-^15^N Cp149 reassembled capsids in absence (for reference) or presence of CAM, with their chemical structure. **b)** Extracts from the NMR DARR spectra showing the A11 and A137 alanine Cα-Cβ region for all tested compounds. In light grey is the control spectrum for comparison (from reference^45^). For full aliphatic regions of DARR spectra, see **Figure S4**. **c)** CSPs induced by the different CAMs mapped on Cp149-HAP18 structure (PDB 5d7y^32^) where unaffected residues are shown in grey, and residues affected by the binding are colored orange (medium CSPs, in sticks) and red (strong CSPs, in spheres). HAP18 is represented as blue spheres. For CSP graphs for all residues see **Figure S5** and values in **Table S5**).

To summarize, NMR can distinguish different classes of CAMs by their spectral fingerprint: CAM-E show distinct patterns of multiple peaks, while the aberrant assemblies resulting from CAM-A show a major single resonance per amino acid, and the largest CSPs, also affecting the largest number of residues. This atomic-level fingerprint presents an interesting complement to kinetic methods (e.g. microfluidic screening^49^) used to distinguish CAM-A from CAM-E compounds.

### CAM-A bound Cp183 and Cp149 are different at a mesoscopic but identical at the molecular level

A question not answered before is how CAMs affect, at the molecular level, the structure of capsids assembled from the full-length Cp183 protein. We thus analyzed, in addition to truncated Cp140 and Cp149, Cp183 and phosphorylated P7-Cp183, prepared by sedimentation^50,51^ of the capsids into the NMR rotor in the presence of JNJ-890 CAM-A. **Figure 3a** shows the EM micrographs of the different capsids after incubation with JNJ-890. The superstructures formed are for Cp140 and Cp149 clearly open, while full-length capsids seem nearly unaffected, also at higher Cp:CAM-A ratios (**Figure S6**). Notably, P7-Cp183 capsids (**Figure S6c**) remained mainly circular, even if some seem to have disrupted contours. Even at very high ratios, up to 1:20, their diameters seem to increase slightly, but they do not substantially open (**Figure S7**). In contrast, Cp149 already at a ratio Cp:JNJ-890 of 1:1 has nearly disassembled (**Figure S6b**). When analyzing changes at different points in time (**Figure S8**), Cp183 capsids started to detectably open up after one month of storage, but still showed a more compact structure than Cp149 with CAM-A. Altogether this indicates that there are additional inter-dimer interactions in Cp183 which maintain a roughly spherical structure of the multimer. It is interesting to note also that CAM-A seems to bind preferentially on open assemblies, as can be seen when sub-stoichiometric amounts are added in **Figure S6** and **Figure S8**, where intact capsids are observed next to flat assemblies.

**Figure 3:**
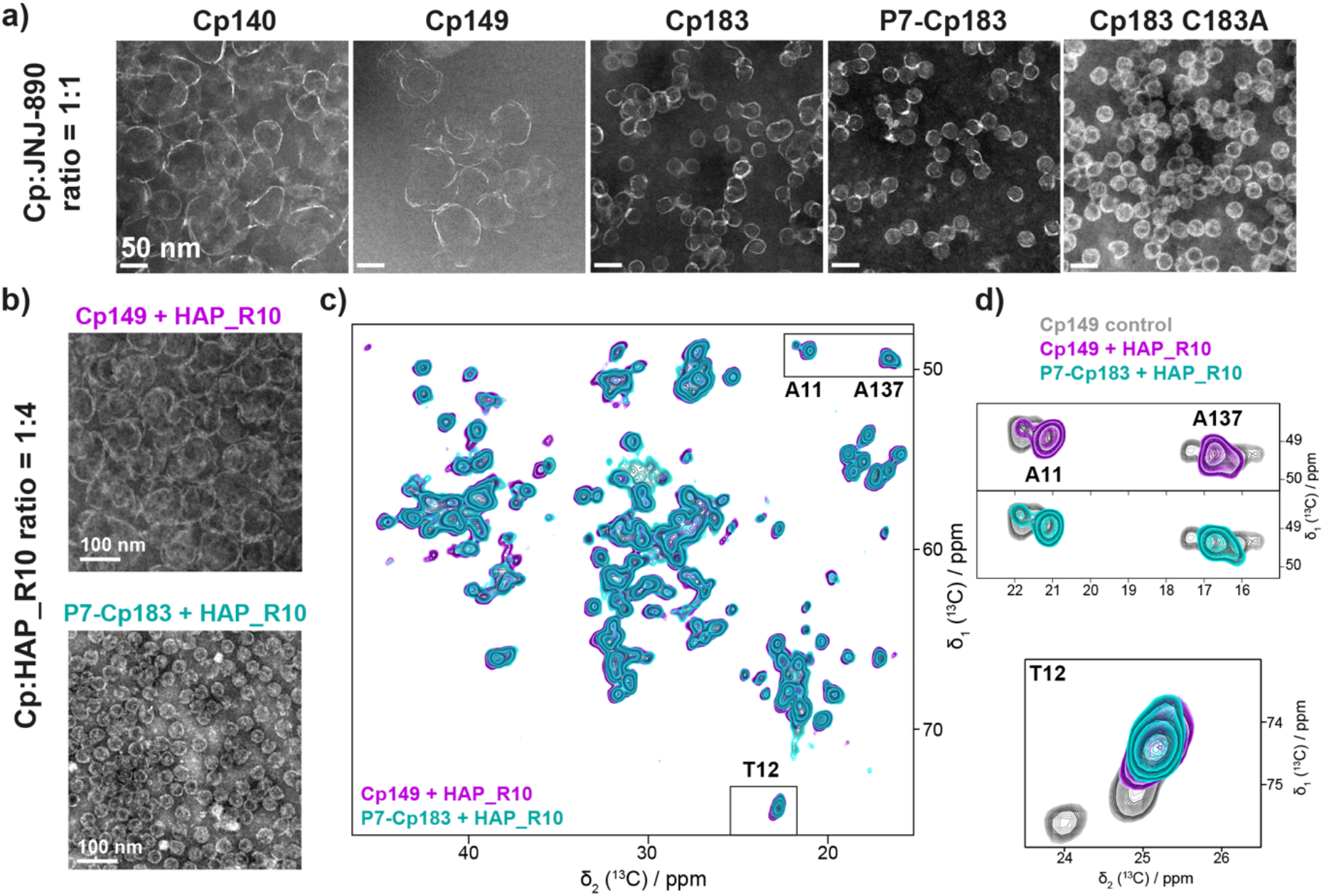
Effect of CAM-A at the mesoscopic and atomic level on truncated and full-length HBV core protein. **a)** Negative-staining EM micrographs of different capsid constructs after incubation with 1-molar equivalent of the JNJ-890 CAM-A. Capsids made of truncated Cp149 result fully open, while those assembled from full-length Cp183 appear mostly closed. For EM micrographs at four different JNJ-890 ratios see **Figure S6**. **b)** EM micrographs corresponding to the NMR samples of Cp149 and P7-Cp183 bound to HAP_R10. **c)** Overlay of ^13^C-^13^C DARR solid-state NMR spectra of Cp149 (purple) and P7-Cp183 (cyan) bound to HAP_R10. **d)** Zooms from the DARR spectra showing A11, A137 and T12 resonances, with the reference Cp149 spectrum in grey (from reference^47^).

In order to assess whether these differences on the mesoscopic level between open and closed capsids are reflected at the molecular level, we recorded NMR spectra on Cp183 and P7-Cp183 capsids in the presence of JNJ-890 and HAP_R10 CAM-A (**Figure S9**). We have previously shown that the spectra of the Cp149, Cp183 and P7-Cp183 apo forms virtually coincide^47^. The spectra of Cp149 and P7-Cp183 with HAP_R10 (EM in **Figure 3b**, NMR in **Figure 3c**, and zooms in **3d**), reveal that this is surprisingly also the case for the HAP_R10-bound capsids. The same observation is made for Cp183 with JNJ-890, as shown in the spectra in **Figure S10**. Notably, in both Cp183 and P7-Cp183 preparations in presence of CAM-A, a small residual signal could be observed, either stemming from pentamers that might still be present at NMR detectable levels/ratios, or from CAM-free sites, or both.

We realized when analyzing different NMR rotor contents of Cp/CAM-A preparations by EM that 1,4-dithiothreitol (DTT) can enhance capsid opening when used on Cp183 and P7-Cp183. Indeed, while all samples had yielded the same NMR spectra, appearance under EM was quite different, and correlated with the presence of DTT in the buffer. DTT alone had no impact on the NMR spectra of Cp besides reducing the peak intensity for oxidized Cys^47^. **Figure S11** shows the impact of DTT on Cp183 capsids in presence of JNJ-890. While Cp183 were largely spherical at a Cp:CAM 1:1 molar ratio, they were heavily deformed or even disassembled when DTT was present, similar to the effect observed on Cp149+CAM-A in absence of DTT (**Figure 3a**).

We subsequently tested whether a straightforward interpretation of DTT opening disulfide bonds, notably a bond involving C183 and C48 as postulated recently^52^, or the bonds present in the Cp150 with C-terminal Cys used for cryoEM^21,28,29,32^, might hold, through using different Cys mutants (**Figure S11**). Interestingly, notably the Cp183 C183A mutant showed no opening in presence of CAM-A, as also observed for wild-type Cp183. This excludes a stabilizing effect of this cysteine as cause for increased capsid integrity, and also confirms that CTD-meditated stabilization is not due to oxidative crosslinking and formation of a covalent intra-capsid network^53^. All other Cp183 cysteine mutants also resisted to disassembly in presence of CAM-A. Upon addition of DTT, Cp183 C183A readily disassembled, while C107A, and to a lesser extent C61A, were least affected by addition of DTT, suggesting that in Cp183 WT the mutated Cys residues are somehow involved in stabilizing the spherical capsid-like structure in a DTT-sensitive fashion, but not by S-S bond formation, which are sterically not possible. However, how this might work on a molecular basis remains unclear, and no simple picture emerged from our experiments.

Another capsid-stabilizing factor in CTD-containing vs. CTD-less Cp variants can be packaged RNA^54–56^, as its multivalent electrostatic interactions with the positively charged CTDs could also counteract particle disruption. Similarly, interactions between phosphorylated CTDs and positively charged arginine residues in P7-Cp183 can stabilize capsids ^57^. Still, how DTT could impact these interactions remains to be determined.

Finally, we assessed the packaged RNA content of *E. coli* Cp183 capsids incubated with both types of CAMs. For this, we compared the ^13^C RNA NMR signals in the 2D DARR spectra recorded on Cp183 capsids with and without CAM (**Figure S12**). Through its arginine-rich domain, Cp183 can package the equivalent of roughly 4,000 nucleotides of *E. coli* RNA^54,56,58^. Both CAM-A compounds reduced the RNA signals by around 25 % for JNJ-890, and 50 % for HAP_R10 in the aberrant assemblies, which either indicates that RNA was released from the capsid CTDs, or that RNA is more mobile, which would result in reduced dipolar polarization transfers. In contrast, the JNJ-632 CAM-E had no significant impact on the RNA signal intensity, meaning that the RNA remained protected in the non-disassembled capsids, as expected for CAM-E.

In summary, we conclude that, despite the differences observed by microscopy, Cp149, Cp183 and P7-Cp183 all undergo similar molecular-level changes in response to the JNJ-890 CAM-A. At the mesoscopic level, the opening of capsids in the presence of CAM-A is in a yet unexplained manner assisted by DTT.

### Modulation by CAMs directly during capsid assembly

Directly investigating re-assembly of Cp183 dimers in the presence of CAMs is difficult, as it often leads to aggregation of the protein. As an alternative, we have recently shown that capsid assembly of Cp183 in the presence of mRNA and CAMs can be achieved by using wheat-germ extract cell-free protein synthesis (WG-CFPS)^59^. We here use this approach to investigate whether capsid assembly directly upon Cp183 exit from the ribosome is modulated in the same way as on preformed capsids, and how nucleic acid packaging and phosphorylation are orchestrated in presence of CAM. One should mention that DTT is systematically present in the cell-free reaction^60^. We synthesized isotope-labeled Cp183 capsids in the cell-free system (in the following referred to as CF-Cp183) in the presence of different CAMs. By negative-staining EM of the assemblies directly in the crude CFPS reactions carried out with increasing amounts of CAM-A, we found that a Cp monomer:JNJ-890 ratio of 1:0.4 was sufficient to open most capsids (**Figure S13**), which was therefore chosen for NMR sample preparation with JNJ-890 and HAP_R10 CAM-A. For the JNJ-632 CAM-E, a 1:1 molar ratio was used (**Figure S14**). Interestingly, while CF-Cp183 is, as in absence of CAM (**Figure S14a**), largely found in the soluble fraction in presence of JNJ-632 (**Figure S14b**), it localizes to a large part to the insoluble pellet fraction with JNJ-890 and HAP_R10 (**Figure S14c,d**).

The resulting EM micrographs and 1D ^31^P and 2D hNH spectra are shown in **Figure 4**. As observed with *E. coli*-produced Cp149 and Cp183, JNJ-632 gave rise to mesoscopically normal capsids, whereas both CAM-A compounds induced large aberrant assemblies that appeared more open with JNJ-890 than with HAP_R10 (**Figure 4a**). It can be noted that even Cp183 capsids open in the presence of CAM-A when synthesized in the CF system. This is likely due to the presence of DTT in the CF reaction, which we have shown above has an impact on the morphology of Cp183 in presence of CAM-A (see above and **Figure S11**). In line with the DARR spectra of the *E. coli* samples, the hNH spectra recorded with both CAM-A compounds lacked peak multiplicity, indicating a loss of T=4 icosahedral symmetry (**Figure 4c**). Moreover, the spectra from both the bacterial and cell-free samples were virtually superimposable, (**Figure S15**), indicating the absence of local conformational differences.

**Figure 4:**
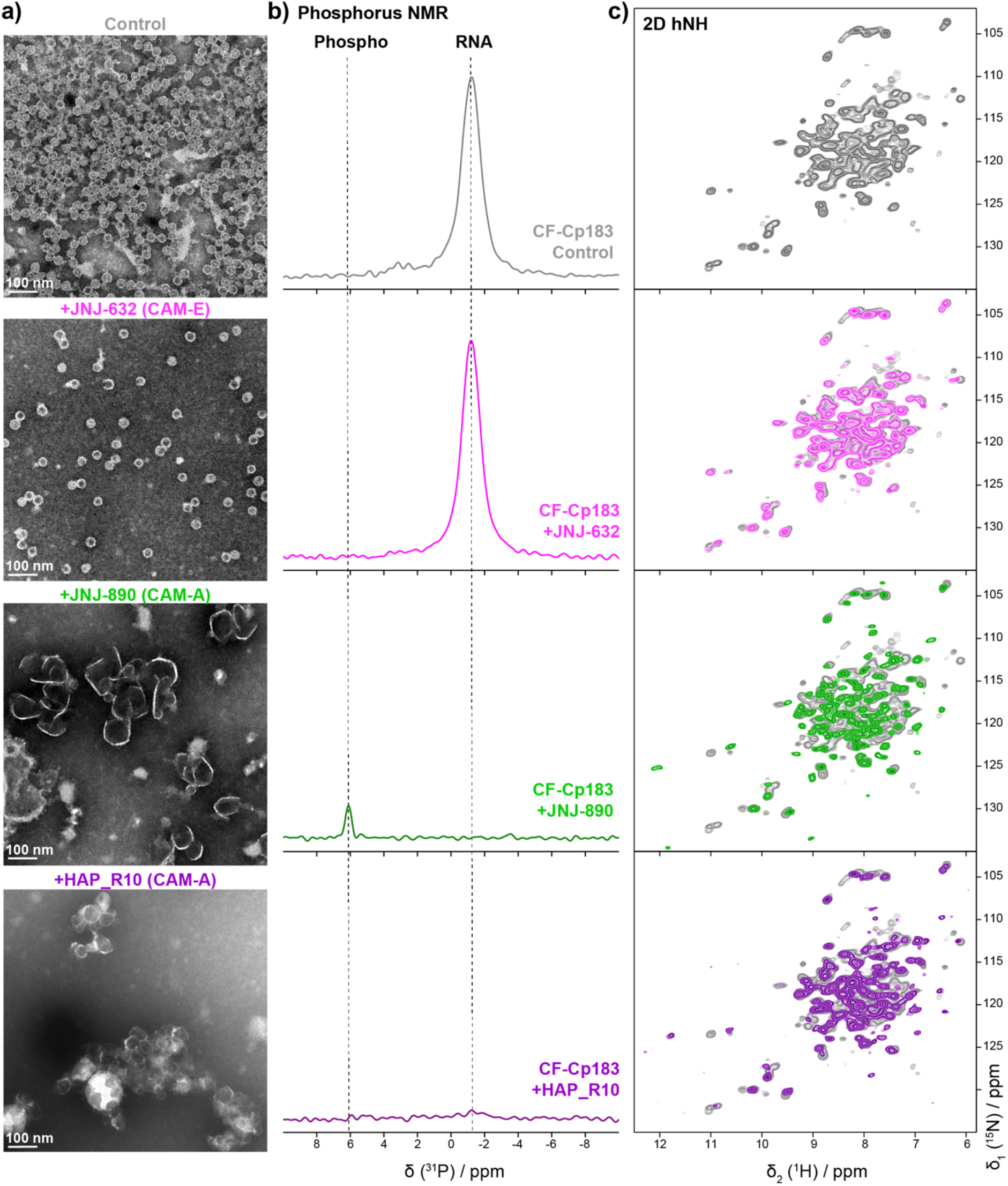
Effect of CAM-A and CAM-E on Cp183 CTD phosphorylation and RNA packaging. **a)** Negative staining EM micrographs of NMR samples of Cp183 from WG-CFPS as control. EM micrographs were taken on resuspended sediments after 1.3 mm rotor filling (see corresponding SDS-PAGE in **Figure S13**). **b)** 1D ^1^H-^31^P CP spectra of CF-Cp183 without CAM (grey), assembled in presence of 1 molar equivalent of JNJ-632 (pink), and 0.1 molar equivalent of JNJ-890 (green) and HAP_R10 (purple) recorded at 55-60 kHz MAS on a 500 MHz spectrometer. **c)** 2D hNH spectra of the corresponding samples recorded at 60 kHz MAS on a 800 MHz spectrometer. Spectra of the control sample are shown in light grey (from reference^45^) for comparison. Corresponding CSPs are shown in **Figure S15**.

We have recently shown that the WGCF system supports phosphorylation of viral proteins ^61–63^, and were therefore able to analyze the phosphorylation state of CF-Cp183 in the presence of CAMs. The ^31^P spectrum of CF-Cp183 in absence of CAM indicates that no phosphorylation occurred in these capsids (since no ^31^P signal at 6 ppm is observed), while clear signals for Cp-associated RNA were seen around −1.5 ppm (**Figure 4b**) (the RNA packaged by the capsid during cell-free synthesis is the mRNA coding for the protein^59^). This result is in line with data for *E. coli* produced Cp183 capsids^56^ which package RNA when unphosphorylated, but not when highly phosphorylated, as in P7-Cp183. The same experiment done with addition of JNJ-632 reveals a similar RNA content as judged by the intensity of the signal, and also absence of phosphorylation. Hence RNA is packaged into capsids in the presence of a CAM-E compound when assembled in the cell-free system, although in mammalian cells pgRNA content was found to be substantially reduced in the presence of other CAM-E compounds, such as AT-61^64^. In contrast, when JNJ-890 was added during cell-free protein synthesis, no RNA signal was detected in the ^31^P spectrum, while a signal typical for phosphorylated amino acids was detected around 6 ppm^47^. The signal for phosphorylation was weaker than the one detected for RNA in the CAM-E samples, despite comparable NMR spectrum acquisition times. Explanations could include a smaller number of ^31^P atoms in the phosphorylated proteins than in the equivalent of about 4,000 nucleotides of RNA (if the RNA content is similar to the one observed in *E. coli* ^54,56,58^), or a higher mobility of the phosphorylated Cp183 CTD when compared to the RNA phosphate backbone, leading to a less efficient cross-polarization transfer and hence reduced peak intensity in the NMR experiment. Clearly, no rigidly bound RNA can be detected in CAM-A Cp assemblies.

In order to assess the extent of phosphorylation, mass spectrometry of capsids assembled in the presence of JNJ-890 was performed, and revealed eight-fold phosphorylation, and no signal for unphosphorylated protein (**Figure S16**). The fact that eight phosphorylated sites are observed suggests that there are more kinases present in the WGE than just SRPK1, whose coexpression in *E. coli* results in phosphorylation of seven sites^56^. It is therefore likely that, while the lower number of ^31^P atoms per capsid can account for about a factor of two in the loss of signal, at least part of the lower signal intensity indeed comes from the higher dynamics of the CTD relative to the phosphate backbone of the RNA in the RNA-containing capsids. In addition, when HAP_R10 was added, neither phosphorylation nor bound RNA were detected (**Figure 4**). This could support the interpretation that either the CTD is too mobile to detect phosphorylation in presence of HAP_R10, or, less likely, that RNA remains loosely bound and is thus rather mobile in the presence of CAM-A, not yielding strong signals in the cross-polarization-type transfers used for the ^31^P experiments.

In summary, we observe that capsids assembled in the presence of CAM-E and nucleic acids indeed package the mRNA present in the cell-free synthesis reaction, even while in cells it does not package pgRNA. Capsids assembled in presence of CAM-A on the contrary are fully phosphorylated in this *in-vitro* WG-CFPS set up.

## Discussion

We here present insight into the structural details of aberrant HBV Cp assemblies formed in the presence of CAM-A capsid assembly modulators. We showed that in the resulting assemblies, a single conformation of Cp is present, in contrast to the four distinct, quasi-equivalent Cp monomer conformations in the icosahedral T=4 capsid. We further demonstrate that the NMR fingerprint directly identifies residues involved in CAM binding, allowing to use NMR chemical-shift perturbations as discriminators between CAM-E and CAM-A. In addition, our data reveal that, despite the different appearance on the mesoscopic level as seen in electron micrographs, the assembly domains of full-length Cp183 and P7-Cp183 adopt the same structure as Cp149 on the molecular level when bound to CAM-A. We identified how different CAMs modulate Cp183 phosphorylation and RNA packaging upon Cp assembly directly on exit from the ribosome in a cell-free protein synthesis system.

How do capsids with four different molecules in the asymmetric unit transit to objects with only a single conformation when bound to CAM-A? First, one should mention that, as in Cp capsids, crystalline Ubiquitin^65–68^ and SH3^69^ show peak multiplicity related to the number of molecules in the asymmetric unit. This is thus a feature also observed for other proteins. Still, T=3 icosahedral Nackednavirus^70^ capsids do not show the expected peak multiplicity (manuscript under review), but reveal substantially broader NMR signals than the typically very narrow signals of Ubiquitin, GB1 and the HBV capsid, pointing to small, unresolved peak splitting instead of a clear multiplicity. That Cp assemblies in the presence of CAM-A yield very narrow NMR signals, without any sign of multiplicity indicates that, within the limit of the (narrow) line widths, a single molecule is present in the asymmetric unit of the aberrant assemblies formed in the presence of CAM-A. If the T=4 capsids simply opened to form planar lattices, they would transit into a planar trihexagonal tiling showing three different molecules in the asymmetric unit (B, C and D monomers). Since the spectra however show only a single signal, a single molecule must be present in the asymmetric unit, as in a hexagonal lattice shown in Figure 5b. That the aberrant Cp149 assemblies are not fully planar might be due to the fact that inter-dimer interactions in the presence of CAM-A in WT capsids are not compatible with a fully planar arrangement. The single signals also indicate full occupancy with CAM-A, since no (or only very weak in some preparations) NMR signals corresponding to the unbound state are observed. This is in contrast to cryo-EM studies of Cp150 capsids which revealed binding of one CAM-A per dimer^21,29,32^. The partial occupancy there might be due to the forced maintenance of a closed capsid structure by disulfide crosslinking of the C-terminally added cysteine^28^, while the planar symmetry in the Cp149 Y132A crystals sterically allowed saturation of all sites^34^. One can mention that most residues are more flexible in the presence of CAM-A. This globally increased flexibility points to a less constrained assembly, where notably the spike can make larger excursions.

**Figure 5:**
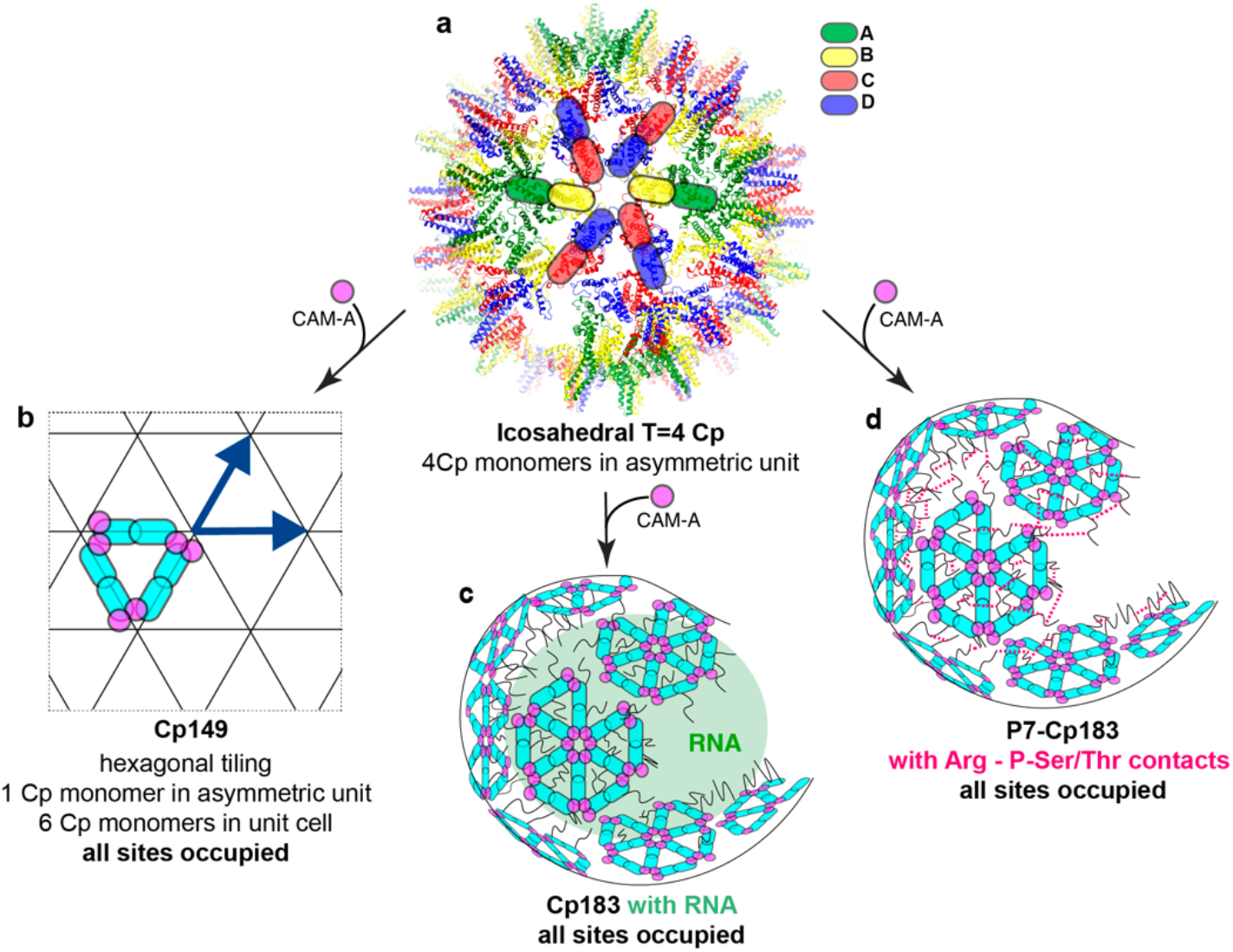
A molecular model for capsid assembly modulation. a) Capsid without CAM. b-c) Models of capsid and dimer rearrangements in the presence of CAM-A. **b**) The hexagonal tiling resulting from capsid opening and loss of the pentameric sites is shown with all molecules colored in cyan; CAM-A are shown in pink. **c**) Effect of CAM-A on Cp183 capsids. All CAM binding sites are occupied, and pentameric axes are abolished. RNA holds the Cp dimers in an approximately round shape. **d**) Effect of CAM-A on P-Cp183 capsids, forming similar superstructures to the capsids in **c**, but with Arg-P-Ser/Thr interactions stabilizing the capsids.

A typical NMR fingerprint is observed for CAM-interacting Cp. The involved amino-acid residues in part coincide with those identified previously^6^, but the NMR chemical-shift analysis interestingly adds the region from residues 14-17, and identifies in greater detail which residues are involved in the region from 124-134.

Most intriguingly, we observe different morphology for Cp149 and Cp183 in the presence of CAM-A on the mesoscopic scale: while the truncated Cp149 capsids open up at much lower CAM-A concentrations, Cp183 capsids retained a virtually globular shape even at high CAM concentrations. This observation becomes central in the context of a recent study in full-length Cp183-expressing Huh7 cells, which investigated a CAM-A, BAY 41–4109^18^. In this cellular context, CAM-A did not induce the irregular assemblies generally seen by EM for Cp149 *in vitro*, but instead dense arrays of what resembled intact capsids. The authors concluded that this might reflect a different mechanism of BAY 41–4109-mediated Cp misassembly under *in-vivo* vs. *in-vitro* conditions^18^. Our data at the mesoscopic level corroborates their findings, but the NMR data demonstrate that the effect of CAM-A on Cp149 and Cp183 is, despite the apparent difference, the same at the molecular level. According to our findings, negative staining electron micrographs used as readout are not resolutive enough to reflect that, while the capsids seem intact, they actually lost the distinctive icosahedral symmetry of intact capsids (**Figure 5c**). The seemingly intact shape/morphology of CAM-A treated Cp183 but not Cp149 capsids at a mesoscopic scale is most likely maintained by intermolecular interactions involving the CTD of Cp183, as sketched in **Figures 5c,d** for Cp183 and P-Cp183 respectively. This is plausibly explained by RNA-CTD interactions in non-phosphorylated capsids, and has also been observed in a previous EM study, where empty Cp183 capsids were destabilized by the synergic action of Importin-β and HAP12, but RNA-filled capsids resisted^42^. However, CTD-RNA interactions cannot account for the high stability of P7-Cp183 capsids which are devoid of RNA. As alternative intermolecular interactions involving the CTDs the positively charged arginine side chains can form salt-bridges with the phosphorylated sidechains from other P7-Cp183 dimers, building a densely woven interaction network which maintains the apparently globular particle shape in a similar way as when interacting with packaged RNA. This view is meanwhile experimentally supported by recent NMR measurements which show that the CTDs of P7-CP183 show a similar dynamic behavior to those of pgRNA-filled Cp183, and must thus be involved in interactions, likely of intermolecular nature, between arginine side chains and phosphorylated Ser/Thr residues^57^.

Intriguingly, the apparent stability of Cp183 particles against CAM-A was reduced upon addition of DTT. Current evidence points to a role for Cys107 and, to a lesser extent, Cys61 although the underlying mechanism is as yet unclear. Still, these observations might be of importance for investigations on co-factors of CAM-A-induced capsid destabilization, as recently reported for importin-β^42^; the presence of DTT in such experiments might induce unexpected effects.

Capsid assembly modulation directly upon exit of Cp183 from the ribosome in a cell-free protein synthesis system revealed how CAMs impact Cp183 capsid assembly in situ and in the presence of RNA. Interestingly, RNA packaging was distinctly affected compared to reported CAM activities in live mammalian cells. There, CAM-E compounds interfere with pgRNA encapsidation^17,64,71,72^, whereas the ^31^P NMR spectra in our work clearly revealed packaged mRNA in capsids from cell-free synthesized Cp183 in the presence of CAM-E. It is indeed difficult to apprehend how CAMs could actually inhibit packaging of RNA on capsid formation, since charge equilibration seems an essential driving force in capsid assembly, and the high concentration of positively charged arginine residues in the CTD makes this process in the absence of RNA difficult under physiological conditions. The empty capsids described to form in cells in the presence of CAM-E must thus either be phosphorylated to a large extent, or they contain nonspecifically packaged cellular RNA, which would typically not have been detected in the corresponding studies, where pgRNA sequence-specific hybridization probes were used^64,71^.

In contrast, in CAM-A treated cell-free synthesized Cp183 samples did not show any signal for Cp183-associated RNA, conceivably because the resulting flat assemblies (due to the presence of DTT) cannot act as protective RNA containers, in contrast to the preformed E. coli capsids treated with CAM-A, which maintain more than 50 % of the packaged RNA (**Figure S12**). Interestingly, however, cell-free synthesized Cp183 assembled in presence of JNJ-890 gave clear signals of protein phosphorylation, supported by the presence of adequate kinases in the wheat-germ extract used for cell-free protein synthesis^62^, whereas no phosphorylation was seen in HAP_R10-treated and neither in untreated Cp183 samples.

## Conclusion

We here used solid-state NMR to show that the molecular level organization of the aberrant HBV Cp assemblies induced by modulation with CAM-A compounds changes from T=4 icosahedral to hexagonal. The fact that the mesoscopically irregular assemblies actually show high order on the molecular level, as reflected in the narrow lines in the NMR spectra, was surprising, and shall further motivate NMR investigations of systems which have a mesoscopically heterogenous appearance. We showed that NMR fingerprints can clearly distinguish between CAM-A and CAM-E type of action, even for full-length Cp183 capsids where EM images can be misleading. Importantly, our data reconcile *in vitro* and *in vivo* views on CAM-A mode of action. Our analyses also reveal an influence of DTT on aberrant capsid formation, but the underlying principles remain not fully understood. Using a combination of advanced synthetic biology and NMR techniques, we also elucidated how CAMs affect phosphorylation and RNA packaging *in vitro* during Cp183 capsid assembly, challenging previously established interpretations of empty capsid formation by CAM-E.

In summary we showed that NMR fills important gaps in the understanding of the molecular effects of notably CAM-A, and by this contribute to the further development of capsid assembly modulation.

## Acknowledgements

This work was supported by the CNRS (L.L. CNRS-Momentum 2018), the French Agence Nationale de Recherches sur le Sida et les hépatites virales (A.B. ANRS, ECTZ71388 & ECTZ100488), and a PhD grant from the Chinese Scientific Council to SW. Financial support from the IR-RMN-THC Fr3050 CNRS for conducting the research is gratefully acknowledged. B.H.M. was supported by an ERC Advanced Grant (grant number 741863, Faster), and by the Swiss National Science Foundation (200020_188711). We thank the Centre d’Imagerie Quantitative Lyon-Est (CIQLE) for support at the EM platform.

## Author contributions

LL recorded ^13^C-detected and ^1^H-detected NMR spectra and analyzed NMR data, together with LB. RH, SW and MD prepared capsid samples from *E. coli* expression. LB and SW, together with MLF, prepared cell-free synthesized capsid samples. RH, LB and MB took EM micrographs. TW recorded ^31^P spectra. AM and MC recorded the relaxation data and did the dynamics analysis. AB and MN, together with LL, BHM, and DD designed the study. DB and JMB selected and provided the CAM compounds. AB, MN, BHM, LL and MLF analyzed data and wrote the manuscript. All authors read and approved the final version of the manuscript.

