## Supplementary Data for "Molecular elucidation of drug-induced abnormal assemblies of the Hepatitis B Virus capsid protein by solid-state NMR"

### Drug-induced aberrant hepatitis B virus capsids form superstructures with a single molecule in the asymmetric unit

#### Material and Methods

##### *Capsid assembly modulators (CAMs)*

GS-832471 and GS-942049 are from Gilead. JNJ-632, JNJ-827 and JNJ-890 are from Johnson & Johnson. GS-837886 (called HAP_R10 here) was a kind gift from Roche. All compounds were dissolved in 100 % DMSO to a final concentration of 10 mM and stored at -20 °C.

##### *Cp from bacterial expression and incubation with CAMs*

For EM studies, all Cp samples (Cp140, Cp149, Cp183, P7-Cp183 and Cp183 C183A mutant) were expressed in *E. coli* using LB medium and purified on a sucrose gradient as described in detail in [1]. Samples were dialyzed in 50 mM HEPES buffer at pH 7.5 and kept at 4 °C.

For solid-state NMR studies, Cp149, Cp183 and P7-Cp183 were expressed in *E. coli* using M9 medium supplied with $^{15}NH_4Cl$ and $^{13}C$-glucose, and purified on a sucrose gradient.

Cp capsid samples were incubated at a concentration of ~1 mg/ml (corresponding to ~60 µM monomer concentration for truncated Cp140 and Cp149 and ~46 µM for full length Cp183 and P7-Cp183) with relevant monomer:CAM ratio for 2 h at 37 °C in 50 mM HEPES buffer at pH 7.5, in presence of 5 mM DTT when specified.

##### *Preparation of solid-state NMR samples produced in E. coli*

$^{13}C$-$^{15}N$-labeled Cp183 and P7-Cp183 capsids produced in *E. coli* were dialyzed in 50 mM TRIS at pH 7.5 and loaded onto a HiPrep™ 16/60 Sephacryl® S-200 HR column (120 ml) to remove Triton-X100 which was shown to bind in the hydrophobic pocket of Cp and induces large CSPs in the NMR spectra[1], except for the samples of Cp183 with JNJ-632 and JNJ-890. For each sample, between 15 and 20 mg of autoassembled capsids at a concentration ~1-1.5 mg/ml were incubated at a monomer:CAM ratio of 1:4 for 2 h at 37 °C.

In the case of Cp149, NMR samples were prepared starting from Cp dimer (see **Table S1**). For this, $^{13}C$-$^{15}N$-labeled and $^{2}H$-$^{13}C$-$^{15}N$-labeled Cp149 capsids were disassembled as described in [1]. Freshly separated dimers after gel filtration were dialyzed in 50 mM HEPES buffer at pH 7.5, 5 mM DTT. Concentration was determined using a nanodrop measurement, adjusted to ~1 mg/ml (~30 µM dimer concentration), and dimers were incubated with a monomer:CAM ratio of 1:4 for 24 h at room temperature (RT) in presence of 150 mM NaCl. Reassembled capsids samples were diluted by 2 in 50 mM HEPES buffer at pH 7.5 to reduce the final NaCl concentration to 75 mM, since high salt concentrations may reduce the NMR sensitivity.

For all NMR samples, final DMSO content always stayed below 3 %, and EM grids were prepared at the end of the incubation step. Samples were concentrated to 1 ml final volume and

sedimented into 3.2 mm rotors by an overnight ultracentrifugation at 200,000 *g*, 4 °C. Excess of sediment was removed and 1.5 µl of saturated DSS (0.25 M) were added before closing the rotor for NMR chemical-shifts referencing. In addition, $^{2}$H-$^{13}$C-$^{15}$N-labeled Cp149 capsids in presence of JNJ-890 were filled in 1.3 mm and 0.7 mm rotors for $^{1}$H-detection NMR experiments and comparison with cell-free samples.
A summary of all the NMR samples prepared is shown in **Table S1**.

***Cp from wheat-germ cell-free protein synthesis***

Cp proteins have been produced in the wheat germ cell-free system using the dialysis method, also called CECF for continuous exchange cell-free system[2,3]. Small-scale experiments have been performed in CECF-mini-reactors manufactured at ETH Zurich after models from reference[4], while medium-scale and large-scale experiments have been performed in 500-µl and 3-ml dialysis cassettes, respectively[5,6]. In all cases, transcription and translation have been performed separately. In mini-reactors, the translation mixture contained ½ volume of home-made wheat-germ extract (WGE), ½ volume of mRNA, 40 ng/µl of creatine kinase and 6 mM of amino acid mix for a total volume of 70 µl, while in dialysis cassettes, it was composed of ½ volume of feeding buffer, 1/3 volume of mRNA, 40 ng/µl of creatine kinase, 0.3 mM of amino acid mix, and 1/6 volume of home-made WGE. The feeding buffer (1.5 ml for a mini-reactor, and 20 ml or 120 ml for a 500-µl or a 3-ml dialysis cassette, respectively) contained 30 mM HEPES-KOH pH 7.6, 100 mM potassium acetate, 2.7 mM magnesium acetate, 16 mM creatine phosphate, 0.4 mM spermidine, 1.2 mM ATP, 0.25 mM GTP and 4 mM DTT (SUB-AMIX NA, CellFree Sciences), supplemented with 6 mM of amino acid mix. A mix containing all twenty isotopically labeled amino acids (Cambridge Isotope Laboratory) was used for the production of $^{2}$H-$^{13}$C-$^{15}$N-Cp183 in a 3 ml-translation reaction experiment for NMR studies. A dialysis membrane with a molecular weight cut-off of 10 kDa has been used, and protein synthesis has been performed for 16 h at 22 °C under agitation. CAMs have been added both to the translation mix and to the feeding buffer at different molar ratios, considering the Cp concentration to be 0.25 mg/ml WGE. The total cell-free reaction mixture for CF-Cp183 and CF-Cp183+JNJ-632 was treated with 25,000 units/ml of benzonase for 30 min at room temperature before centrifugation at 20,000 *g*, 4 °C for 30 min. The supernatant (SN) was loaded onto a discontinuous sucrose gradient with layers of 10 to 60 % as described in [7], and 50-60% fractions were used for 1.3 mm NMR rotor filling by overnight ultracentrifugation (200,000 *g*, 4°C). No benzonase treatment was done when CAMs-A were added to the reaction, and the pellet fractions were used for 1.3 mm NMR rotor filling.

##### *Negative staining electron microscopy*

For EM grids preparation, 5 µl of sample were adsorbed to the surface of carbon-coated grid for 2 min, sample excess was removed with paper, and grids were stained with 2% uranyl acetate for 2 min. Samples were imaged with a JEM-1400 transmission electron microscope operating at 100 kV.

##### *NMR experiments for chemical-shift analysis and dynamics measurements*

NMR experiments were conducted using a 1.3 mm and a 3.2 mm triple-resonance ($^{1}H$, $^{13}C$, $^{15}N$) probe heads at static magnetic field of 18.8 T corresponding to 800 MHz proton resonance frequency (Bruker Avance III). Assignments of unbound Cp were taken from references [8] (BMRB number 27317) and [1] (BMRB number 28122). Cp-CAM-bound resonances were assigned using a combination of 2D and 3D correlation experiments including: 2D DARR, 2D NCA, 3D NCACX, and when necessary a 3D CANCO[9,10]. Carbon-detected experiments were recorded at a MAS frequency of 17.5 kHz on all samples and are detailed in **Table S2**. Proton-detected experiments were recorded at a MAS frequency between 55 and 60 kHz and are detailed in **Table S3**. All spectra were referenced using DSS and recorded at a sample temperature of 4 °C in the 3.2 mm probe and between 20 and 25 °C in the 1.3 mm probe, as determined by the resonance frequency of the supernatant water[11].

For dynamics measurements, site-specific solid-state NMR relaxation rate constants $R_{1\rho}(^{15}N)$ were measured on $^{2}H$-$^{13}C$-$^{15}N$ Cp149 alone and in interaction with JNJ-890, using the 3D hCANH sequence described in[12] in an 0.7 mm rotor at 80 kHz MAS, 20 T (850 MHz) external magnetic field strength and with a 13 kHz spin-lock strength. For further experimental information see **Table S4**. Site-specific relaxation-rate constants were obtained within MATLAB using the spectral fitting package INFOS[13]. Relaxation curves were fitted with a mono-exponential fit with two degrees of freedom ($A \cdot \exp(-Rt)$). Error bars thereof were obtained using bootstrapping methods.

$^{31}P$,$^{1}H$ cross-polarization experiments were recorded at 11.7 T (500 MHz proton frequency) in a triple-resonance 1.3 mm probe. The MAS frequency was set to 60 kHz and the sample temperature was in-between 20 to 25 °C. The CP experiments were optimized on solid $NH_4H_2PO_4$ and typical radio-frequency field strengths used in the CP were 100 kHz for $^{1}H$ and 40 kHz for $^{31}P$. The CP contact time was set to 1 ms and the repetition time to 1.5 s. The acquisition time was set to 10.2 ms. WALTZ-64 $^{1}H$ decoupling (5 kHz rf-field strength) was applied during acquisition. The number of scans were 52240 (CF-Cp183 and CF-Cp183 + JNJ-632), 41984 (CF-Cp183 + JNJ-890) and 58408 (CF-Cp183 + HAP_R10). Note that the spectrum of CF-Cp183 was recorded at 55 kHz MAS and thus slightly different CP conditions. All spectra were referenced to $H_3PO_4$.

Processing of all spectra was done using TopSpin 3.2 (Bruker Biospin) with zero filling and a squared cosine apodization function (SSB 2.5–3 depending on spectrum and dimension). Spectra analysis and assignment was done with the CcpNmr Analysis package[14,15]. Chemical-shift perturbations induced by CAMs were calculated for all $^{13}C$ and $^{15}N$ assigned nuclei and plotted using a home-made python script. For $^{15}N$ nucleus, the CSPs were calculated by taking into account the gyromagnetic ratio compared to the carbon in order to be comparable:

$$\Delta\delta_N = \left(\frac{\gamma_N}{\gamma_C}\right) |\delta_N[bound] - \delta_N[unbound]| = 0.4 * |\Delta\delta_N|$$

For Cp183 capsids produced in the WG-CFS, CSPs were calculated for $^{1}H$ and $^{15}N$ nuclei using the following formula:

$$\Delta\delta_{HN} = \sqrt{(\Delta\delta_H)^2 + \left(\frac{\gamma_N}{\gamma_H}\Delta\delta_N\right)^2}$$

The figures involving structures were all prepared using the PyMol software (Warren Delano, http://www.pymol.org).

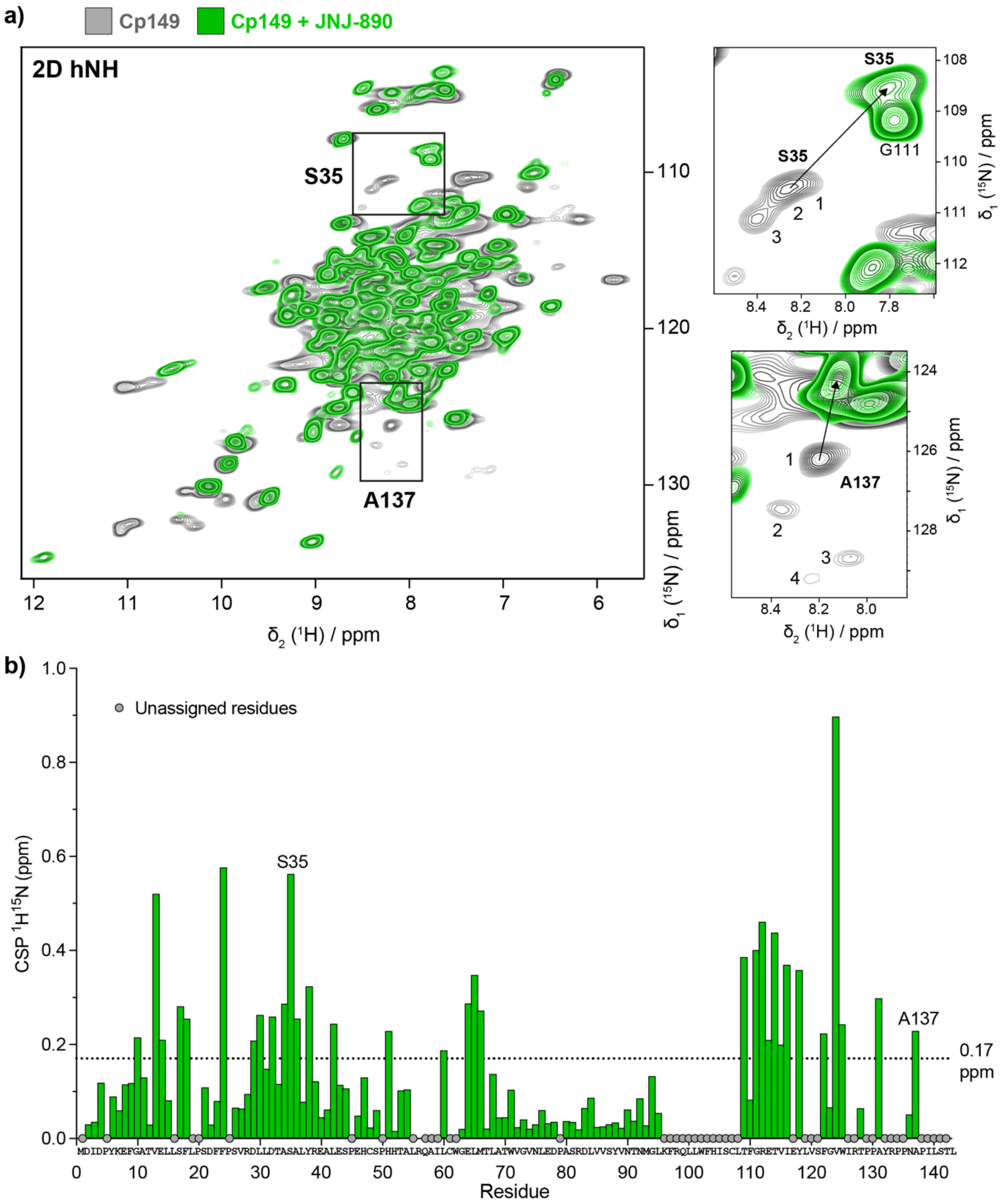


**Figure S1**: **Effect of JNJ-890 on Cp149 capsids at the molecular level. a)** Overlay of 2D hNH spectra of $^{2}$H-$^{13}$C-$^{15}$N-Cp149 in absence (grey) and in presence of JNJ-890 (green) and extracts showing the A137 alanine and S35 regions. In both types of spectra, the peak splitting phenomenon due to subunits asymmetry disappears in presence of CAM-A. Spectra were recorded at 60 kHz MAS on a 800 MHz spectrometer. Corresponding EM micrographs are shown in **Figure 1c**. **b)** Chemical shift perturbations (CSPs) induced by the binding of JNJ-890 and opening of Cp149 capsids. The average CSP for all assigned residues is 0.17 ppm.

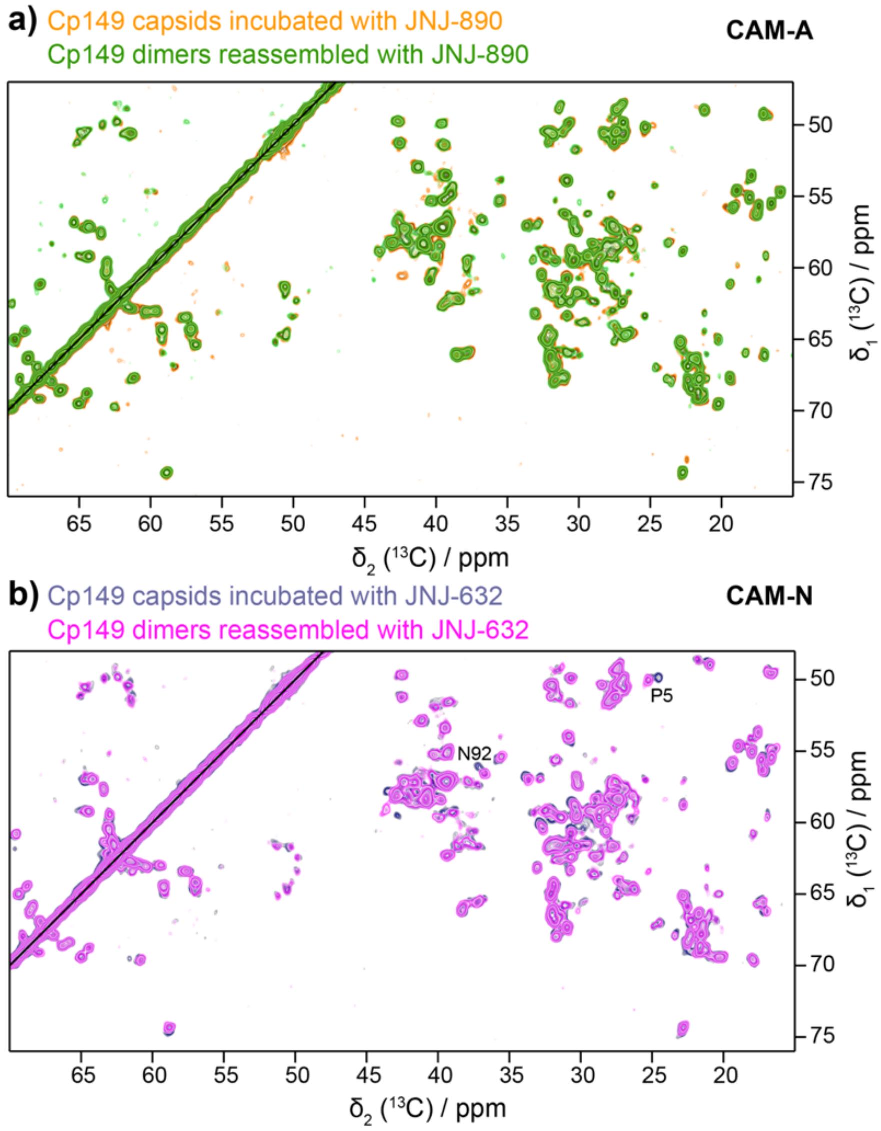


**Figure S2: CAMs have a similar effect on Cp149 dimers and preformed Cp149 capsids.** Zoom on aliphatic region of a 2D DARR of **a)** Cp149 capsid incubated with JNJ-890 for 2 hours at 37 °C (orange) and Cp149 dimer assembled with JNJ-890 overnight at room temperature (green); and **b)** Cp149 capsid incubated with JNJ-632 for 2 hours at 37 °C (grey) and Cp149 dimer reassembled with JNJ-632 overnight at room temperature (pink). The few differences observed in panel b) are due to the bound Triton X-100 in the hydrophobic pocket[1], which was not removed in this sample.

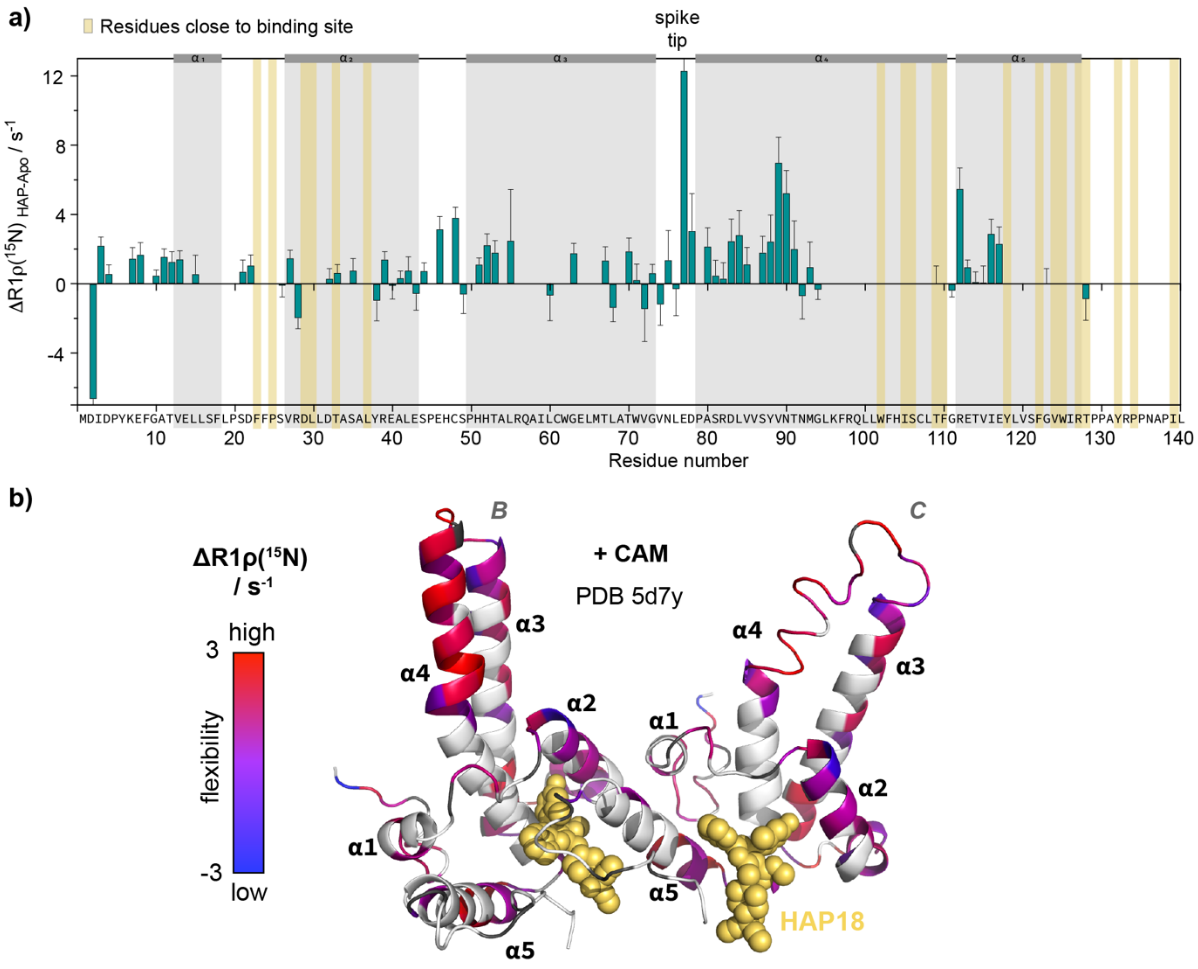


**Figure S3:** a) Differences of $R_{1\rho}(^{15}N)$ rate constants measured at 80 kHz MAS and 13 kJz spin-lock field plotted for each residue upon JNJ-890 binding. **b)** $R_{1\rho}$ relaxation parameter mapped on the Cp149 structures for Cp149 (left, PDB 1qgt[16]) and for Cp149 with JNJ-890 (PDB 5d7y[17] with HAP18 molecule shown in golden spheres). The color scale goes from red (relatively flexible) to blue (more rigid). Error bars have been calculated by Gaussian error propagation.

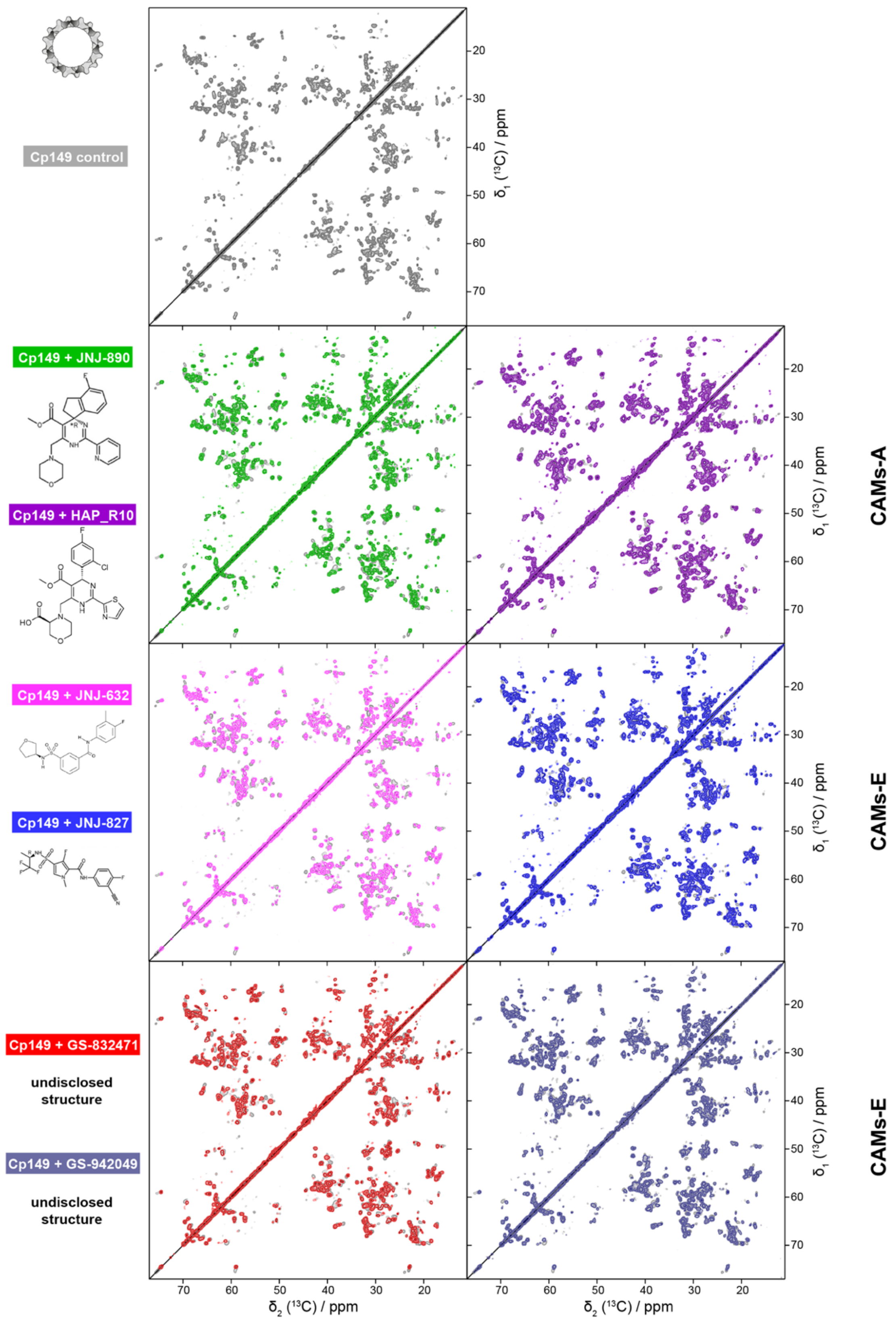


**Figure S4:** NMR DARR spectra of all samples of $^{13}C$-$^{15}N$ Cp149 dimer reassembled in absence (grey) and in presence of different CAM: JNJ-890 (green), HAP_R10 (GS-837886) (purple), JNJ-632 (pink), JNJ-827 (blue), GS-832471 (red) and GS-942049 (steel). The control is shown behind each spectrum in light grey. Corresponding EM pictures are shown in **Figure 2** and chemical shift perturbation in **Figure S5**.

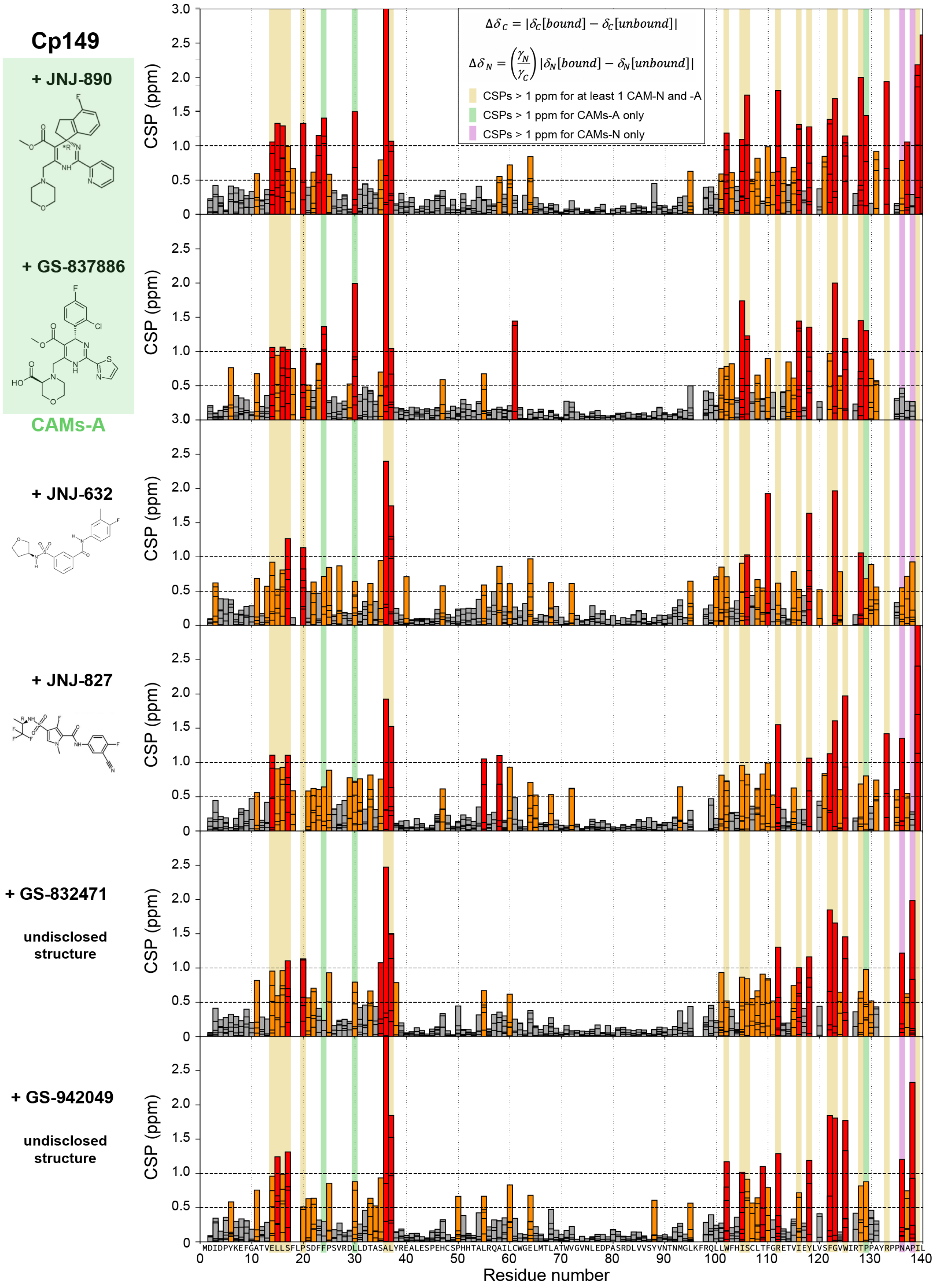


**Figure S5a. Chemical shift perturbations induced by different CAMs on Cp149 capsids.** CSPs were calculated for all assigned $^{15}$N and $^{13}$C nuclei using 2D and 3D spectra compared to the control sample without CAM. For $^{15}$N CSPs, a factor of 0.4 was applied corresponding to the gyromagnetic ratio of nitrogen over carbon. Residues with medium ($0.5<\text{CSP}<1$ ppm) and large (>1 ppm) CSPs are colored in orange and red, respectively. CSPs mapped on the capsid structure are shown in **Figure 2**. Residues the most affected by both CAMs-A and CAMs-E are highlighted in yellow, only by CAMs-A in green and only by CAMs-A in purple.

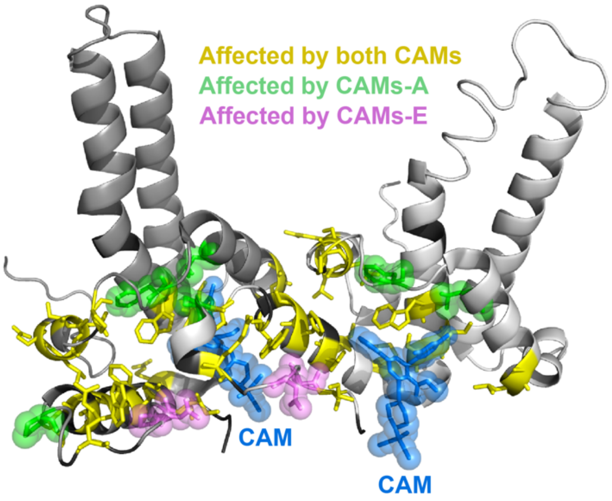


**Figure S5b. CSPs induced by CAM-E and CAM-A compounds mapped on Cp structure.** The structure highlights in yellow the residues which show CSPs for both CAM-E and CAM-A, and thus define the binding atoms. These are indeed rather similar between CAM-A and CAM-E. In green are highlighted residues that show only CSPs in CAM-A, pointing to residues sensitive to the change in lattice organization. The presence of these thus point to a CAM-A. The most straightforward way to distinguish by NMR CAM-E and CAM-A is however through the collapse of the up to fourfold peak multiples which are observed for T=4 icosahedral capsids. PDB used: 5d7y[17].

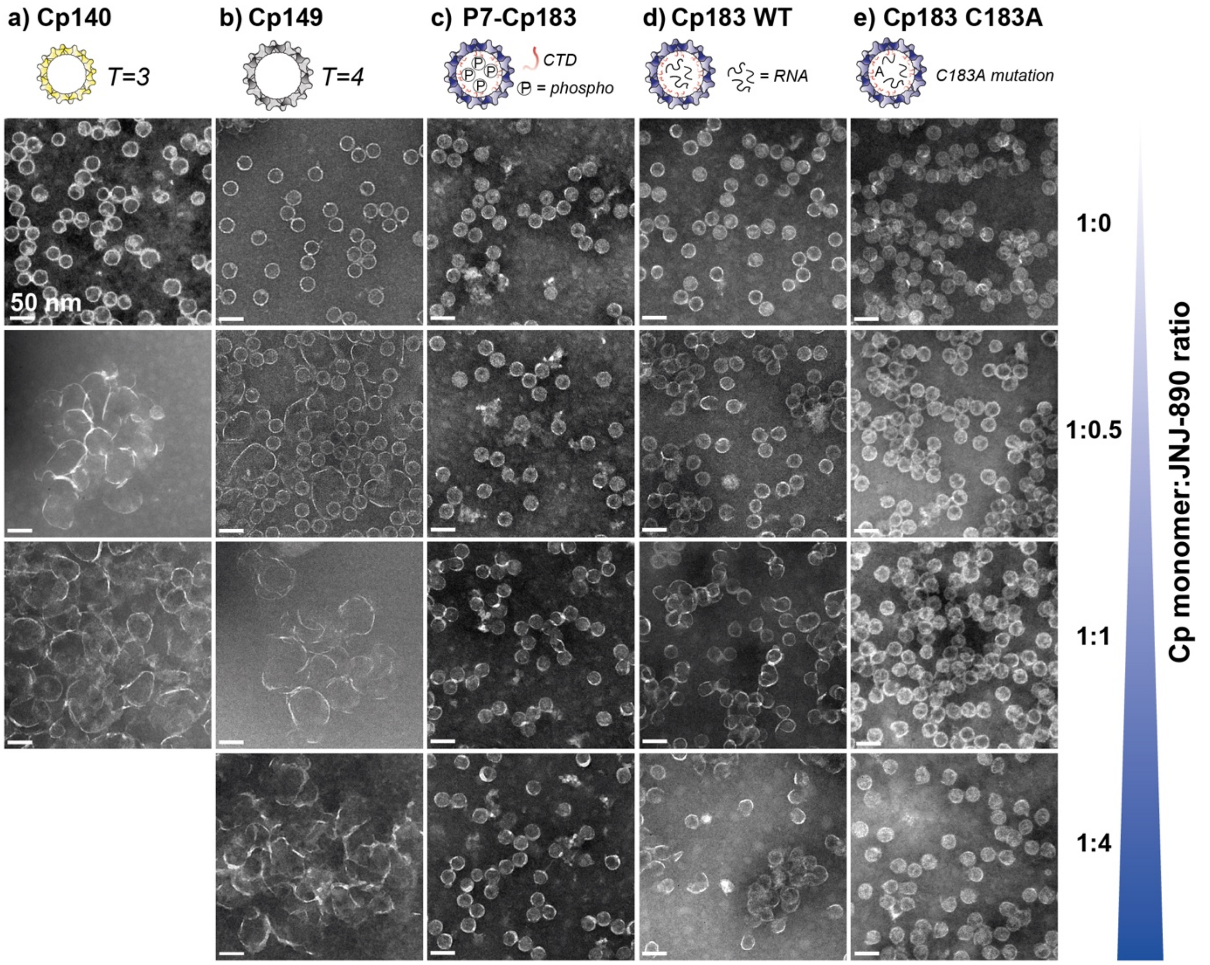


**Figure S6: Impact of different JNJ-890 ratios on capsids.** CAM-A at molar ratios from 0 to 4 molar equivalents were incubated with **a)** Cp140, **b)** Cp149, **c)** P7-Cp183 WT, **d)** Cp183 WT and **e)** Cp183 C183A mutant. Scale bars = 50 nm.

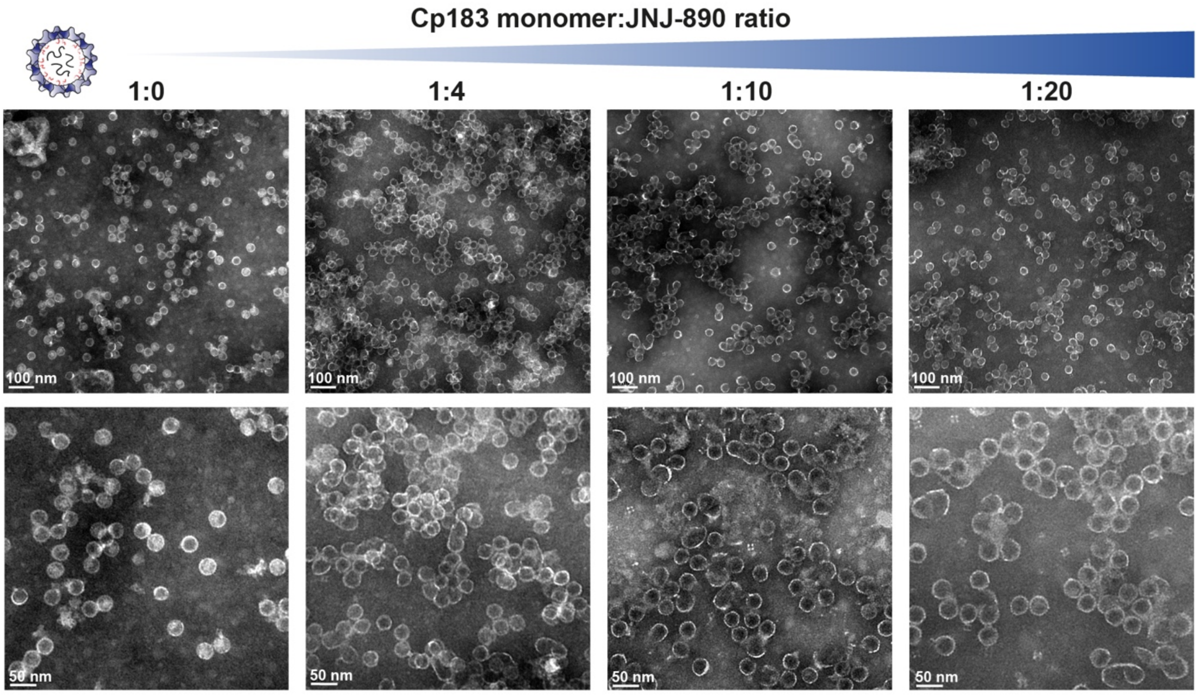


**Figure S7: Impact of increasing ratios of JNJ-890 on wild-type Cp183**. EM micrographs of Cp183 capsids control and incubated with 4, 10 and 20 molar equivalents of JNJ-890. Zooms are given for the four conditions. Even at high molar ratio (1:10 and 1:20 monomer:CAM-A), most capsids remain virtually closed.

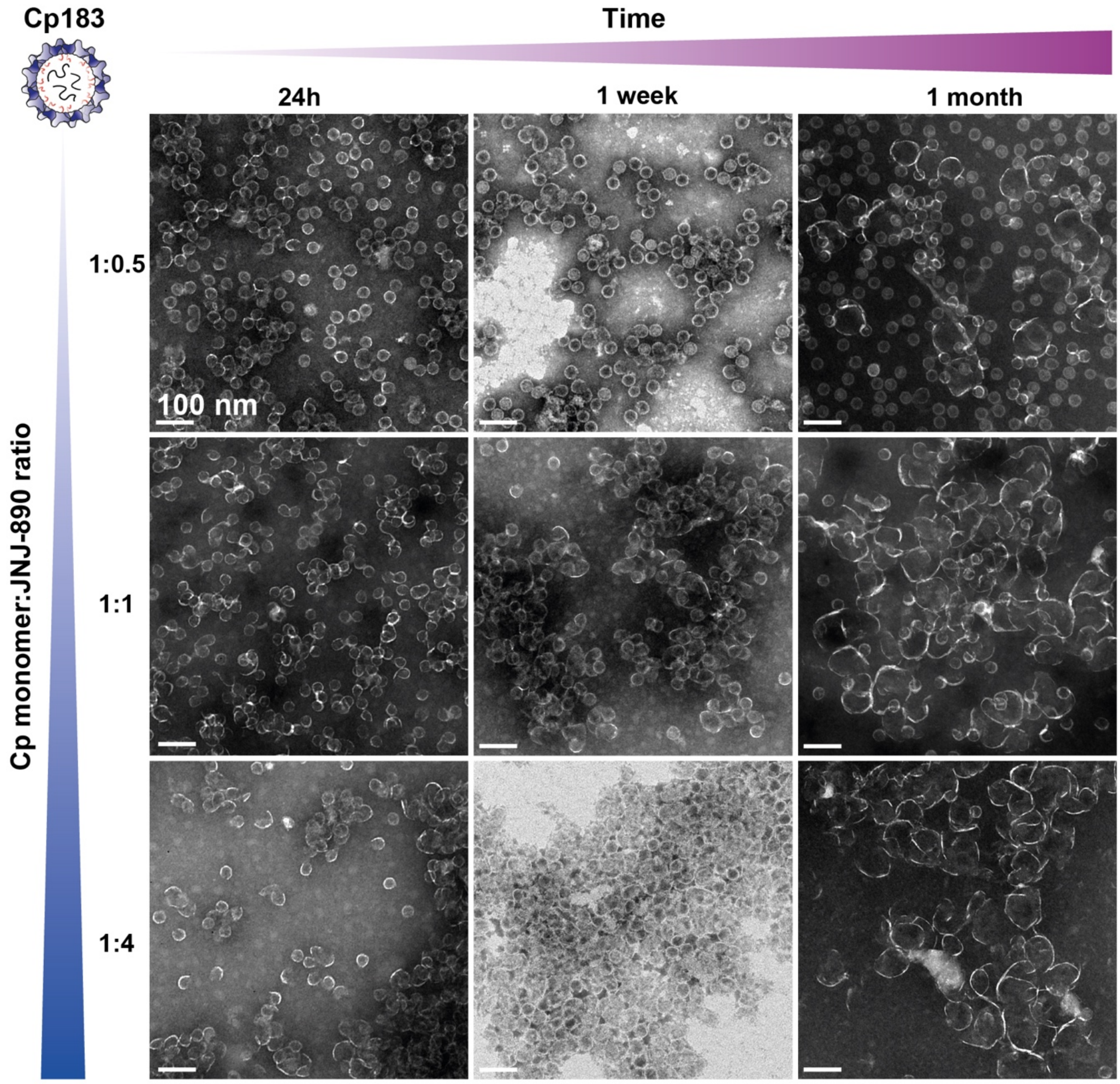


**Figure S8: Impact of time on capsid's opening with different monomer:JNJ-890 ratio**. The effect of CAM-A starts to be visible after 1 week, yet capsids are not all opened even after 1 month at the highest ratio. Scale bar = 100 nm.

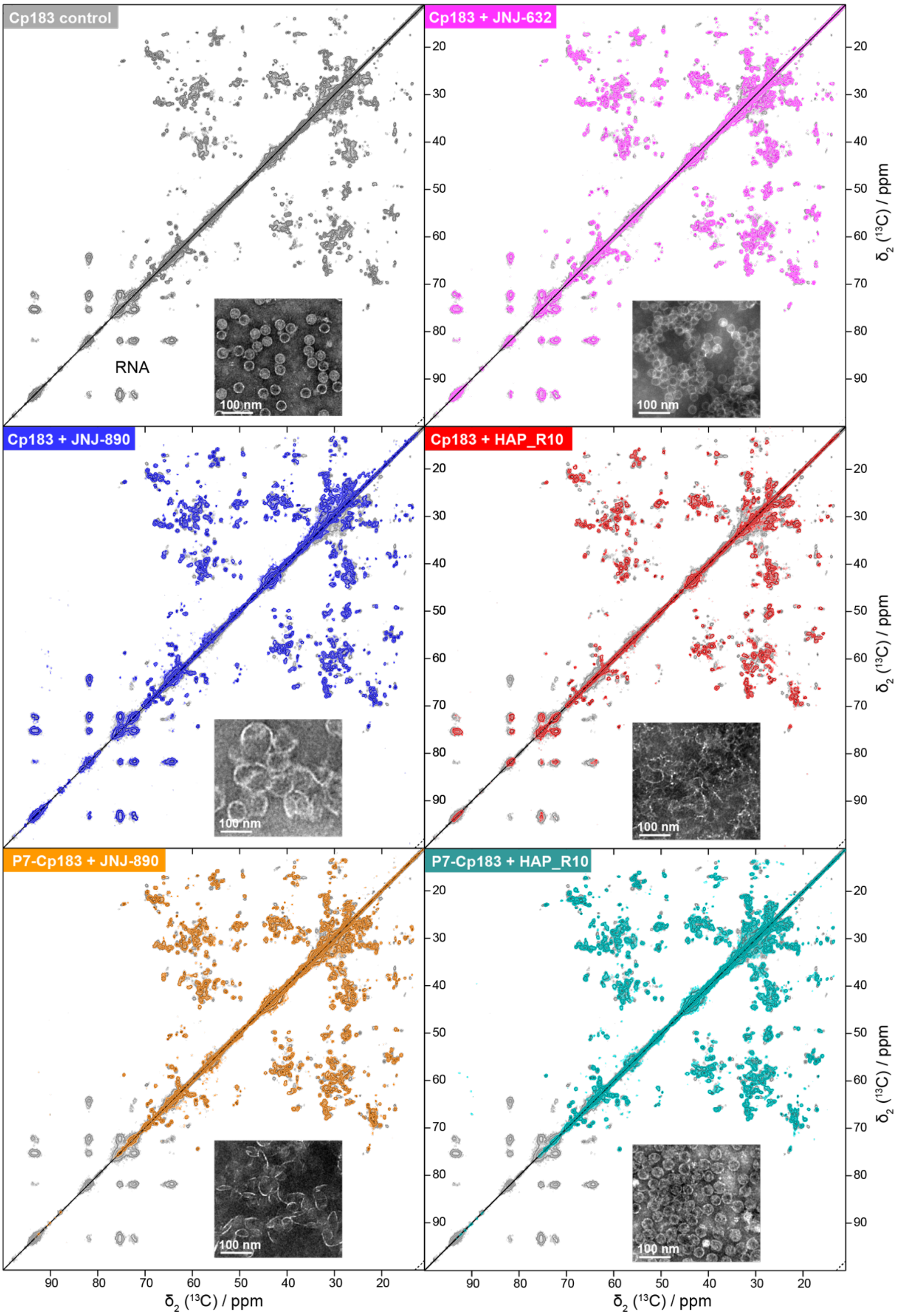


**Figure S9: NMR aliphatic regions of DARR spectra and EM micrographs of Cp183 capsids.** Absence of CAM (grey); $^{13}C$-$^{15}N$ Cp183 capsids incubated with JNJ-632 (pink), JNJ-890 (blue) and HAP_R10 (red), $^{13}C$-$^{15}N$ P7-Cp183 capsids incubated with JNJ-890 (orange) and HAP_R10 (cyan). All capsids are in presence of 5 mM DTT.

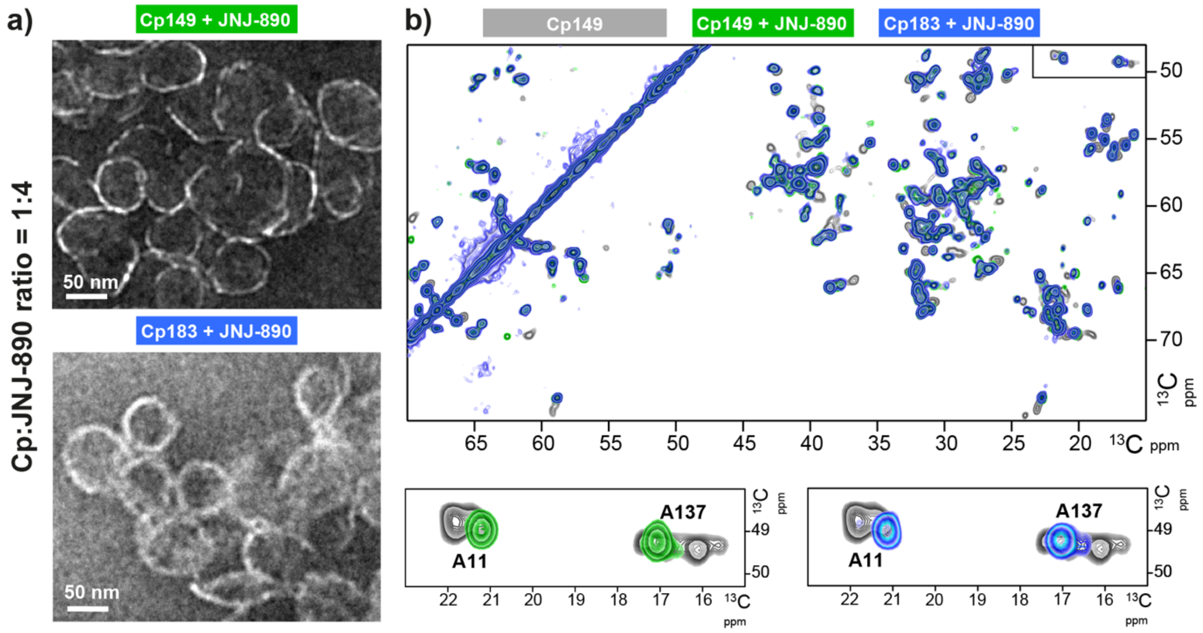


**Figure S10**: **Impact of JNJ-890 HAP on Cp149 and Cp183 capsids. a)** EM micrograph of Cp149 dimer reassembled with JNJ-890 (green) and of Cp183 preformed capsid incubated with JNJ-890 (blue). **b)** Region of 2D DARR spectra of Cp149 without CAM (in grey), Cp149 reassembled with JNJ-890 (green) and Cp183 capsid with JNJ-890 (blue). Extracts from A11 and A137 region are shown below.

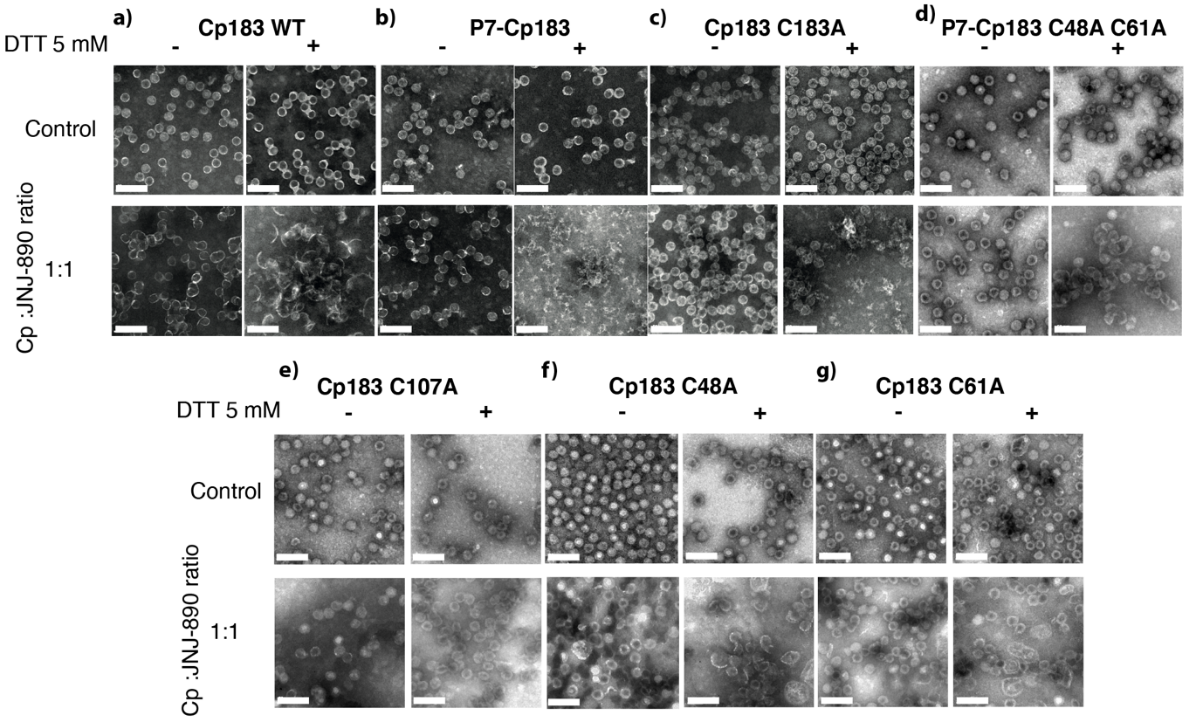


**Figure S11: Impact of DTT on addition of 1 molar equivalent of JNJ-890**. Scale bars 100 nm. It can be seen that for all forms, but C107A and C61A, DTT enhances Cp opening.

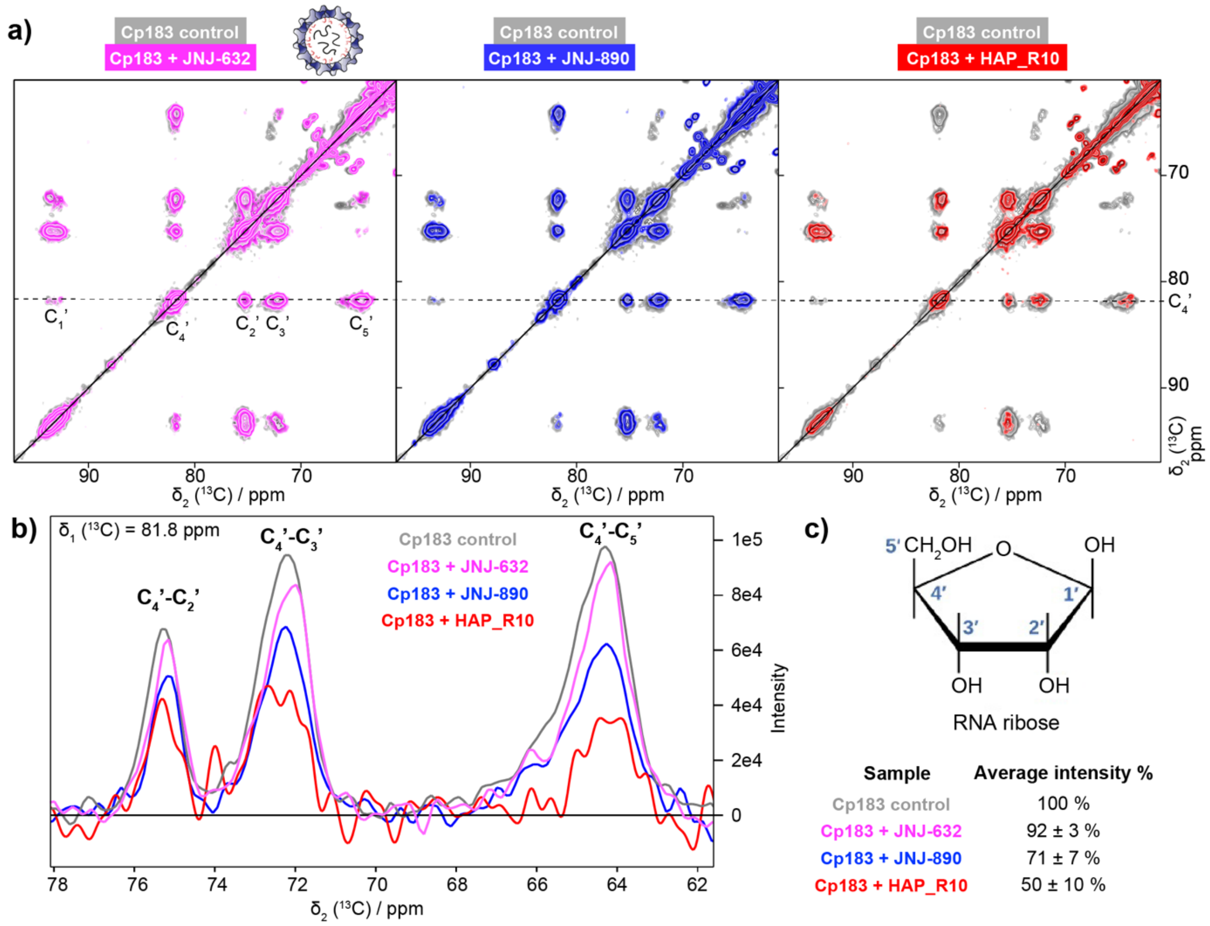


| Sample | Average intensity % |
|---|---|
| Cp183 control | 100 % |
| Cp183 + JNJ-632 | 92 ± 3 % |
| Cp183 + JNJ-890 | 71 ± 7 % |
| Cp183 + HAP_R10 | 50 ± 10 % |

**Figure S12: CAM-A leads to loss of RNA on Cp183 preformed capsids. a)** Region from DARR spectra corresponding to nucleic acids overlayed for Cp183 apo capsids (grey) and Cp183 capsids incubated at a 1:4 ratio Cp:CAM with JNJ-632 (pink), JNJ-890 (blue) and HAP_R10 (red). Corresponding EM micrographs are shown in **Figure S9**. **b)** 1D trace of the three major nucleic acids correlation signals extracted at 81.8 ppm (corresponding to $C_4$' carbon in ribose). Spectra intensities were calibrated on the protein signals. **c)** Representation of RNA ribose and table summarizing the average intensity of the CAMs-bound samples compared to Cp183 sample in absence of CAMs for the three RNA signals. The capsids investigated were purified including by a sucrose gradient (see Material and Methods section), and therefore carry no free RNA before CAM addition. Thus, the RNA observed in the reference and JNJ-632 capsids is the one from inside the capsid. After capsid opening by CAM-A, the capsids are directly ultracentrifuged into the rotor. During this step, free RNA would likely remain in the supernatant, or be too flexible to be observed in these spectra. The decay in signal intensity thus points to partial release of the packaged RNA into solution, while a part remains associated with Cp, but is not enclosed (protected) anymore. Since no nucleases are present in the solution, they are however not degraded.

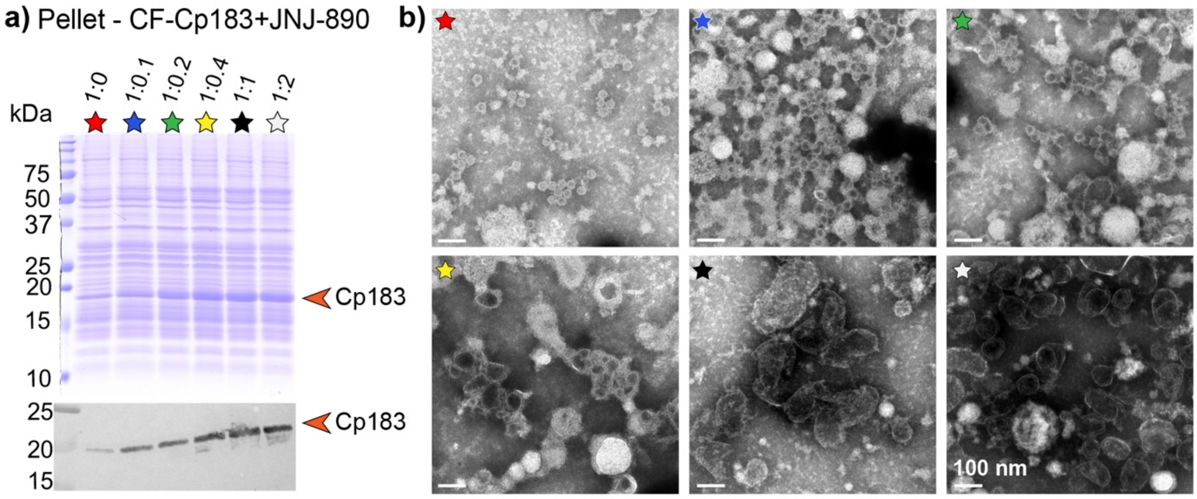


**Figure S13: Capsid solubility and morphology of CF-Cp183 capsids synthesized in the presence of JNJ-890.** **a)** SDS-PAGE and Western-Blot analyses of CF-Cp183 with increasing molar ratios of JNJ-890 (pellet fraction). While in absence of JNJ-890 (red star), most Cp183 is soluble, after addition of CAM-A capsids are mostly found in the pellet. **b)** Negative-staining EM micrographs corresponding to the crude cell-free synthesis reactions, before centrifugation. Capsid's sizes start to increase from the ratio monomer:CAM-A 1:0.1, and capsids are almost fully open from a ratio 1:0.4 (yellow star).

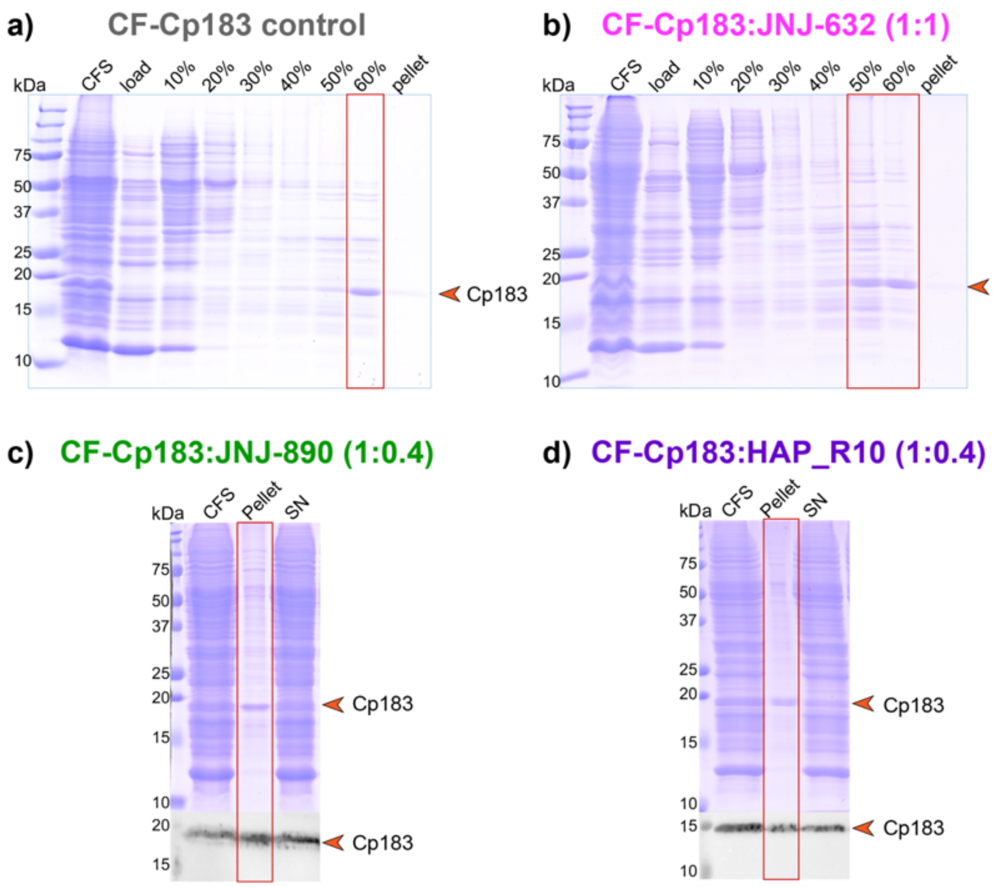


**Figure S14: $^{2}H$-$^{13}C$-$^{15}N$ CF-Cp183 synthesis in absence or presence of different CAMs.** SDS-PAGE of CF-Cp183 purification in presence of **a)** no CAM, **b)** 1 equivalent of JNJ-632, **c)** 0.4 equivalent of JNJ-890 and **d)** 0.4 equivalent of HAP_R10. For the control without CAM and the sample with CAM-E (JNJ-632), the capsids are mainly soluble and migrate on a sucrose gradient, while for both CAMs-A (JNJ-890 and HAP_R10), a large part of the protein sample is found in the pellet after centrifugation and can be directly sedimented into the NMR rotors. The fractions used to fill the 1.3 mm rotors are framed in red. Western-Blots are shown in panel c) and d). CFS: total cell-free sample; 10-60 %: sucrose gradient fractions; SN: supernatant. Corresponding EM pictures and hNH solid-state NMR spectra are shown in the main text in **Figure 4**.

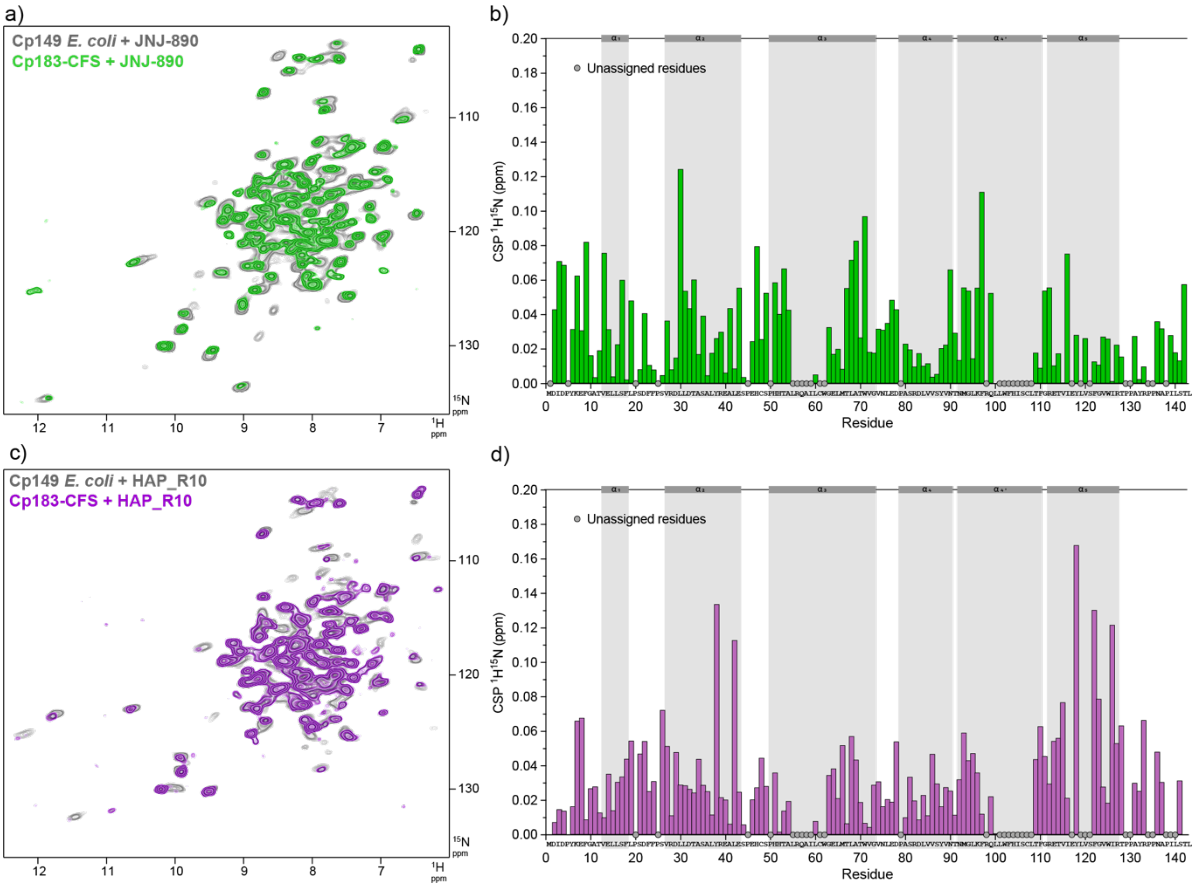


**Figure S15: CSPs cell-free versus *E. coli* capsids in presence of CAMs-A. a)** Overlay of 2D hNH spectra of Cp149 dimer produced in *E. coli* reassembled with JNJ-890 (grey) and CFS-Cp183 produced in presence of JNJ-890 (green). **b)** HN-CSP graph showing the differences between the two spectra. **c)** Overlay of 2D hNH spectra of Cp149 dimer produced in *E. coli* reassembled with HAP_R10 (grey) and CFS-Cp183 produced in presence of HAP_R10 (green). **d)** HN-CSP graph showing the differences between the two spectra, which are consistently small.

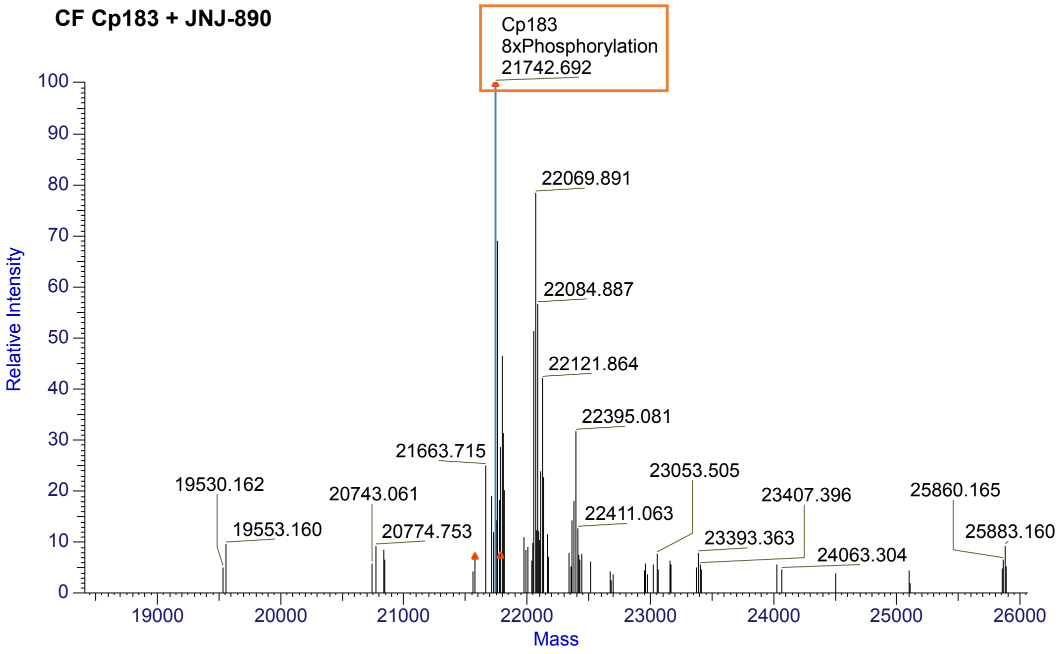


**Figure S16:** Mass spectrometry of unlabeled Cp183 produced in WG-CFPS in presence of 0.4 equivalents of JNJ-890. A mass of 21742 Da is observed, which corresponds to the presence of 8 phosphorylations (expected mass without phosphorylation = 21116 Da).

**Table S1:** List of NMR samples used and experiments recorded. Colors used in the first column correspond to the colors used for the NMR spectra. All samples were devoid of Triton X-100 detergent[1] except Cp149 capsids + JNJ-632 as well as Cp183 with JNJ-632 and JNJ-890, for which the gel filtration step was not performed. NMR experimental details are described in Tables S2 to S4.

| Sample Name | Labeling | NMR experiments | System | Incubation | Figures |
|---|---|---|---|---|---|
| Cp149 control | $^{2}H$-$^{13}C$-$^{15}N$ | hNH, hCANH, hCONH, hcaCBcaNH, $R_{1\rho}(^{15}N)$ hCANH | *E. coli* | Cp149 dimer reassembled with 150 mM NaCl | 1f, S1, S3, S16 |
| Cp149 control | $^{13}C$-$^{15}N$ | DARR, NCA, NCO, NCACX, NCOCX, CANCO | *E. coli* | | 1d, 2, S4, S5 |
| Cp149 + JNJ-890 | $^{2}H$-$^{13}C$-$^{15}N$ | hNH, hCANH, hCONH, hcaCBcaNH, hCAcoNH, $R_{1\rho}(^{15}N)$ hCANH | *E. coli* | Cp149 dimer reassembled with CAMs for 24h at RT with 150 mM NaCl | 1f, S1, S3, S16 |
| Cp149 + JNJ-890 | $^{13}C$-$^{15}N$ | DARR, NCA, NCO, NCACX, NCOCX, CANCO | *E. coli* | | 1d, 2, S4, S5 |
| Cp149 + HAP_R10 | $^{2}H$-$^{13}C$-$^{15}N$ | hNH, hCANH, hCONH, hcaCBcaNH, hCAcoNH | *E. coli* | | S16 |
| Cp149 + HAP_R10 | $^{13}C$-$^{15}N$ | DARR, NCA, NCO, NCACX | *E. coli* | | 2, 3, S4, S5 |
| Cp149 + GS-942049 | $^{13}C$-$^{15}N$ | DARR, NCA, NCACX | *E. coli* | | 2, S4, S5 |
| Cp149 + GS-832471 | $^{13}C$-$^{15}N$ | DARR, NCA, NCACX | *E. coli* | | 2, S4, S5 |
| Cp149 + JNJ-827 | $^{13}C$-$^{15}N$ | DARR, NCA, NCO, NCACX, NCOCX, CANCO | *E. coli* | | 2, S4, S5 |
| Cp149 + JNJ-632 | $^{13}C$-$^{15}N$ | DARR, NCA, NCO, CANCO | *E. coli* | | 2, S2b, S4, S5 |
| Cp149 + HAP_R10 | $^{13}C$-$^{15}N$ | DARR, NCA, NCO, NCACX | *E. coli* | | 2, 3, S4, S5 |
| Cp149 capsids + JNJ-890 | $^{13}C$-$^{15}N$ | DARR, NCA, NCO | *E. coli* | Cp149 purified capsids incubated with 4 eq. CAMs 2h at 37 °C | S2a |
| Cp149 capsids + JNJ-632 | $^{13}C$-$^{15}N$ | DARR, NCA, NCO, NCACX, NCOCX | *E. coli* | | S2b |
| Cp183 control | $^{13}C$-$^{15}N$ | DARR, NCA | *E. coli* | | S9, S13 |
| Cp183 + JNJ-632 | $^{13}C$-$^{15}N$ | DARR, NCA, NCO | *E. coli* | Cp183 purified capsids incubated with 4 eq. CAMs 2h at 37 °C | S9, S13 |
| Cp183 + JNJ-890 | $^{13}C$-$^{15}N$ | DARR, NCA, NCO | *E. coli* | | S9, S13 |
| Cp183 + HAP_R10 | $^{13}C$-$^{15}N$ | DARR | *E. coli* | | S9, S13 |
| P7-Cp183 + JNJ-890 | $^{13}C$-$^{15}N$ | DARR | *E. coli* | | S9 |
| P7-Cp183 + HAP_R10 | $^{13}C$-$^{15}N$ | DARR, NCA, NCO, NCACX | *E. coli* | | 3, S9 |
| CF-Cp183 control | $^{2}H$-$^{13}C$-$^{15}N$ | hNH, HP-CP | Cell-free | | 4, S15 |
| CF-Cp183 + JNJ-632 | $^{2}H$-$^{13}C$-$^{15}N$ | hNH, HP-CP | Cell-free | 1 eq added upon synthesis | 4, S15 |
| CF-Cp183 + JNJ-890 | $^{2}H$-$^{13}C$-$^{15}N$ | hNH, HP-CP | Cell-free | 0.4 eq. added upon synthesis, sedimented from the pellet | 4, S15 |
| CF-Cp183 + HAP_R10 | $^{2}H$-$^{13}C$-$^{15}N$ | hNH, HP-CP | Cell-free | | 4, S15 |

**Table S2: Experimental parameters for ssNMR experiments using $^{13}C$-detection at 17.5 kHz MAS frequency.** Experiments were run on a 3.2 mm probe on a 800 MHz spectrometer at a temperature estimated around 4 °C. 90° pulses were: 2.5 µs in $^{1}H$, 5 µs in $^{13}C$ and 6.1 µs in $^{15}N$. For CP frequencies and transfer times, the average values with the standard deviation are given, which were calculated amongst the concerned samples as listed in Table S1. For increments, sw and acquisition times of 3D spectra as well as for the number of scans and experimental time, the range of the different values used are indicated. sw stands for spectral width.

| **Experiment** | **DARR** | **NCA** | **NCO** | **NCACX** | **NCOCX** | **CANCO** |
|---|---|---|---|---|---|---|
| MAS [kHz] | 17.5 | | | | | |
| Field [T] | | | | | | |
| **Transfer I** | HC-CP | HN-CP | HN-CP | HN-CP | HN-CP | HC-CP |
| $^{1}H$ field [kHz] | 65 ± 1 | 55.4 ± 0.4 | | | | 65 ± 1 |
| X field [kHz] | 50 ($^{13}C$) | 41 ($^{15}N$) | | | | 50 ($^{13}C$) |
| Shape | Tangent $^{1}H$ | | | | | |
| $^{13}C$ carrier [ppm] | 58.6 | | | | | |
| time [ms] | 0.80 ± 0.15 | 0.85 ± 0.15 | | | | 0.80 ± 0.15 |
| **Transfer II** | DARR | NCa-CP | NCO-CP | NCa-CP | NCO-CP | CaN-CP |
| Field [kHz] | 17.5 ($^{1}H$) | 6 ($^{13}C$) | | | | |
| Field [kHz] ($^{15}N$) | | 11.2 ± 0.2 | 11.8 ± 0.3 | 11.2 ± 0.2 | 11.8 ± 0.3 | 11.2 ± 0.2 |
| Shape | - | Tangent $^{13}C$ | | | | |
| $^{13}C$ carrier [ppm] | 100 | 58.6 | 177 | 58.6 | 177 | 58.6 |
| time [ms] | 20 | 9.7 ± 0.7 | 8 ± 2 | 9.7 ± 0.7 | 6.5 ± 1.0 | 9.7 ± 0.7 |
| **Transfer III** | | | | DARR | DARR | NCO-CP |
| Field [kHz] | | | | 17.5 ($^{1}H$) | 17.5 ($^{1}H$) | 6 ($^{13}C$) / 11.8 ± 0.3 ($^{15}N$) |
| Shape | | | | - | - | Tangent $^{13}C$ |
| Carrier [ppm] | | | | 58.6 | 177 | 177 |
| time [ms] | | | | 70 | 30 | 6.5 ± 1.0 |
| $t_1$ increments | 2560 | 1344 | 1344 | 74-90 | 32-50 | 38-70 |
| sw ($t_1$) [ppm] | 466 | 771 | 771 | 35 ($^{13}Ca$) | 15 ($^{13}CO$) | 40-60 ($^{15}N$) |
| Acq. time ($t_1$) [ms] | 13.7 | 10.8 | 10.8 | 5.2 - 6.3 | 9.1 | 5.3 - 7.2 |
| $t_2$ increments | 3072 | 2304 | 2304 | 44-58 | 34-50 | 66-80 |
| sw ($t_2$) [ppm] | 466 | 497 | 497 | 40-50 ($^{15}N$) | 40-42 ($^{15}N$) | 32-40 ($^{13}Ca$) |
| Acq. time ($t_2$) [ms] | 16.4 | 11.5 | 11.5 | 5.4-7.2 | 5.0-7.7 | 5.0-5.4 |
| $t_3$ increments | | | | 2304 | 2304 | 2304 |
| sw ($t_3$) [ppm] | | | | 497 | 497 | 497 |
| Acq. time ($t_3$) [ms] | | | | 11.5 | 11.5 | 11.5 |
| $^{1}H$ decoupling during acq. | SPINAL64 | | | | | |
| Field [kHz] | 90 | | | | | |
| Interscan delay [s] | 2.6 | 2.6 | 2.6 | 2.6 | 2.3 | 3 |
| Number of scans | 8-16 | 8-16 | 8-16 | 16-32 | 16-64 | 16 |
| Measurement time | 15-30h | 8-16h | 8-16h | 2-3 days | 1-3 days | 1-3 days |

**Table S3:** Experimental parameters for solid-state NMR experiments using $^{1}$H-detection at 60 kHz MAS frequency. Experiments were run on a 1.3 mm probe at a temperature estimated between 20 and 25 °C. WALTZ16 was used for $^{1}$H, $^{13}$C and $^{15}$N decoupling. Sw stands for spectra width. Carriers for $^{1}$H and $^{15}$N were at 4.8 and 118 ppm respectively.

| **Samples**<br>$^{2}$H-$^{13}$C-$^{15}$N | Cp149 control<br>Cp149 + JNJ-890<br>Cp149 + HAP_R10 | | | | | CF-Cp183 control<br>CF-Cp183 + JNJ-632<br>CF-Cp183 + JNJ-890<br>CF-Cp183 + HAP_R10 | |
|---|---|---|---|---|---|---|---|
| **Experiment** | **CP hNH 2D** | **hCANH 3D** | **hCONH 3D** | **hCAcoNH 3D** | **hcaCBcaNH 3D** | **CP hNH 2D** | **HP-CP 1D** |
| Field / T | 18.8 | | | | | 18.8 | 11.7 |
| MAS / kHz | 60 | | | | | 60 | 55-60 |
| 90° pulse (µs) | 2.5 ($^{1}$H) / 4 ($^{13}$C) / 4 ($^{15}$N) | | | | | 2.5($^{1}$H)/ 4($^{13}$C)/ 4($^{15}$N) | 2.5($^{1}$H)/ 5($^{31}$P)/ |
| **Transfer I** | **HN CP** | **HC CP** | | | | **HN CP** | **HP CP** |
| Field / kHz | 43 ($^{1}$H)/ 17 ($^{15}$N) | 25 ($^{1}$H) / 35 ($^{13}$C) | | | | 43 ($^{1}$H)/ 17 ($^{15}$N) | 100 ($^{1}$H)/ 40 ($^{31}$P) |
| Shape | Tangent $^{1}$H | Tangent $^{1}$H | | | | Tangent $^{1}$H | |
| Carrier $^{13}$C / ppm | - | 56 | 178 | 56 | 56 | - | - |
| Time / ms | ~0.7 | ~4 | 2.5 | ~4 | ~4 | 0.9 | 1 |
| **Transfer II** | **NH CP** | **CaN CP** | **CON CP** | **DREAM** | **CA-CB scalar** | **NH CP** | - |
| Field / kHz | 43 ($^{1}$H)/ 17 ($^{15}$N) | 25 ($^{15}$N) / 35 ($^{13}$C) | 25 ($^{15}$N) / 35 ($^{13}$C) | 30 ($^{13}$C) | - | 43 ($^{1}$H)/ 17 ($^{15}$N) | - |
| Shape | Tangent $^{1}$H | Tangent $^{13}$C | Tangent $^{13}$C | Tangent $^{13}$C | - | Tangent $^{1}$H | - |
| Carrier $^{13}$C / ppm | - | 56 | 178 | 178 | 42 | - | - |
| Time / ms | ~0.8 | ~8 | ~10 | ~10 | ~4 | 0.9 | - |
| **Transfer III** | - | **NH CP** | | **CaN CP** | **CaN CP** | - | - |
| Field / kHz | - | 43 ($^{1}$H)/ 17 ($^{15}$N) | | 25 ($^{15}$N) / 35 ($^{13}$C) | 25 ($^{15}$N) / 35 ($^{13}$C) | - | - |
| Shape | - | Tangent $^{1}$H | | Tangent $^{13}$C | Tangent $^{13}$C | - | - |
| Carrier $^{13}$C / ppm | - | - | - | 56 | 56 | - | - |
| Time / ms | - | ~0.8 | ~0.8 | ~8 | ~8 | - | - |
| **Transfer IV** | - | - | - | **NH CP** | **NH CP** | - | - |
| Field / kHz | - | - | - | 43 ($^{1}$H)/ 17 ($^{15}$N) | 43 ($^{1}$H)/ 17 ($^{15}$N) | - | - |
| Shape | - | - | - | Tangent $^{1}$H | Tangent $^{1}$H | - | - |
| Time / ms | - | - | - | ~0.8 | ~0.8 | - | - |
| t1 increments | 160 ($^{15}$N) | 50-80 ($^{13}$C) | 48-60 ($^{13}$C) | 75 ($^{13}$C) | 30-40 ($^{15}$N) | 320 | - |
| sw (t1) / ppm | 40 | 22-30 | 14-16 | 30 | 24-35 | 80 | - |
| Acq time (t1) / ms | 24.7 | ~7 | ~10 | 6.2 | ~8 | 24.7 | 10.2 |
| t2 increments | 2048 | 32-58 ($^{15}$N) | 40-64 ($^{15}$N) | 32 ($^{15}$N) | ~100 ($^{13}$C) | 2048 | - |
| sw (t2) / ppm | 100 | 24-40 | 30-40 | 24 | 50-55 | 97.7 | - |
| Acq time (t2) / ms | 12.9 | ~9 | ~10 | 8.2 | ~5 | 13 | - |
| t3 increments | - | 2048 | 2048 | 2048 | 2048 | - | - |
| sw (t3) / ppm | - | 100 | 100 | 100 | 100 | - | - |
| Acq time (t3) / ms | - | 12.9 | 12.9 | 12.9 | 12.9 | - | - |
| $^{1}$H dec. / kHz | 10 | 10 | 10 | 10 | 10 | 10 | 5 (WALTZ64) |
| $^{15}$N dec. / kHz | 10 | 5 | 5 | 5 | 10 | 10 | - |
| $^{13}$C dec. / kHz | 10 | 10 | 10 | 10 | 10 | 10 | - |
| Water sup. (100 ms) / kHz | 15 | 20 | 20 | 20 | 20 | 15 | - |
| Interscan delay / s | 1.5 | 1.5 | 1.5 | 1.4 | 1.2 | 1.5 | 1.5 |
| Number of scans | 80 | 32-64 | 32-80 | 64-80 | 48-128 | 80 | 41984-58408 |
| Experiment time | **11h30** | **2-3 days** | **2-3 days** | **2-3 days** | **3-6 days** | **~12h** | |

**Table S4:** Experimental parameters for measurement of site-specific 3D hCANH $R_{1\rho}(^{15}N)$ relaxation-rate constants shown in **Figure 1d**.

| **Sample** | **$^{2}H$-$^{13}C$-$^{15}N$-Cp149 + JNJ-890** | **$^{2}H$-$^{13}C$-$^{15}N$ -Cp149** |
|---|---|---|
| MAS frequency/ kHz | 80 | 80 |
| Field/ T | 20 | 20 |
| **Transfer I** | HC-CP(DQ) | HC-CP(DQ) |
| $^{1}H$ field/ kHz | 64 | 61 |
| $^{13}C$ field/ kHz | 14 | 15 |
| Shape | Tangent $^{1}H$ | Tangent $^{1}H$ |
| Carrier / ppm | 55 | 52 |
| Time/ ms | 5.25 | 4.5 |
| **Transfer II** | CN-CP | CN-CP |
| $^{13}C$ field/ kHz | 49 | 50 |
| $^{15}N$ field/ kHz | 30 | 29 |
| Shape | Tangent $^{13}C$ | Tangent $^{13}C$ |
| Carrier/ ppm | 117.5 | 117.5 |
| Time/ ms | 18 | 17 |
| **Transfer III** | NH-CP | NH-CP |
| $^{1}H$ field/ kHz | 60 | 59 |
| $^{15}N$ field/ kHz | 17 | 17 |
| Shape | Tangent $^{1}H$ | Tangent $^{1}H$ |
| Carrier/ ppm | 4.8 | 4.8 |
| Time/ ms | 2 | 2.4 |
| **$T_{1\rho}(^{15}N)$ Measurement** | 13 kHz Spin-Lock $^{15}N$ | 13 kHz Spin-Lock $^{15}N$ |
| Relaxation delays / ms | 0.001, 26, 51, 101, 126, 151, 201, 251 | 0.001, 26, 51, 101, 126, 151, 201, 251 |
| t1 increments | 50 | 50 |
| Sweep width (t1)/ ppm | 30 | 30 |
| Acquisition time (t1)/ ms | 3.9 | 3.9 |
| t2 increments | 30 | 30 |
| Sweep width (t2)/ ppm | 40 | 40 |
| Acquisition time (t2)/ ms | 4.4 | 4.4 |
| t3 increments | 2048 | 2048 |
| Sweep width (t3)/ ppm | 47 | 47 |
| Acquisition time (t3)/ ms | 25.8 | 25.8 |
| $^{1}H$ swfTPPM decoupling/ kHz | 10 | 10 |
| $^{15}N$ WALTZ64 decoupling/ kHz | 10 | 5 |
| $^{13}C$ WALTZ64 decoupling/ kHz | 5 | 5 |
| Water Suppression | MISSISIPPI | MISSISIPPI |
| $^{1}H$ field / kHz | 20 | 20 |
| Time / ms | 120 | 120 |
| Interscan delay/ s | 2.19 | 2.19 |
| Number of scans | 24 | 24 |
| Measurement time/ h | 184 | 184 |

**Table S5:** Carbon and nitrogen chemical shift perturbations values measured on 3D NCACX spectra for the six CAMs, as plotted in **Figure S5a**. For nitrogen, CSPs were calculated as

$$\Delta\delta_N = \left(\frac{\gamma_N}{\gamma_C}\right)|\delta_N[bound] - \delta_N[unbound]| = 0.4 * |\Delta\delta_N|$$

| ResNum | ResName | Atom | JNJ-890 | HAP_R10 | JNJ-632 | JNJ-827 | GS-832471 | GS-942049 |
|---|---|---|---|---|---|---|---|---|
| **2** | Asp | C | 0.2266 | 0.1270 | 0.0983 | 0.1883 | 0.0573 | 0.0911 |
| **2** | Asp | Ca | 0.0350 | 0.0393 | 0.0319 | 0.0689 | 0.0204 | 0.0697 |
| **2** | Asp | Cb | 0.0305 | 0.2112 | 0.0465 | 0.0248 | 0.0097 | 0.0002 |
| **2** | Asp | Cg | 0.0938 | 0.0880 | 0.0857 | 0.0686 | 0.0406 | 0.0830 |
| **2** | Asp | N | 0.2010 | 0.1873 | 0.0935 | 0.1332 | 0.0239 | 0.0733 |
| **3** | Ile | C | 0.2550 | 0.0502 | 0.4197 | 0.4444 | 0.0045 | 0.0478 |
| **3** | Ile | Ca | 0.0284 | 0.0899 | 0.4221 | 0.1647 | 0.4131 | 0.3457 |
| **3** | Ile | Cb | 0.4190 | 0.0493 | 0.6209 | 0.2713 | 0.0265 | 0.2767 |
| **3** | Ile | Cd1 | 0.2919 | 0.1042 | 0.5638 | 0.0513 | 0.2194 | 0.2918 |
| **3** | Ile | Cg1 | 0.0017 | 0.0074 | 0.1358 | 0.1330 | 0.0906 | 0.0444 |
| **3** | Ile | Cg2 | 0.0215 | 0.0272 | 0.0459 | 0.0339 | 0.0063 | 0.0242 |
| **3** | Ile | N | 0.0377 | 0.0428 | 0.0534 | 0.0247 | 0.0535 | 0.0058 |
| **4** | Asp | C | 0.2592 | 0.1413 | 0.3879 | 0.0260 | 0.2439 | 0.1560 |
| **4** | Asp | Ca | 0.1301 | 0.0144 | 0.2119 | 0.1680 | 0.0624 | 0.0311 |
| **4** | Asp | Cb | 0.1665 | 0.1352 | 0.1049 | 0.2126 | 0.1314 | 0.0291 |
| **4** | Asp | Cg | 0.1530 | 0.0389 | 0.0103 | 0.0063 | 0.1449 | 0.0523 |
| **4** | Asp | N | 0.1144 | 0.0523 | 0.2261 | 0.0476 | 0.0023 | 0.0300 |
| **5** | Pro | C | 0.0365 | 0.1507 | 0.1652 | 0.1226 | 0.2770 | 0.1330 |
| **5** | Pro | Ca | 0.0458 | 0.1236 | 0.0644 | 0.1099 | 0.0454 | 0.0336 |
| **5** | Pro | Cb |  | 0.0499 | 0.3851 |  | 0.0578 | 0.3571 |
| **5** | Pro | Cd |  | 0.2022 |  | 0.1901 | 0.1051 | 0.3128 |
| **5** | Pro | Cg |  | 0.1842 |  | 0.0592 | 0.1316 | 0.0988 |
| **5** | Pro | N | 0.0610 | 0.1572 | 0.1658 | 0.0308 | 0.1297 | 0.0533 |
| **6** | Tyr | C | 0.1839 | 0.0571 | 0.3095 | 0.0493 | 0.0767 | 0.0809 |
| **6** | Tyr | Ca | 0.2147 | 0.2196 | 0.1849 | 0.0063 | 0.2009 | 0.1416 |
| **6** | Tyr | Cb | 0.4115 | 0.7642 | 0.2664 |  | 0.3213 | 0.5839 |
| **6** | Tyr | Cd |  |  | 0.0002 | 0.0000 |  | 0.1002 |
| **6** | Tyr | N | 0.2977 | 0.1759 | 0.3742 | 0.0447 | 0.1353 | 0.1226 |
| **7** | Lys | C | 0.0316 | 0.0123 | 0.0134 | 0.0105 | 0.0153 | 0.0358 |
| **7** | Lys | Ca | 0.3869 | 0.3564 | 0.1682 | 0.1832 | 0.2913 | 0.2991 |
| **7** | Lys | Cb | 0.3099 | 0.1903 | 0.0822 | 0.0880 | 0.1701 | 0.2102 |
| **7** | Lys | Cd | 0.1857 | 0.0445 | 0.0036 | 0.0687 | 0.1337 | 0.0231 |
| **7** | Lys | Cg | 0.1739 | 0.1044 | 0.0509 | 0.0332 | 0.0529 | 0.0489 |
| **7** | Lys | N | 0.0096 | 0.0722 | 0.1621 | 0.0169 | 0.0120 | 0.0885 |
| **8** | Glu | C |  | 0.0956 | 0.2230 | 0.3225 | 0.2307 | 0.2154 |
| **8** | Glu | Ca | 0.1518 | 0.1613 | 0.0165 | 0.2721 | 0.0598 | 0.0241 |
| **8** | Glu | Cb | 0.2011 | 0.1236 | 0.0001 | 0.2433 | 0.0137 | 0.0371 |
| **8** | Glu | Cd | 0.1764 | 0.0797 | 0.1617 | 0.0598 | 0.0110 | 0.0683 |
| **8** | Glu | Cg | 0.3558 | 0.3210 | 0.1506 | 0.0554 | 0.1453 | 0.3358 |
| **8** | Glu | N | 0.0925 | 0.0770 | 0.0377 | 0.0304 | 0.0619 | 0.0322 |
| **9** | Phe | C | 0.2738 | 0.1050 | 0.2774 | 0.2977 | 0.3996 | 0.3060 |
| **9** | Phe | Ca | 0.0954 | 0.1373 | 0.1187 | 0.1071 | 0.1384 | 0.1276 |
| **9** | Phe | Cb | 0.2265 | 0.2928 | 0.1496 | 0.0750 | 0.2847 | 0.3107 |
| **9** | Phe | Cg |  | 0.1091 | 0.0002 |  |  |  |
| **9** | Phe | N | 0.1033 | 0.1063 | 0.1234 | 0.1355 | 0.1817 | 0.2165 |
| **10** | Gly | C | 0.2295 | 0.1172 | 0.1269 | 0.1677 | 0.1201 | 0.0200 |
| **10** | Gly | Ca | 0.0960 | 0.0653 | 0.0502 | 0.0307 | 0.0540 | 0.0404 |
| **10** | Gly | N | 0.3295 | 0.0768 | 0.2861 | 0.4722 | 0.2049 | 0.1849 |
| **11** | Ala | C | 0.2443 | 0.2542 | 0.0696 | 0.0823 | 0.2082 | 0.1766 |
| **11** | Ala | Ca | 0.1138 | 0.1275 | 0.0213 | 0.0068 | 0.1236 | 0.0754 |
| **11** | Ala | Cb | 0.5935 | 0.6759 | 0.6865 | 0.5600 | 0.8192 | 0.7565 |

| | | | | | | | | |
|---|---|---|---|---|---|---|---|---|
| **11** | Ala | N | 0.2081 | 0.0588 | 0.1226 | 0.0294 | 0.0494 | 0.0149 |
| **12** | Thr | C | 0.0773 | 0.0461 | 0.0024 | 0.0086 | 0.0338 | 0.1058 |
| **12** | Thr | Ca | 0.1245 | 0.1276 | 0.1733 | 0.1258 | 0.1043 | 0.0727 |
| **12** | Thr | Cb | 0.2099 | 0.0977 | 0.0442 | 0.0469 | 0.0756 | 0.1682 |
| **12** | Thr | Cg2 | 0.2013 | 0.0571 | 0.0122 | 0.0768 | 0.0744 | 0.0931 |
| **12** | Thr | N | 0.1933 | 0.1970 | 0.1391 | 0.1559 | 0.2829 | 0.2219 |
| **13** | Val | C | 0.2120 | 0.1750 | 0.0406 | 0.1086 | 0.0003 | 0.0016 |
| **13** | Val | Ca | 0.1550 | 0.1195 | 0.0656 | 0.1164 | 0.0385 | 0.0095 |
| **13** | Val | Cb | 0.0174 | 0.0403 | 0.1124 | 0.0485 | 0.0168 | 0.0104 |
| **13** | Val | Cga | 0.2866 | 0.0918 | 0.5715 | 0.4165 | 0.2544 | 0.2855 |
| **13** | Val | N | 0.3594 | 0.3559 | 0.0369 | 0.1210 | 0.0361 | 0.0546 |
| **14** | Glu | C | 0.1389 | 0.0478 | 0.1720 | 0.9152 | 0.5134 | 0.5134 |
| **14** | Glu | Ca | 1.0563 | 1.0614 | 0.9233 | 0.4810 | 0.9547 | 0.9596 |
| **14** | Glu | Cb | 0.2861 | 0.2071 | 0.0766 | 0.0500 | 0.0503 | 0.1226 |
| **14** | Glu | Cd | 0.3500 | 0.3758 | 0.0775 | 1.1061 | 0.0942 | 0.0942 |
| **14** | Glu | Cg | 0.8886 | 0.9111 | 0.7273 | 0.4763 | 0.7479 | 0.7479 |
| **14** | Glu | N | 0.2795 | 0.2096 | 0.1967 | 0.0264 | 0.3038 | 0.3074 |
| **15** | Leu | C | 0.6510 | 0.5194 | 0.5272 | | 0.5945 | 0.5901 |
| **15** | Leu | Ca | 0.3869 | 0.3936 | 0.2515 | 0.9062 | 0.2722 | 0.2666 |
| **15** | Leu | Cb | 1.3272 | 0.9466 | 0.4420 | 0.1332 | 0.4961 | 0.6731 |
| **15** | Leu | Cda | 0.6297 | 0.1322 | | 0.2490 | | 1.2429 |
| **15** | Leu | N | 0.2650 | 0.3272 | 0.1810 | 0.0880 | 0.2436 | 0.2540 |
| **16** | Leu | C | 0.2429 | 0.1947 | 0.1335 | 0.1523 | 0.1414 | 0.1362 |
| **16** | Leu | Ca | 0.8226 | 0.7583 | 0.5617 | 0.6795 | 0.6982 | 0.5970 |
| **16** | Leu | Cb | 1.2909 | 1.0643 | 0.7560 | 0.9268 | 0.9603 | 0.9871 |
| **16** | Leu | Cg | 0.8115 | 0.6450 | 0.8095 | 0.7469 | 0.7397 | 0.7681 |
| **16** | Leu | N | 0.4248 | 0.4214 | 0.5180 | 0.4524 | 0.6049 | 0.5532 |
| **17** | Ser | C | 0.9907 | 1.0308 | 1.2667 | 1.1040 | 1.1053 | 1.3131 |
| **17** | Ser | Ca | 0.6211 | 0.7455 | 0.8198 | 0.7765 | 0.6990 | 0.8038 |
| **17** | Ser | Cb | 0.1843 | 0.2394 | 0.2429 | 0.2733 | 0.2057 | 0.2067 |
| **17** | Ser | N | 0.3353 | 0.6232 | 0.6684 | 0.6778 | 0.5802 | 0.7561 |
| **18** | Phe | Ca | 0.3118 | 0.3699 | 0.1145 | 0.2429 | | |
| **18** | Phe | Cb | 0.0771 | 0.0747 | | 0.0915 | | |
| **18** | Phe | N | 0.6713 | 0.7525 | 0.0146 | 0.5871 | | |
| **20** | Pro | C | | 0.0376 | 0.6606 | | | |
| **20** | Pro | Ca | 0.2116 | 0.4013 | 0.2758 | | 0.2236 | 0.5108 |
| **20** | Pro | Cb | 1.3266 | 1.0463 | 0.8251 | | 1.1338 | |
| **20** | Pro | Cd | | | 1.1355 | | 1.1160 | |
| **20** | Pro | Cg | | 0.3227 | 0.2389 | | 0.4516 | |
| **20** | Pro | N | 0.4695 | 0.5178 | 0.5482 | | 0.5721 | 0.4742 |
| **21** | Ser | C | 0.3946 | 0.4333 | 0.4920 | 0.5829 | 0.5644 | 0.6305 |
| **21** | Ser | Ca | 0.0539 | 0.1554 | 0.1057 | 0.0169 | 0.0808 | 0.0592 |
| **21** | Ser | Cb | 0.0237 | 0.1963 | 0.0513 | 0.0617 | 0.0239 | 0.0563 |
| **21** | Ser | N | 0.2776 | 0.5110 | 0.0410 | 0.0087 | 0.2765 | 0.0229 |
| **22** | Asp | C | 0.0833 | 0.2702 | 0.1558 | 0.1250 | 0.2621 | 0.1840 |
| **22** | Asp | Ca | 0.6153 | 0.4296 | 0.4643 | 0.5181 | 0.7037 | 0.5498 |
| **22** | Asp | Cb | 0.0233 | 0.3297 | 0.3386 | 0.3010 | 0.1340 | 0.1905 |
| **22** | Asp | Cg | 0.4083 | 0.1713 | 0.6336 | 0.6181 | 0.6524 | 0.6374 |
| **22** | Asp | N | 0.1620 | 0.1911 | 0.0130 | 0.2656 | 0.3077 | 0.2577 |
| **23** | Phe | C | 1.0038 | 0.8085 | 0.4684 | 0.1587 | 0.2377 | 0.1924 |
| **23** | Phe | Ca | 0.8587 | 0.5792 | 0.0936 | 0.3603 | 0.0586 | 0.2273 |
| **23** | Phe | Cb | 1.1482 | 0.8273 | 0.0453 | 0.6052 | 0.4090 | 0.3190 |
| **23** | Phe | Cd | | | 0.0003 | 0.0000 | | 0.1003 |
| **23** | Phe | Ce | | | 0.0002 | 0.0000 | | 0.1002 |
| **23** | Phe | N | 0.4592 | 0.4663 | 0.0685 | 0.2266 | 0.2947 | 0.1656 |
| **24** | Phe | C | 1.4041 | 1.3624 | 0.7137 | 0.6426 | | |
| **24** | Phe | Ca | 1.1420 | 1.0216 | 0.2711 | 0.1424 | 0.2328 | 0.2939 |
| **24** | Phe | N | 1.0334 | 1.2526 | 0.1642 | 0.0791 | 0.0847 | 0.0853 |

| | | | | | | | | |
|---|---|---|---|---|---|---|---|---|
| **25** | Pro | Ca | 0.0085 | 0.0932 | 0.1207 | 0.1530 | 0.0048 | 0.1583 |
| **25** | Pro | Cb | | 0.2549 | 0.0107 | 0.2406 | 0.0531 | 0.0265 |
| **25** | Pro | Cd | 0.0266 | 0.0461 | 0.2911 | 0.0457 | 0.0618 | 0.0219 |
| **25** | Pro | Cg | 0.5845 | | 0.2168 | | 0.5043 | 0.1709 |
| **25** | Pro | N | 0.2595 | 0.2803 | 0.8483 | 0.8871 | 0.9312 | 0.8554 |
| **26** | Ser | C | 0.3116 | 0.0233 | 0.0664 | 0.2396 | 0.0427 | 0.0623 |
| **26** | Ser | Ca | 0.1186 | 0.1106 | 0.1313 | 0.0462 | 0.0349 | 0.0521 |
| **26** | Ser | Cb | 0.0900 | 0.0459 | 0.1531 | 0.3160 | 0.0921 | 0.0605 |
| **26** | Ser | N | 0.3448 | 0.3387 | 0.0279 | 0.3213 | 0.1583 | 0.1770 |
| **27** | Val | C | 0.0070 | 0.1235 | 0.1998 | 0.3873 | 0.1801 | 0.1591 |
| **27** | Val | Ca | 0.2143 | 0.0871 | 0.0210 | 0.0913 | 0.0027 | 0.1629 |
| **27** | Val | Cb | | 0.0100 | 0.2198 | 0.0391 | 0.0834 | 0.1027 |
| **27** | Val | Cga | 0.0957 | 0.2467 | 0.1754 | 0.0512 | 0.1661 | 0.1018 |
| **27** | Val | Cgb | 0.0456 | 0.0839 | 0.8695 | 0.0985 | 0.2378 | 0.2754 |
| **27** | Val | N | 0.0502 | 0.0296 | 0.0141 | 0.0790 | 0.2179 | 0.0046 |
| **28** | Arg | C | 0.2833 | 0.1489 | 0.1895 | 0.0761 | 0.2517 | 0.0752 |
| **28** | Arg | Ca | 0.1249 | 0.1613 | 0.1808 | 0.2229 | 0.1175 | 0.0074 |
| **28** | Arg | Cb | 0.2236 | 0.2119 | 0.1414 | 0.1152 | 0.1608 | 0.0113 |
| **28** | Arg | Cg | 0.0647 | 0.0353 | 0.1044 | 0.0552 | 0.0871 | 0.0716 |
| **28** | Arg | Cd | | | | | 0.2458 | |
| **28** | Arg | Cz | | | | 0.0000 | | 0.0995 |
| **28** | Arg | N | 0.2655 | 0.2766 | 0.0055 | 0.4445 | 0.1548 | 0.1346 |
| **29** | Asp | C | 0.0701 | 0.0246 | 0.0684 | 0.7752 | 0.2210 | 0.1712 |
| **29** | Asp | Ca | 0.0761 | 0.1780 | 0.0111 | 0.1784 | 0.0950 | 0.1582 |
| **29** | Asp | Cb | 0.0201 | 0.5254 | 0.1752 | 0.5035 | 0.1909 | 0.0848 |
| **29** | Asp | N | 0.0351 | 0.0316 | 0.0979 | 0.1849 | 0.0021 | 0.0011 |
| **30** | Leu | C | 0.2162 | 0.3401 | 0.1447 | 0.2737 | 0.1434 | 0.2152 |
| **30** | Leu | Ca | 0.1336 | 0.9213 | 0.1815 | 0.1047 | 0.1133 | 0.0470 |
| **30** | Leu | Cb | 0.0091 | 0.2193 | 0.2564 | 0.5219 | 0.1135 | 0.1124 |
| **30** | Leu | Cda | | 1.9935 | 0.4276 | 0.7104 | 0.6445 | 0.7538 |
| **30** | Leu | Cg | 1.4983 | 0.8805 | 0.6417 | 0.7255 | 0.7925 | 0.8786 |
| **30** | Leu | N | 0.6805 | 0.6355 | 0.4440 | 0.0883 | 0.4771 | 0.2982 |
| **31** | Leu | C | 0.0813 | 0.0686 | 0.0470 | 0.3039 | 0.1507 | 0.1245 |
| **31** | Leu | Ca | 0.0374 | 0.0417 | 0.0113 | 0.4141 | 0.0260 | 0.0331 |
| **31** | Leu | Cb | 0.0194 | 0.0236 | 0.1747 | 0.0951 | 0.1798 | 0.0968 |
| **31** | Leu | Cda | 0.3666 | 0.3717 | 0.2057 | 0.2861 | 0.4260 | 0.2593 |
| **31** | Leu | Cg | 0.0483 | 0.0227 | 0.0581 | 0.7611 | 0.0641 | 0.0356 |
| **31** | Leu | N | 0.2912 | 0.2720 | 0.1648 | 0.0072 | 0.2498 | 0.0632 |
| **32** | Asp | C | 0.0973 | 0.1258 | 0.2739 | 0.0093 | 0.1554 | 0.2195 |
| **32** | Asp | Ca | 0.1280 | 0.0488 | 0.2349 | 0.1301 | 0.1287 | 0.1589 |
| **32** | Asp | Cb | 0.4437 | 0.4808 | 0.3474 | 0.1041 | 0.4579 | 0.4594 |
| **32** | Asp | Cg | 0.4389 | 0.3046 | 0.1442 | 0.0084 | 0.1685 | 0.1686 |
| **32** | Asp | N | 0.2781 | 0.0773 | 0.0885 | 0.3178 | 0.0397 | 0.1203 |
| **33** | Thr | C | 0.3723 | 0.2656 | 0.1918 | 0.0007 | 0.2365 | 0.3685 |
| **33** | Thr | Ca | 0.0818 | 0.0449 | 0.6131 | 0.6053 | 0.6649 | 0.6367 |
| **33** | Thr | Cb | 0.1622 | 0.3868 | 0.3005 | 0.3495 | 0.2319 | |
| **33** | Thr | Cg2 | 0.0896 | 0.4318 | 0.0099 | 0.8157 | 0.4458 | 0.5612 |
| **33** | Thr | N | 0.3333 | 0.4120 | 0.0164 | 0.4636 | 0.4784 | 0.1072 |
| **34** | Ala | C | 0.3118 | 0.2746 | 0.3510 | 0.3151 | 0.4397 | 0.5248 |
| **34** | Ala | Ca | 0.0898 | 0.2276 | 0.0732 | 0.1910 | 0.2270 | 0.1596 |
| **34** | Ala | Cb | 0.2853 | 0.0547 | 0.0870 | 0.0163 | 0.0263 | 0.1495 |
| **34** | Ala | N | 0.2325 | 0.1282 | 0.0150 | 0.0575 | 0.1754 | 0.0567 |
| **35** | Ser | C | 0.2351 | 0.3495 | 0.0811 | 0.0050 | 0.1891 | 0.0418 |
| **35** | Ser | Ca | 0.3572 | 0.3593 | 0.2453 | 0.2125 | 0.2551 | 0.1442 |
| **35** | Ser | Cb | 0.0644 | 0.0495 | 0.0517 | 0.1108 | 0.0427 | 0.0176 |
| **35** | Ser | N | 0.7939 | 0.7001 | 0.9471 | 0.7582 | 1.0758 | 0.9340 |
| **36** | Ala | C | 0.5648 | 0.4820 | 0.3088 | 0.3030 | 0.3427 | 0.8500 |
| **36** | Ala | Ca | 0.1708 | 0.2118 | 0.2652 | 0.3090 | 0.2758 | 0.3003 |

| | | | | | | | | |
|---|---|---|---|---|---|---|---|---|
| **36** | Ala | Cb | 3.2432 | 3.0663 | 2.3977 | 1.9244 | 2.4734 | 3.0003 |
| **36** | Ala | N | 0.1440 | 0.1195 | 0.0144 | 0.0207 | 0.0097 | 0.0065 |
| **37** | Leu | C | 0.2988 | 0.2296 | 0.1957 | 0.2938 | 0.3759 | 0.2750 |
| **37** | Leu | Ca | 1.0666 | 1.0438 | 0.9753 | 0.2268 | 1.5040 | 0.9664 |
| **37** | Leu | Cb | 0.2703 | 0.2612 | 1.3010 | 1.5255 | 1.1395 | 1.8443 |
| **37** | Leu | Cdb | | | 1.2599 | | | |
| **37** | Leu | Cg | 0.9192 | 0.6696 | 1.7448 | 0.3535 | 1.4937 | 1.5260 |
| **37** | Leu | N | 0.2043 | 0.1837 | 0.1689 | 0.6345 | 0.2169 | 0.3153 |
| **38** | Tyr | C | | 0.1774 | 0.0434 | 0.0826 | 0.1584 | 0.1319 |
| **38** | Tyr | Ca | 0.1522 | 0.1134 | 0.0650 | 0.0571 | 0.0060 | 0.1467 |
| **38** | Tyr | Cb | 0.0597 | | 0.0279 | 0.0781 | 0.7864 | 0.0568 |
| **38** | Tyr | Cg | | | 0.1798 | 0.1002 | | |
| **38** | Tyr | N | 0.1316 | 0.1134 | 0.2467 | 0.1123 | 0.0234 | 0.0054 |
| **39** | Arg | C | 0.0206 | 0.1311 | 0.0049 | 0.0801 | 0.0804 | 0.0760 |
| **39** | Arg | Ca | 0.0897 | 0.0149 | 0.1520 | 0.1437 | 0.0813 | 0.0857 |
| **39** | Arg | Cb | 0.3046 | 0.1683 | 0.0103 | 0.0287 | 0.1549 | 0.1618 |
| **39** | Arg | Cd | 0.1123 | 0.0253 | 0.0477 | 0.0591 | 0.0528 | 0.0562 |
| **39** | Arg | Cg | 0.2095 | 0.0791 | 0.0907 | 0.0527 | 0.1987 | 0.1884 |
| **39** | Arg | N | 0.1722 | 0.1565 | 0.0188 | 0.0373 | 0.0206 | 0.0086 |
| **40** | Glu | C | 0.0949 | 0.0047 | 0.0316 | 0.0045 | 0.0402 | 0.0041 |
| **40** | Glu | Ca | 0.1165 | 0.0638 | 0.0968 | 0.0493 | 0.0536 | 0.0198 |
| **40** | Glu | Cb | 0.0467 | 0.1119 | 0.0092 | 0.0107 | 0.0062 | 0.0394 |
| **40** | Glu | Cd | | | 0.7119 | 0.0000 | | 0.0903 |
| **40** | Glu | Cg | 0.1810 | 0.1348 | 0.0271 | 0.1523 | 0.0510 | 0.0964 |
| **40** | Glu | N | 0.0278 | 0.1060 | 0.0311 | 0.0393 | 0.0280 | 0.0629 |
| **41** | Ala | C | 0.2444 | 0.1867 | 0.0379 | 0.0540 | 0.0513 | 0.0437 |
| **41** | Ala | Ca | 0.1685 | 0.0104 | 0.0708 | 0.0576 | 0.0353 | 0.0225 |
| **41** | Ala | Cb | 0.1153 | 0.0195 | 0.0253 | 0.0393 | 0.0689 | 0.1230 |
| **41** | Ala | N | 0.1088 | 0.0750 | 0.0820 | 0.0341 | 0.0200 | 0.0649 |
| **42** | Leu | C | | 0.1024 | 0.0649 | 0.0015 | 0.0043 | 0.0275 |
| **42** | Leu | Ca | 0.0314 | 0.0849 | 0.0875 | 0.0894 | 0.0711 | 0.0765 |
| **42** | Leu | Cb | 0.0697 | 0.1083 | 0.0991 | 0.0470 | 0.0254 | 0.0949 |
| **42** | Leu | Cdb | | 0.1770 | 0.0960 | 0.0415 | 0.1280 | 0.1265 |
| **42** | Leu | Cg | 0.0301 | 0.0582 | 0.0142 | 0.0092 | 0.0206 | 0.0195 |
| **42** | Leu | N | 0.0739 | 0.1305 | 0.0679 | 0.1796 | 0.0064 | 0.0241 |
| **43** | Glu | C | 0.0424 | 0.1493 | 0.0686 | 0.0844 | 0.0415 | 0.0032 |
| **43** | Glu | Ca | 0.0365 | 0.0009 | 0.1029 | 0.0530 | 0.0234 | 0.0076 |
| **43** | Glu | Cb | 0.0893 | 0.0442 | 0.0129 | 0.0849 | 0.0864 | 0.0753 |
| **43** | Glu | Cd | 0.0466 | 0.0719 | 0.0844 | 0.0561 | 0.0531 | 0.0187 |
| **43** | Glu | Cg | 0.0466 | 0.0860 | 0.0577 | 0.0486 | 0.0161 | 0.0180 |
| **43** | Glu | N | 0.1184 | 0.0058 | 0.0792 | 0.1066 | 0.0446 | 0.0261 |
| **44** | Ser | C | 0.0724 | 0.1661 | 0.0052 | 0.0148 | 0.0155 | 0.0152 |
| **44** | Ser | Ca | 0.2146 | 0.0961 | 0.0998 | 0.0434 | 0.0349 | 0.0576 |
| **44** | Ser | Cb | 0.0195 | 0.1457 | 0.0304 | 0.0243 | 0.0121 | 0.0295 |
| **44** | Ser | N | 0.1091 | 0.1284 | 0.0824 | 0.0252 | 0.0156 | 0.0495 |
| **45** | Pro | C | | 0.0661 | 0.0518 | 0.0821 | 0.0122 | 0.1597 |
| **45** | Pro | Ca | 0.0245 | 0.0009 | 0.0064 | 0.0171 | 0.0365 | 0.0369 |
| **45** | Pro | Cb | 0.1999 | 0.0063 | 0.0936 | 0.0222 | 0.1082 | 0.1368 |
| **45** | Pro | Cd | | 0.0362 | 0.0611 | 0.0205 | 0.0387 | 0.0397 |
| **45** | Pro | Cg | | 0.1492 | 0.0176 | 0.0432 | 0.0683 | 0.0545 |
| **45** | Pro | N | 0.0354 | 0.0161 | 0.0423 | 0.0360 | 0.0741 | 0.0156 |
| **46** | Glu | C | 0.0814 | 0.1617 | 0.0574 | 0.1574 | 0.0922 | 0.0215 |
| **46** | Glu | Ca | 0.2002 | 0.1328 | 0.0039 | 0.2396 | 0.0321 | 0.0602 |
| **46** | Glu | Cb | 0.2316 | 0.1816 | 0.1687 | 0.1262 | 0.1269 | 0.1002 |
| **46** | Glu | Cd | 0.2623 | 0.0697 | 0.2348 | 0.2059 | 0.1043 | 0.1385 |
| **46** | Glu | Cg | 0.2797 | 0.0594 | 0.0304 | 0.1828 | 0.1410 | 0.1827 |
| **46** | Glu | N | 0.0506 | 0.0661 | 0.0949 | 0.1379 | 0.0484 | 0.0899 |
| **47** | His | C | 0.3492 | 0.0958 | 0.1802 | 0.3205 | 0.1323 | 0.0428 |

| | | | | | | | | |
|---|---|---|---|---|---|---|---|---|
| **47** | His | Ca | 0.1272 | 0.0946 | 0.0881 | 0.1298 | 0.1372 | 0.1361 |
| **47** | His | Cb | 0.2595 | 0.5903 | 0.5789 | 0.6129 | 0.3329 | 0.3002 |
| **47** | His | Cd2 | | | 0.0005 | | | 0.0585 |
| **47** | His | N | 0.2320 | 0.2225 | 0.1233 | 0.3127 | 0.3015 | 0.3242 |
| **48** | Cys | C | 0.0244 | 0.0361 | 0.1566 | 0.1165 | 0.0203 | 0.0200 |
| **48** | Cys | Ca | 0.0916 | 0.0936 | 0.1082 | 0.1010 | 0.0133 | 0.0194 |
| **48** | Cys | Cb | 0.0637 | 0.0306 | 0.0742 | 0.0383 | 0.0373 | 0.1046 |
| **48** | Cys | N | 0.0224 | 0.1300 | 0.0512 | 0.2301 | 0.0825 | 0.0158 |
| **49** | Ser | C | 0.0649 | 0.0299 | 0.0525 | 0.0275 | 0.0095 | 0.0154 |
| **49** | Ser | Ca | 0.1343 | 0.0223 | 0.1204 | 0.0551 | 0.0091 | 0.0149 |
| **49** | Ser | Cb | 0.2168 | 0.2021 | 0.0538 | 0.0358 | 0.0253 | 0.0336 |
| **49** | Ser | N | 0.0277 | 0.1417 | 0.0297 | 0.0685 | 0.0232 | 0.0640 |
| **50** | Pro | C | 0.0284 | 0.1762 | 0.2202 | 0.1190 | 0.0351 | 0.0498 |
| **50** | Pro | Ca | 0.0564 | 0.0974 | 0.1393 | 0.1402 | 0.0494 | 0.1057 |
| **50** | Pro | Cb | | 0.0805 | 0.2388 | | 0.4451 | 0.6640 |
| **50** | Pro | Cd | | 0.0654 | 0.0320 | 0.0442 | 0.0945 | 0.0852 |
| **50** | Pro | Cg | | 0.0097 | 0.0524 | 0.0575 | 0.1560 | 0.1602 |
| **50** | Pro | N | 0.0724 | 0.0537 | 0.0115 | 0.0281 | 0.0048 | 0.0064 |
| **51** | His | C | 0.0603 | 0.1110 | 0.0022 | 0.0388 | 0.0911 | 0.0302 |
| **51** | His | Ca | 0.0923 | 0.1195 | 0.2416 | 0.1686 | 0.0303 | 0.0880 |
| **51** | His | Cb | 0.2194 | 0.1607 | 0.1619 | 0.2335 | 0.1046 | 0.0705 |
| **51** | His | Ce1 | 0.3001 | 0.3001 | | 0.0000 | 0.0127 | 0.0964 |
| **51** | His | Cg | 0.0428 | 0.0449 | 0.0394 | 0.0654 | 0.0259 | 0.3850 |
| **51** | His | N | 0.1729 | 0.2342 | 0.0788 | 0.0825 | 0.0179 | 0.0366 |
| **52** | His | C | 0.0879 | 0.0635 | 0.0750 | 0.0945 | 0.0274 | 0.0498 |
| **52** | His | Ca | 0.0685 | 0.1555 | 0.2509 | 0.1917 | 0.0554 | 0.0931 |
| **52** | His | Cb | 0.0745 | 0.2291 | 0.0522 | 0.0804 | 0.0663 | 0.0603 |
| **52** | His | Cd2 | | 0.0657 | 0.0004 | 0.0000 | 0.1004 | 0.1004 |
| **52** | His | Ce1 | | | 0.0135 | 0.0135 | 0.0578 | 0.0509 |
| **52** | His | Cg | | 0.0053 | 0.0123 | 0.0185 | 0.0600 | 0.0859 |
| **52** | His | N | 0.1250 | 0.0092 | 0.1804 | 0.1584 | 0.0378 | 0.1223 |
| **53** | Thr | C | 0.0042 | 0.0449 | 0.0876 | 0.0754 | 0.0399 | 0.0070 |
| **53** | Thr | Ca | 0.0812 | 0.1358 | 0.1708 | 0.0440 | 0.0138 | 0.0512 |
| **53** | Thr | Cb | 0.0248 | 0.1309 | 0.0042 | 0.0262 | 0.0321 | 0.0014 |
| **53** | Thr | Cg2 | 0.0497 | 0.0041 | 0.2437 | 0.0365 | 0.0584 | 0.0933 |
| **53** | Thr | N | 0.2041 | 0.1216 | 0.0935 | 0.1394 | 0.0024 | 0.0635 |
| **54** | Ala | C | 0.4093 | 0.2401 | 0.3798 | 0.3505 | 0.3158 | 0.4246 |
| **54** | Ala | Ca | 0.0071 | 0.0044 | 0.0504 | 0.1128 | 0.0690 | 0.0289 |
| **54** | Ala | Cb | 0.2443 | 0.2680 | 0.0122 | 0.0698 | 0.0806 | 0.0774 |
| **54** | Ala | N | 0.0431 | 0.0792 | 0.1027 | 0.0866 | 0.0275 | 0.0408 |
| **55** | Leu | C | 0.0076 | 0.0942 | 0.0996 | 0.1232 | 0.4602 | 0.4687 |
| **55** | Leu | Ca | 0.2160 | 0.4471 | 0.0631 | 0.0279 | 0.3505 | 0.3410 |
| **55** | Leu | Cb | 0.2145 | 0.1223 | 0.4401 | 1.0514 | 0.4280 | 0.4345 |
| **55** | Leu | Cg | 0.0181 | 0.6753 | 0.0100 | 0.0254 | 0.0932 | 0.1160 |
| **55** | Leu | N | 0.2031 | 0.1344 | 0.5903 | 0.6754 | 0.6668 | 0.6649 |
| **56** | Arg | C | 0.2119 | 0.0168 | 0.2088 | 0.2373 | 0.1064 | 0.0830 |
| **56** | Arg | Ca | 0.0646 | 0.0750 | 0.0228 | 0.0052 | 0.0204 | 0.0180 |
| **56** | Arg | Cb | 0.0718 | 0.1369 | 0.0213 | 0.0534 | 0.0352 | 0.0406 |
| **56** | Arg | Cd | 0.0283 | 0.0719 | 0.0236 | 0.0865 | 0.0530 | 0.0220 |
| **56** | Arg | Cg | 0.1083 | 0.0869 | 0.2669 | | 0.0714 | 0.2187 |
| **56** | Arg | Cz | | 0.0675 | | 0.0387 | 0.1084 | 0.1731 |
| **56** | Arg | N | 0.1316 | 0.1071 | 0.0925 | 0.0999 | 0.0083 | 0.0807 |
| **57** | Gln | C | 0.1161 | 0.1580 | 0.1988 | 0.1627 | 0.3203 | 0.3197 |
| **57** | Gln | Ca | 0.1240 | 0.0168 | 0.2954 | 0.1119 | 0.0853 | 0.0515 |
| **57** | Gln | Cb | 0.0546 | 0.0161 | 0.4956 | 0.1624 | | 0.2127 |
| **57** | Gln | N | 0.1506 | 0.1524 | 0.1589 | 0.2727 | 0.1928 | 0.1659 |
| **58** | Ala | C | | 0.1867 | 0.2190 | 0.0676 | 0.2771 | 0.3046 |
| **58** | Ala | Ca | 0.1700 | 0.1145 | 0.1845 | 0.1470 | 0.0726 | 0.0431 |

| | | | | | | | | |
|---|---|---|---|---|---|---|---|---|
| **58** | Ala | Cb | 0.4615 | 0.1547 | 0.8639 | 0.6813 | 0.3556 | 0.3095 |
| **58** | Ala | N | 0.5516 | 0.0053 | 0.6106 | 1.1003 | 0.2050 | 0.0670 |
| **59** | Ile | C | 0.1105 | 0.1402 | 0.0250 | 0.0242 | 0.0441 | 0.0446 |
| **59** | Ile | Ca | 0.1403 | 0.0110 | 0.1958 | 0.1250 | 0.1491 | 0.0442 |
| **59** | Ile | Cb | 0.0213 | 0.1534 | 0.0411 | 0.0404 | 0.1503 | 0.1136 |
| **59** | Ile | Cd1 | 0.0336 | 0.0488 | 0.1489 | 0.1555 | 0.2307 | 0.2482 |
| **59** | Ile | Cg1 | 0.2932 | 0.0951 | 0.0916 | 0.1320 | 0.2996 | 0.3500 |
| **59** | Ile | Cg2 | 0.2255 | 0.0462 | 0.2799 | 0.0474 | 0.0593 | 0.1379 |
| **59** | Ile | N | 0.0021 | 0.0362 | 0.0394 | 0.0105 | 0.0651 | 0.0483 |
| **60** | Leu | C | 0.0042 | 0.0052 | 0.1821 | 0.9301 | 0.6167 | 0.8320 |
| **60** | Leu | Ca | 0.3893 | 0.1336 | 0.0113 | 0.2965 | 0.1333 | 0.2969 |
| **60** | Leu | Cb | 0.5032 | 0.1242 | 0.6095 | 0.1868 | 0.2066 | 0.2850 |
| **60** | Leu | Cg | 0.7208 | 0.1316 | 0.0962 | | 0.1399 | 0.2078 |
| **60** | Leu | N | 0.4421 | 0.2069 | 0.2028 | 0.0003 | 0.0078 | 0.0020 |
| **61** | Cys | C | 0.1622 | 1.3771 | 0.0910 | | | 0.0825 |
| **61** | Cys | Ca | 0.0969 | 0.1232 | 0.0322 | 0.1710 | 0.2530 | 0.3422 |
| **61** | Cys | Cb | 0.2042 | 1.4452 | 0.0980 | 0.4864 | 0.0788 | 0.0216 |
| **61** | Cys | N | 0.2759 | 0.0044 | 0.2677 | 0.0989 | 0.0599 | 0.0506 |
| **62** | Trp | C | 0.0399 | 0.0106 | 0.0106 | 0.0090 | 0.0954 | 0.0047 |
| **62** | Trp | Ca | 0.0370 | 0.0731 | 0.0075 | 0.0434 | 0.1843 | 0.0560 |
| **62** | Trp | Cb | 0.0063 | 0.1586 | 0.1366 | 0.0590 | 0.1969 | 0.1168 |
| **62** | Trp | Cd1 | | 0.0239 | 0.2136 | 0.1573 | | |
| **62** | Trp | Cg | | 0.1817 | 0.0001 | 0.0301 | | |
| **62** | Trp | N | 0.0399 | 0.0176 | 0.1130 | 0.0003 | 0.0232 | 0.0062 |
| **63** | Gly | C | 0.2787 | 0.0151 | 0.3177 | 0.0284 | 0.0339 | 0.0433 |
| **63** | Gly | Ca | 0.0163 | 0.0034 | 0.0650 | 0.0287 | 0.0095 | 0.1097 |
| **63** | Gly | N | 0.0653 | 0.0207 | 0.0671 | 0.0599 | 0.0825 | 0.0599 |
| **64** | Glu | C | 0.1447 | 0.0048 | 0.1331 | 0.1285 | 0.3415 | 0.2916 |
| **64** | Glu | Ca | 0.0752 | 0.0820 | 0.1551 | 0.2202 | 0.0482 | 0.0474 |
| **64** | Glu | Cb | 0.4746 | 0.0860 | 0.4163 | | 0.0509 | 0.0946 |
| **64** | Glu | Cd | 0.5201 | 0.1479 | 0.6100 | 0.7093 | 0.2559 | 0.3005 |
| **64** | Glu | Cg | 0.8419 | 0.1530 | 0.9727 | 0.5178 | 0.3786 | 0.6788 |
| **64** | Glu | N | 0.1712 | 0.0135 | 0.2189 | 0.3318 | 0.2012 | 0.1850 |
| **65** | Leu | C | 0.0309 | 0.0274 | 0.0443 | 0.0486 | 0.1547 | 0.0021 |
| **65** | Leu | Ca | 0.0268 | 0.0023 | 0.0695 | 0.0542 | 0.1434 | 0.0272 |
| **65** | Leu | Cb | 0.0851 | 0.0190 | 0.0714 | 0.3703 | 0.0146 | 0.0892 |
| **65** | Leu | Cda | 0.2406 | 0.1103 | 0.1050 | 0.5248 | 0.1477 | 0.2197 |
| **65** | Leu | Cdb | 0.1135 | 0.0414 | | | | |
| **65** | Leu | Cg | 0.0237 | 0.0389 | 0.0062 | 0.1883 | 0.1278 | 0.0529 |
| **65** | Leu | N | 0.0592 | 0.0304 | 0.0068 | 0.1762 | 0.0051 | 0.1204 |
| **66** | Met | C | 0.0701 | 0.1213 | 0.1568 | 0.0027 | 0.0628 | 0.0903 |
| **66** | Met | Ca | 0.0788 | 0.1573 | 0.0462 | 0.1142 | 0.0197 | 0.0130 |
| **66** | Met | Cb | 0.1539 | 0.1263 | 0.0638 | 0.1159 | 0.0193 | 0.0413 |
| **66** | Met | Cg | 0.0742 | 0.1081 | 0.0449 | 0.0640 | 0.0032 | 0.0060 |
| **66** | Met | N | 0.0506 | 0.0368 | 0.0512 | 0.0943 | 0.1049 | 0.1142 |
| **67** | Thr | C | 0.0339 | 0.0148 | 0.1371 | 0.0929 | 0.0193 | 0.0188 |
| **67** | Thr | Ca | 0.0952 | 0.0653 | 0.0503 | 0.0855 | 0.0380 | 0.0098 |
| **67** | Thr | Cb | 0.1039 | 0.0155 | 0.0514 | 0.0620 | 0.0064 | 0.0431 |
| **67** | Thr | Cg2 | 0.0144 | 0.0149 | 0.1382 | 0.1217 | 0.0238 | 0.0264 |
| **67** | Thr | N | 0.0030 | 0.0073 | 0.1458 | 0.0718 | 0.1608 | 0.0186 |
| **68** | Leu | C | 0.0276 | 0.0523 | 0.0422 | 0.0094 | 0.1258 | 0.1081 |
| **68** | Leu | Ca | 0.0657 | 0.0002 | 0.1265 | 0.0937 | 0.0121 | 0.0438 |
| **68** | Leu | Cb | 0.0917 | 0.1232 | 0.2539 | 0.2524 | 0.2730 | 0.2710 |
| **68** | Leu | Cda | 0.1532 | 0.1399 | | 0.0890 | 0.0751 | 0.4738 |
| **68** | Leu | Cdb | 0.1309 | 0.0038 | 0.6249 | 0.5300 | 0.1897 | 0.1898 |
| **68** | Leu | Cg | 0.0807 | 0.0550 | 0.2417 | | 0.1484 | 0.1155 |
| **68** | Leu | N | 0.0880 | 0.1089 | 0.1098 | 0.0005 | 0.0385 | 0.0353 |
| **69** | Ala | C | 0.1382 | 0.0852 | 0.2773 | 0.1580 | 0.1203 | 0.1717 |

| | | | | | | | | |
|---|---|---|---|---|---|---|---|---|
| **69** | Ala | Ca | 0.0505 | 0.0369 | 0.0693 | 0.0979 | 0.0101 | 0.0245 |
| **69** | Ala | Cb | 0.0213 | 0.0666 | 0.0072 | 0.0564 | 0.0389 | 0.0641 |
| **69** | Ala | N | 0.0398 | 0.0253 | 0.0419 | 0.0038 | 0.0503 | 0.0330 |
| **70** | Thr | C | 0.0249 | 0.0197 | 0.0004 | 0.0175 | 0.0331 | 0.0470 |
| **70** | Thr | Ca | 0.0151 | 0.0224 | 0.0015 | 0.0158 | 0.0222 | 0.0231 |
| **70** | Thr | Cb | 0.0164 | 0.0067 | 0.0161 | 0.0004 | 0.0191 | 0.1109 |
| **70** | Thr | Cg2 | 0.0075 | 0.0099 | 0.1204 | 0.0819 | 0.0227 | 0.0371 |
| **70** | Thr | N | 0.0317 | 0.0396 | 0.0028 | 0.0371 | 0.0644 | 0.0151 |
| **71** | Trp | C | 0.0558 | 0.0103 | 0.0641 | 0.0405 | 0.0408 | 0.0501 |
| **71** | Trp | Ca | 0.0408 | 0.0781 | 0.0613 | 0.0120 | 0.0437 | 0.0172 |
| **71** | Trp | Cb | 0.0764 | 0.2719 | 0.0259 | 0.0520 | 0.0961 | 0.0614 |
| **71** | Trp | Cd2 | | | 0.0056 | 0.0056 | 0.0309 | 0.0673 |
| **71** | Trp | Cg | | 0.0170 | 0.0055 | 0.0711 | 0.1653 | 0.1666 |
| **71** | Trp | N | 0.0154 | 0.0654 | 0.0172 | 0.0145 | 0.0380 | 0.0095 |
| **72** | Val | C | 0.0434 | 0.0306 | 0.0373 | 0.0177 | | 0.1106 |
| **72** | Val | Ca | 0.0356 | 0.1195 | 0.1774 | 0.0803 | 0.0824 | 0.0578 |
| **72** | Val | Cb | 0.0649 | 0.1154 | 0.6134 | 0.6152 | 0.0653 | 0.2260 |
| **72** | Val | Cga | | 0.2616 | | 0.6228 | | |
| **72** | Val | N | 0.1956 | 0.1866 | 0.0271 | 0.0186 | 0.0360 | 0.0346 |
| **73** | Gly | C | 0.0907 | 0.0026 | 0.0921 | 0.0583 | 0.0625 | 0.0367 |
| **73** | Gly | Ca | 0.0663 | 0.1174 | 0.1458 | 0.0970 | 0.0470 | 0.0384 |
| **73** | Gly | N | 0.0637 | 0.0025 | 0.0829 | 0.0715 | 0.0851 | 0.0032 |
| **74** | Val | C | 0.0323 | 0.0527 | 0.0761 | 0.0047 | 0.0129 | 0.0129 |
| **74** | Val | Ca | 0.0144 | 0.0266 | 0.0093 | 0.0175 | 0.0798 | 0.0807 |
| **74** | Val | Cb | 0.0104 | 0.0887 | 0.0542 | 0.0342 | 0.0267 | 0.0068 |
| **74** | Val | Cga | 0.0382 | 0.0838 | | 0.0054 | | |
| **74** | Val | N | 0.0360 | 0.0254 | 0.0491 | 0.0160 | 0.0198 | 0.0119 |
| **75** | Asn | C | 0.0219 | 0.0492 | 0.0101 | 0.0375 | 0.0759 | 0.0053 |
| **75** | Asn | Ca | 0.0366 | 0.0567 | 0.0752 | 0.0630 | 0.0004 | 0.0083 |
| **75** | Asn | Cb | 0.0740 | 0.0387 | 0.0060 | 0.0030 | 0.0397 | 0.0436 |
| **75** | Asn | Cg | 0.0529 | 0.0547 | 0.0838 | 0.0440 | 0.0039 | 0.0065 |
| **75** | Asn | N | 0.0297 | 0.0546 | 0.0828 | 0.0449 | 0.0151 | 0.0441 |
| **76** | Leu | C | 0.0129 | 0.0015 | 0.0847 | 0.0651 | 0.0432 | 0.0413 |
| **76** | Leu | Ca | 0.0423 | 0.0385 | 0.0995 | 0.1711 | 0.0454 | 0.0427 |
| **76** | Leu | Cb | 0.0214 | 0.0958 | 0.1312 | 0.0299 | 0.1066 | 0.0757 |
| **76** | Leu | Cda | 0.0346 | 0.1176 | | 0.0926 | | |
| **76** | Leu | Cg | 0.0003 | 0.0065 | 0.2880 | 0.1400 | 0.0737 | 0.0586 |
| **76** | Leu | N | 0.0005 | 0.0565 | 0.0579 | 0.0517 | 0.0131 | 0.0328 |
| **77** | Glu | C | 0.0175 | 0.0664 | 0.0515 | 0.0453 | 0.1677 | 0.0019 |
| **77** | Glu | Ca | 0.0429 | 0.0308 | 0.0455 | 0.0154 | 0.1122 | 0.0425 |
| **77** | Glu | Cb | 0.0987 | 0.0328 | 0.1041 | 0.1213 | 0.0108 | 0.0177 |
| **77** | Glu | Cd | | 0.0395 | | 0.0257 | | 0.0782 |
| **77** | Glu | Cg | | 0.0057 | 0.0394 | 0.0434 | 0.0203 | 0.1052 |
| **77** | Glu | N | 0.0659 | 0.1368 | 0.1687 | 0.0558 | 0.0167 | 0.1098 |
| **78** | Asp | C | 0.0348 | 0.0432 | 0.0272 | 0.0120 | 0.0566 | 0.0339 |
| **78** | Asp | Ca | 0.1439 | 0.0664 | 0.1614 | 0.1536 | 0.0440 | 0.0108 |
| **78** | Asp | Cb | 0.0469 | 0.0648 | 0.0502 | 0.0384 | 0.0045 | 0.1111 |
| **78** | Asp | Cg | 0.0518 | 0.0255 | 0.1701 | 0.0551 | 0.0082 | 0.0044 |
| **78** | Asp | N | 0.0609 | 0.0679 | 0.2141 | 0.1298 | 0.0421 | 0.0057 |
| **79** | Pro | C | | | | | 0.1619 | |
| **79** | Pro | Ca | 0.0108 | 0.0576 | 0.0171 | 0.0996 | 0.0524 | 0.0402 |
| **79** | Pro | Cb | | | | | 0.0302 | |
| **79** | Pro | Cd | 0.0009 | 0.0355 | 0.0077 | 0.0500 | 0.0336 | 0.0735 |
| **79** | Pro | Cg | | | | | 0.0078 | |
| **79** | Pro | N | 0.0021 | 0.0325 | 0.0894 | 0.1028 | 0.0005 | 0.0125 |
| **80** | Ala | C | 0.0766 | 0.0407 | 0.1414 | 0.0582 | 0.0186 | 0.1744 |
| **80** | Ala | Ca | 0.0143 | 0.0557 | 0.0055 | 0.0394 | 0.0233 | 0.1069 |
| **80** | Ala | Cb | 0.0511 | 0.0645 | 0.0002 | 0.0157 | 0.0529 | 0.1798 |

| | | | | | | | | |
|---|---|---|---|---|---|---|---|---|
| **80** | Ala | N | 0.0096 | 0.0470 | 0.0021 | 0.0115 | 0.0175 | 0.0247 |
| **81** | Ser | C | 0.0869 | 0.0806 | 0.0707 | 0.0510 | 0.1055 | 0.1004 |
| **81** | Ser | Ca | 0.0841 | 0.0989 | 0.0232 | 0.0275 | 0.1251 | 0.1529 |
| **81** | Ser | N | 0.0211 | 0.0186 | 0.0277 | 0.0207 | 0.0864 | 0.0771 |
| **82** | Arg | C | 0.0854 | 0.0295 | 0.0001 | 0.0049 | 0.0320 | 0.0068 |
| **82** | Arg | Ca | 0.0030 | 0.0032 | 0.0153 | 0.0464 | 0.0344 | 0.0368 |
| **82** | Arg | Cb | 0.0762 | 0.0076 | 0.1129 | 0.0921 | 0.0579 | 0.0708 |
| **82** | Arg | Cd | 0.1204 | 0.0958 | 0.0001 | 0.0426 | 0.1322 | 0.0828 |
| **82** | Arg | Cg | 0.0534 | 0.0709 | 0.0005 | 0.0736 | 0.0636 | 0.0360 |
| **82** | Arg | Cz | | | 0.0005 | 0.0007 | | |
| **82** | Arg | N | 0.0424 | 0.0469 | 0.0127 | 0.0136 | 0.0984 | 0.0142 |
| **83** | Asp | C | | 0.0270 | 0.1002 | 0.0687 | 0.0258 | 0.0768 |
| **83** | Asp | Ca | 0.0322 | 0.0303 | 0.0109 | 0.0593 | 0.0538 | 0.0006 |
| **83** | Asp | Cb | | 0.1110 | 0.0617 | 0.0623 | 0.0013 | 0.0425 |
| **83** | Asp | N | | 0.0200 | 0.0198 | 0.0389 | 0.0957 | 0.0230 |
| **84** | Leu | C | 0.1540 | 0.0236 | 0.1520 | 0.0539 | 0.0140 | 0.0146 |
| **84** | Leu | Ca | 0.0068 | 0.0172 | 0.0504 | 0.0538 | 0.0042 | 0.0063 |
| **84** | Leu | Cb | 0.0566 | 0.0254 | 0.1557 | 0.0935 | 0.0445 | 0.1440 |
| **84** | Leu | Cg | 0.0817 | 0.0854 | 0.0557 | 0.0741 | 0.0864 | 0.1010 |
| **84** | Leu | N | 0.0379 | 0.0260 | 0.0116 | 0.0017 | 0.0347 | 0.0345 |
| **85** | Val | C | 0.0922 | 0.0779 | 0.0026 | 0.0504 | 0.0242 | 0.0340 |
| **85** | Val | Ca | 0.0117 | 0.0163 | 0.0597 | 0.0095 | 0.0639 | 0.0496 |
| **85** | Val | Cb | 0.0527 | 0.0938 | 0.0537 | 0.0557 | 0.0270 | 0.0080 |
| **85** | Val | Cga | 0.1206 | 0.0314 | 0.2176 | 0.1431 | 0.0083 | 0.0668 |
| **85** | Val | Cgb | 0.0951 | 0.0039 | 0.1973 | 0.1653 | 0.0989 | 0.1421 |
| **85** | Val | N | 0.0475 | 0.0519 | 0.0479 | 0.0284 | 0.0040 | 0.0025 |
| **86** | Val | C | 0.1056 | 0.1027 | 0.0956 | 0.0611 | 0.0379 | 0.0759 |
| **86** | Val | Ca | 0.0891 | 0.0726 | 0.1770 | 0.1375 | 0.0361 | 0.0558 |
| **86** | Val | Cb | 0.0042 | 0.0219 | 0.0251 | 0.0057 | 0.0624 | 0.0220 |
| **86** | Val | Cga | 0.0198 | 0.0117 | 0.0881 | 0.0010 | 0.0402 | 0.0577 |
| **86** | Val | Cgb | 0.0035 | 0.0140 | 0.0403 | 0.0273 | 0.0943 | 0.0877 |
| **86** | Val | N | 0.0405 | 0.0274 | 0.1044 | 0.0347 | 0.0499 | 0.0161 |
| **87** | Ser | C | 0.0359 | 0.0571 | 0.0257 | 0.1925 | 0.0078 | 0.0382 |
| **87** | Ser | Ca | 0.0221 | 0.0530 | 0.0564 | 0.0211 | 0.0218 | 0.0145 |
| **87** | Ser | Cb | 0.0415 | 0.0590 | 0.0100 | 0.0227 | 0.0189 | 0.0037 |
| **87** | Ser | N | 0.0016 | 0.0143 | 0.0137 | 0.0494 | 0.0438 | 0.0098 |
| **88** | Tyr | C | 0.1046 | 0.0373 | 0.0263 | 0.0029 | 0.1044 | 0.0983 |
| **88** | Tyr | Ca | 0.0337 | 0.0262 | 0.0433 | 0.0114 | 0.0166 | 0.0276 |
| **88** | Tyr | Cb | 0.1686 | 0.1227 | 0.1147 | 0.1402 | 0.0971 | 0.1016 |
| **88** | Tyr | Cd | | | | 0.0000 | | 0.6090 |
| **88** | Tyr | Cg | 0.4502 | 0.0354 | | 0.0000 | | 0.0930 |
| **88** | Tyr | N | 0.0014 | 0.0207 | 0.0451 | 0.0201 | 0.0321 | 0.0371 |
| **89** | Val | C | 0.0721 | 0.0621 | 0.0443 | 0.0973 | 0.0301 | 0.0413 |
| **89** | Val | Ca | 0.1289 | 0.1774 | 0.0726 | 0.0918 | 0.1363 | 0.0601 |
| **89** | Val | Cb | 0.0018 | 0.0186 | 0.0020 | 0.0556 | 0.0705 | 0.0055 |
| **89** | Val | Cga | 0.1262 | 0.0003 | 0.1510 | 0.0806 | 0.1035 | 0.0884 |
| **89** | Val | N | 0.0898 | 0.1854 | 0.0454 | 0.0092 | 0.0471 | 0.0138 |
| **90** | Asn | C | 0.0407 | 0.1069 | 0.0293 | 0.1195 | 0.0529 | 0.0712 |
| **90** | Asn | Ca | 0.0974 | 0.0757 | 0.0052 | 0.0783 | 0.0117 | 0.0335 |
| **90** | Asn | Cb | 0.1196 | 0.0919 | 0.0166 | 0.0119 | 0.0670 | 0.1035 |
| **90** | Asn | Cg | 0.0160 | 0.0180 | 0.1433 | 0.0066 | 0.0634 | 0.0357 |
| **90** | Asn | N | 0.1228 | 0.0916 | 0.0885 | 0.1007 | 0.1266 | 0.0512 |
| **91** | Thr | C | 0.0731 | 0.0531 | 0.0309 | 0.0143 | 0.0223 | 0.0355 |
| **91** | Thr | Ca | 0.0153 | 0.0149 | 0.1054 | 0.0157 | 0.1028 | 0.0720 |
| **91** | Thr | Cb | 0.0266 | 0.0524 | 0.1229 | 0.0017 | 0.0052 | 0.0244 |
| **91** | Thr | Cg2 | 0.0009 | 0.0743 | 0.0324 | 0.0297 | 0.0013 | 0.0291 |
| **91** | Thr | N | 0.0482 | 0.0150 | 0.1536 | 0.1230 | 0.0027 | 0.0556 |
| **92** | Asn | C | 0.1129 | 0.0820 | 0.1731 | 0.2017 | 0.0968 | 0.2121 |

| | | | | | | | | |
|---|---|---|---|---|---|---|---|---|
| **92** | Asn | Ca | 0.0868 | 0.0916 | 0.2599 | 0.0794 | 0.0298 | 0.0269 |
| **92** | Asn | Cb | 0.1178 | 0.0638 | 0.1283 | 0.1052 | 0.1451 | 0.1448 |
| **92** | Asn | Cg | 0.0838 | 0.0810 | 0.1317 | 0.0267 | 0.0019 | 0.0114 |
| **92** | Asn | N | 0.0203 | 0.0064 | 0.2231 | 0.1869 | 0.0239 | 0.0374 |
| **93** | Met | C | 0.0536 | 0.0554 | 0.1846 | 0.1398 | 0.1309 | 0.2273 |
| **93** | Met | Ca | 0.0095 | 0.0275 | 0.0845 | 0.0530 | 0.0880 | 0.0762 |
| **93** | Met | Cb | 0.2482 | 0.1580 | 0.3732 | 0.6428 | 0.3748 | 0.2711 |
| **93** | Met | Cg | 0.1269 | 0.0352 | 0.1312 | 0.1508 | 0.0479 | 0.0847 |
| **93** | Met | N | 0.1746 | 0.1172 | 0.2869 | 0.1578 | 0.1097 | 0.1317 |
| **94** | Gly | C | 0.0325 | 0.1398 | 0.1154 | 0.1071 | 0.0714 | 0.0473 |
| **94** | Gly | Ca | 0.2658 | 0.0238 | 0.0262 | 0.1090 | 0.1970 | 0.1147 |
| **94** | Gly | N | 0.0217 | 0.0181 | 0.1093 | 0.1152 | 0.0398 | 0.0241 |
| **95** | Leu | C | 0.1176 | 0.0093 | 0.2735 | 0.0640 | 0.0512 | 0.0562 |
| **95** | Leu | Ca | 0.0216 | 0.0044 | 0.3215 | 0.1630 | 0.0059 | 0.0099 |
| **95** | Leu | Cb | 0.0038 | 0.1039 | 0.1065 | 0.1509 | 0.0010 | 0.0217 |
| **95** | Leu | Cda | 0.6264 | 0.4986 | 0.3203 | 0.3000 | 0.4391 | 0.5646 |
| **95** | Leu | Cdb | 0.0651 | 0.0615 | 0.6455 | 0.0188 | 0.0740 | 0.0387 |
| **95** | Leu | Cg | 0.1669 | 0.1763 | 0.0665 | 0.0528 | 0.1194 | 0.1788 |
| **95** | Leu | N | 0.1204 | 0.0152 | 0.1628 | 0.0888 | 0.0208 | 0.0258 |
| **98** | Arg | C | | 0.2823 | 0.1006 | | 0.2031 | 0.1021 |
| **98** | Arg | Ca | 0.3941 | 0.4243 | 0.1330 | | 0.2873 | 0.2831 |
| **98** | Arg | Cb | 0.0677 | 0.1517 | 0.1587 | | 0.1400 | 0.0012 |
| **98** | Arg | Cd | 0.2285 | 0.1039 | | | 0.1546 | 0.1124 |
| **98** | Arg | Cg | 0.2455 | 0.0918 | 0.2418 | | 0.1792 | 0.1071 |
| **98** | Arg | Cz | | 0.0646 | 0.0003 | | | |
| **98** | Arg | N | 0.0808 | 0.0838 | 0.0067 | | 0.1032 | 0.1040 |
| **99** | Gln | C | 0.1314 | 0.1083 | 0.1805 | 0.0236 | 0.0179 | 0.1083 |
| **99** | Gln | Ca | 0.3260 | 0.3049 | 0.0374 | 0.0189 | 0.1458 | 0.1532 |
| **99** | Gln | Cb | 0.3985 | 0.3964 | 0.0773 | 0.0063 | 0.4086 | 0.3964 |
| **99** | Gln | Cd | 0.0952 | 0.2149 | 0.0428 | 0.4690 | 0.4553 | 0.1574 |
| **99** | Gln | Cg | 0.0459 | 0.2148 | 0.2986 | 0.3774 | 0.0445 | 0.1918 |
| **99** | Gln | N | 0.0809 | 0.1055 | 0.0301 | 0.0339 | 0.2289 | 0.1011 |
| **100** | Leu | C | 0.0228 | 0.2158 | 0.2572 | 0.0226 | 0.0624 | 0.0430 |
| **100** | Leu | Ca | 0.2917 | 0.0110 | 0.7101 | 0.1820 | 0.2010 | 0.1923 |
| **100** | Leu | Cb | 0.2235 | 0.1965 | 0.6744 | 0.0885 | 0.1279 | 0.1714 |
| **100** | Leu | N | 0.0369 | 0.1427 | 0.0659 | 0.0523 | 0.2301 | 0.2439 |
| **101** | Leu | C | | 0.0139 | 0.2369 | 0.1691 | 0.1330 | 0.2099 |
| **101** | Leu | Ca | 0.0213 | 0.0092 | 0.1852 | 0.0717 | 0.0975 | 0.0132 |
| **101** | Leu | Cb | 0.2154 | 0.3626 | 0.0557 | 0.1022 | 0.0609 | 0.0536 |
| **101** | Leu | Cda | 0.5879 | 0.7545 | 0.1851 | | 0.2138 | 0.1931 |
| **101** | Leu | Cdb | 0.4777 | 0.3200 | 0.8518 | 0.8142 | 0.9338 | 0.2743 |
| **101** | Leu | Cg | 0.0850 | 0.1161 | 0.1808 | 0.1762 | 0.0858 | 0.1393 |
| **101** | Leu | N | 0.0610 | 0.1544 | 0.0700 | 0.0021 | 0.0093 | 0.0560 |
| **102** | Trp | C | 0.5743 | 0.7806 | 0.7106 | 0.5889 | 0.4205 | 0.7990 |
| **102** | Trp | Ca | 0.4724 | 0.0718 | 0.2260 | 0.7308 | 0.5177 | 0.6040 |
| **102** | Trp | Cb | 0.3022 | 0.3022 | 0.4064 | 0.3628 | 0.0722 | 0.1643 |
| **102** | Trp | Cd1 | 0.9401 | 0.6392 | 0.1937 | 0.7204 | 0.0674 | 1.1703 |
| **102** | Trp | Cd2 | | 0.5894 | | 0.1390 | 0.0041 | |
| **102** | Trp | Cg | 1.1846 | 0.2542 | 0.3303 | 0.2346 | 0.0780 | 0.0392 |
| **102** | Trp | N | 0.3735 | 0.1271 | 0.1243 | 0.0444 | 0.0082 | 0.1567 |
| **103** | Phe | C | 0.3867 | 0.0323 | 0.1262 | 0.2566 | 0.1139 | 0.0882 |
| **103** | Phe | Ca | 0.2236 | 0.2963 | 0.2065 | 0.1425 | 0.0911 | 0.1166 |
| **103** | Phe | Cb | 0.0237 | 0.2331 | 0.0955 | 0.2532 | 0.2908 | 0.0438 |
| **103** | Phe | N | 0.6091 | 0.8209 | 0.2524 | 0.2688 | 0.3952 | 0.2074 |
| **104** | His | C | 0.4075 | 0.2588 | 0.0031 | 0.0309 | 0.0909 | 0.1311 |
| **104** | His | Ca | 0.0536 | 0.0299 | 0.1790 | 0.1819 | 0.0751 | 0.0838 |
| **104** | His | Cb | 0.1896 | 0.3221 | 0.0549 | 0.0344 | 0.1027 | 0.0493 |
| **104** | His | Cd2 | 0.0601 | 0.0890 | 0.2385 | 0.2261 | 0.1972 | 0.3588 |

| | | | | | | | | |
|---|---|---|---|---|---|---|---|---|
| **104** | His | Ce1 | 0.1870 | 0.0433 | 0.3587 | 0.2707 | 0.0392 | 0.2212 |
| **104** | His | N | 0.3455 | 0.2826 | 0.0981 | 0.0134 | 0.0590 | 0.1497 |
| **105** | Ile | C | 0.2599 | 0.0452 | 0.9080 | 0.9559 | 0.8650 | 0.7176 |
| **105** | Ile | Ca | 0.2687 | 0.1831 | 0.0019 | 0.1421 | 0.0556 | 0.0138 |
| **105** | Ile | Cb | 0.6239 | 0.6066 | 0.1769 | 0.1501 | 0.4439 | 0.4282 |
| **105** | Ile | Cd1 | 0.2510 | 0.4130 | 0.0762 | 0.0176 | 0.1043 | 0.0364 |
| **105** | Ile | Cg1 | | 0.3022 | 0.1353 | 0.0288 | 0.0764 | 0.1216 |
| **105** | Ile | Cg2 | 1.0932 | 1.7399 | 0.4801 | 0.7306 | 0.3874 | 1.0159 |
| **105** | Ile | N | 0.1984 | 0.0830 | 0.0680 | 0.2157 | 0.0244 | 0.1846 |
| **106** | Ser | C | 1.7398 | 1.1771 | 0.1232 | 0.1915 | 0.1522 | 0.6413 |
| **106** | Ser | Ca | 0.2310 | 1.2280 | 0.4049 | 0.6714 | 0.4732 | 0.6091 |
| **106** | Ser | Cb | 1.0129 | 0.0873 | 0.0797 | 0.0286 | 0.0164 | 0.7308 |
| **106** | Ser | N | 0.0205 | 0.5057 | 1.0293 | 0.8279 | 0.8438 | 0.9135 |
| **107** | Cys | C | 0.4454 | 0.3964 | 0.2621 | 0.3198 | 0.2244 | 0.0528 |
| **107** | Cys | Ca | 0.4808 | 0.2441 | 0.4386 | 0.3611 | 0.5525 | 0.5665 |
| **107** | Cys | Cb | 0.3321 | 0.1273 | 0.2731 | 0.1608 | 0.1623 | 0.0492 |
| **107** | Cys | N | 0.5134 | 0.2639 | 0.2837 | 0.2458 | 0.4634 | 0.4155 |
| **108** | Leu | C | 0.8174 | 0.6556 | 0.5563 | 0.6232 | 0.6655 | 0.6415 |
| **108** | Leu | Ca | 0.0047 | 0.0140 | 0.2563 | 0.2592 | 0.1802 | 0.2292 |
| **108** | Leu | Cb | 0.1328 | 0.0077 | 0.2318 | 0.3215 | 0.2018 | 0.0396 |
| **108** | Leu | Cda | 0.3678 | 0.3057 | | 0.3462 | 0.3061 | 0.2460 |
| **108** | Leu | Cg | 0.2648 | 0.2707 | 0.0938 | 0.2390 | 0.0979 | 0.1416 |
| **108** | Leu | N | 0.6154 | 0.5344 | 0.6686 | 0.5992 | 0.5779 | 0.5560 |
| **109** | Thr | C | 0.1562 | 0.3004 | 0.5465 | | 0.4641 | 0.4906 |
| **109** | Thr | Ca | 0.1930 | 0.1300 | 0.1869 | 0.1021 | 0.1411 | 0.0670 |
| **109** | Thr | Cb | | 0.1235 | 0.1653 | 0.0255 | 0.0562 | 0.0423 |
| **109** | Thr | Cg2 | | 0.3149 | 0.5024 | 0.8309 | 0.9107 | 1.1000 |
| **109** | Thr | N | 0.1892 | 0.2062 | 0.0604 | 0.2175 | 0.1928 | 0.2499 |
| **110** | Phe | C | 0.1713 | 0.0221 | 0.3222 | 0.6085 | 0.8447 | |
| **110** | Phe | Ca | 0.2410 | 0.0105 | 1.9259 | 0.9901 | 0.8245 | 0.7927 |
| **110** | Phe | Cb | 0.9852 | 0.8988 | 0.1916 | 0.0635 | 0.4822 | 0.3879 |
| **110** | Phe | Cg | | 0.3255 | | | | |
| **110** | Phe | N | 0.1290 | 0.1281 | 0.1721 | 0.1091 | 0.1534 | 0.1661 |
| **111** | Gly | C | 0.6125 | 0.4053 | 0.2329 | 0.3888 | 0.2515 | 0.1850 |
| **111** | Gly | Ca | 0.1384 | 0.0538 | 0.0528 | 0.0569 | 0.0227 | 0.0635 |
| **111** | Gly | N | 0.4198 | 0.1748 | 0.4988 | 0.5251 | 0.5188 | 0.3206 |
| **112** | Arg | C | 0.6206 | | 0.0612 | 0.0654 | 0.0177 | 0.0050 |
| **112** | Arg | Ca | 0.2728 | | 0.0819 | 0.1090 | 0.1224 | 0.0344 |
| **112** | Arg | Cb | 0.9951 | | 0.1674 | 0.1186 | 0.0958 | 0.0817 |
| **112** | Arg | Cd | 0.0613 | | | 1.5526 | 1.3045 | 1.2881 |
| **112** | Arg | Cg | | | 0.6166 | | 0.5691 | 0.3816 |
| **112** | Arg | Cz | | | 0.0000 | 0.0239 | | 0.1240 |
| **112** | Arg | N | 1.8052 | | 0.2946 | 0.3436 | 0.2881 | 0.2194 |
| **113** | Glu | C | 0.3335 | 0.1744 | 0.0346 | 0.0086 | 0.1799 | 0.1153 |
| **113** | Glu | Ca | 0.1648 | 0.0160 | 0.0631 | 0.0759 | 0.0998 | 0.0739 |
| **113** | Glu | Cb | 0.0162 | 0.1949 | 0.1450 | 0.1891 | 0.1629 | 0.1576 |
| **113** | Glu | Cd | 0.4471 | 0.4119 | 0.2422 | 0.3171 | 0.1939 | 0.1886 |
| **113** | Glu | Cg | | 0.1254 | 0.1177 | 0.0701 | 0.0816 | 0.0688 |
| **113** | Glu | N | 0.8265 | 0.4001 | 0.0352 | 0.0434 | 0.0089 | 0.0357 |
| **114** | Thr | C | 0.1734 | 0.8485 | 0.0070 | 0.1191 | 0.0997 | 0.0431 |
| **114** | Thr | Ca | 0.4204 | 0.4269 | 0.0854 | 0.0439 | 0.0419 | 0.0410 |
| **114** | Thr | Cb | 0.1979 | 0.1121 | 0.0929 | 0.0735 | 0.1967 | 0.1429 |
| **114** | Thr | Cg2 | 0.4360 | 0.0479 | 0.0645 | 0.1360 | 0.0201 | 0.0102 |
| **114** | Thr | N | 0.0370 | 0.0239 | 0.0059 | 0.0625 | 0.0763 | 0.0088 |
| **115** | Val | C | 0.4536 | 0.1511 | 0.2522 | 0.2788 | 0.4065 | 0.3335 |
| **115** | Val | Ca | 0.6835 | 0.1262 | 0.0802 | 0.0928 | 0.2454 | 0.2434 |
| **115** | Val | Cb | 0.6185 | 0.3073 | 0.2034 | 0.1218 | 0.1287 | 0.0974 |
| **115** | Val | Cga | 0.4606 | 0.1532 | 0.0368 | 0.0196 | 0.0063 | 0.0002 |

| | | | | | | | | |
|---|---|---|---|---|---|---|---|---|
| **115** | Val | Cgb | 0.2965 | 0.3629 | 0.7855 | 0.6275 | 0.7437 | 0.2926 |
| **115** | Val | N | 0.1387 | 0.6126 | 0.2022 | 0.1812 | 0.3660 | 0.2698 |
| **116** | Ile | C | 0.0590 | 0.0500 | 0.0802 | 0.3087 | 0.6253 | 0.1428 |
| **116** | Ile | Ca | 0.5084 | 0.4399 | 0.0006 | 0.0023 | 0.2240 | 0.1912 |
| **116** | Ile | Cb | 1.2469 | 1.3413 | 0.5261 | | 0.7814 | 0.6508 |
| **116** | Ile | Cd1 | 1.3086 | 1.4439 | | | 1.0047 | 0.7167 |
| **116** | Ile | Cg1 | 1.2397 | 1.2985 | 0.4041 | | 0.5906 | 0.1581 |
| **116** | Ile | Cg2 | 0.2873 | 0.1188 | 0.0743 | 0.0092 | 0.0300 | 0.0675 |
| **116** | Ile | N | 0.3449 | 0.3469 | 0.1996 | 0.1156 | 0.2784 | 0.1434 |
| **117** | Glu | C | 0.3404 | 0.0587 | 0.0433 | 0.0935 | 0.4083 | 0.2748 |
| **117** | Glu | Ca | 0.2098 | 0.1875 | 0.1423 | 0.1168 | 0.2045 | 0.1726 |
| **117** | Glu | Cb | 0.1671 | 0.1346 | 0.1963 | 0.0865 | 0.2966 | 0.2210 |
| **117** | Glu | Cd | | | 0.4367 | 0.3157 | 0.1450 | 0.0619 |
| **117** | Glu | N | 0.1933 | 0.1619 | 0.2138 | 0.1566 | 0.0535 | 0.1705 |
| **118** | Tyr | C | 0.3863 | 0.9182 | 0.7998 | 1.0619 | 1.1620 | 1.1863 |
| **118** | Tyr | Ca | 0.1395 | 0.7129 | 0.5322 | 0.5845 | 0.7217 | 0.4951 |
| **118** | Tyr | Cb | 0.7472 | 0.4565 | 0.5703 | | 0.8555 | 0.8529 |
| **118** | Tyr | Cd | 1.2751 | 1.3541 | 1.6374 | 0.0000 | | |
| **118** | Tyr | Ce | | 0.3944 | 0.3484 | 0.0000 | | |
| **118** | Tyr | N | 0.1887 | 0.0099 | 0.2273 | 0.0637 | 0.0398 | 0.1123 |
| **120** | Val | C | 0.1061 | 0.1862 | 0.5213 | 0.3716 | 0.4412 | 0.4102 |
| **120** | Val | Ca | 0.0640 | 0.2685 | 0.1009 | 0.4353 | 0.1550 | 0.3740 |
| **120** | Val | N | 0.3724 | 0.0104 | 0.0978 | 0.1984 | 0.0184 | 0.0670 |
| **121** | Ser | C | 0.6906 | | | 0.3269 | | |
| **121** | Ser | Ca | 0.8458 | | | 0.8367 | | |
| **121** | Ser | N | 0.7627 | | | 0.8094 | | |
| **122** | Phe | C | 1.3841 | 0.8609 | | 1.1262 | 1.8475 | 1.8432 |
| **122** | Phe | Ca | 0.5868 | 0.1548 | | 0.2991 | 0.1687 | 0.1630 |
| **122** | Phe | Cb | 0.1724 | 0.4862 | | | 0.6613 | 0.6694 |
| **122** | Phe | N | 1.3186 | 0.9699 | | 0.2698 | 0.0960 | 0.0827 |
| **123** | Gly | C | 1.6894 | 1.9991 | 1.9655 | 1.6073 | 1.6566 | 1.8079 |
| **123** | Gly | Ca | 0.2483 | 0.1017 | 0.1451 | 0.0672 | 0.0528 | 0.0221 |
| **123** | Gly | N | 0.3326 | 0.1981 | 0.0316 | 0.0092 | 0.1486 | 0.0953 |
| **124** | Val | C | | 0.6407 | 0.1394 | 0.6016 | 0.5106 | 0.0279 |
| **124** | Val | Ca | | 0.6441 | 0.0497 | 0.0455 | 0.3407 | 0.0549 |
| **124** | Val | Cb | | 0.4333 | 0.0868 | 0.0342 | 0.0256 | 0.1002 |
| **124** | Val | Cga | | | 0.0683 | 0.0150 | | 0.1438 |
| **124** | Val | Cgb | | | 0.7837 | 0.1138 | 0.6450 | 0.0726 |
| **124** | Val | N | | 0.1705 | 0.0679 | 0.0133 | 0.0290 | 0.0193 |
| **125** | Trp | C | 0.8489 | 0.9847 | | 0.9334 | 1.1345 | 1.3321 |
| **125** | Trp | Ca | 0.4755 | 0.1440 | | 0.1897 | 0.2328 | 0.2091 |
| **125** | Trp | Cb | | 0.9963 | | | 0.3453 | 0.3347 |
| **125** | Trp | Cd1 | 1.1425 | 0.7315 | | | | |
| **125** | Trp | Cg | | 0.1626 | | | | 0.3412 |
| **125** | Trp | N | 1.0425 | 1.1893 | | 1.9714 | 1.4553 | 1.7726 |
| **127** | Arg | Ca | 0.3036 | 0.1335 | 0.1554 | 0.1443 | 0.3612 | 0.4168 |
| **127** | Arg | N | 0.1283 | 0.2328 | 0.0131 | 0.1884 | 0.1441 | 0.1832 |
| **128** | Thr | C | 0.4246 | 0.1934 | 0.0970 | 0.0998 | 0.0593 | 0.3196 |
| **128** | Thr | Ca | 0.1938 | 0.6015 | 0.1652 | 0.0661 | 0.1859 | 0.1309 |
| **128** | Thr | Cb | 1.9991 | 1.4512 | 0.0196 | 0.1763 | 0.5603 | 0.8161 |
| **128** | Thr | Cg2 | 0.3540 | 0.7120 | 1.0601 | 0.6846 | 0.6529 | 0.2823 |
| **128** | Thr | N | 0.1331 | 0.0626 | 0.1428 | 0.2023 | 0.0327 | 0.0016 |
| **129** | Pro | C | | | | | 0.4678 | |
| **129** | Pro | Ca | 0.1229 | 0.0044 | 0.2497 | 0.0996 | 0.2713 | 0.2231 |
| **129** | Pro | Cb | | | | | 0.2427 | |
| **129** | Pro | Cd | 1.4439 | 1.3062 | 0.6808 | 0.8034 | 0.9775 | 0.8766 |
| **129** | Pro | Cg | | | | | 0.2394 | |
| **129** | Pro | N | 0.7699 | 0.4985 | 0.0973 | 0.0167 | 0.4385 | 0.0242 |

| | | | | | | | | |
|---|---|---|---|---|---|---|---|---|
| **130** | Pro | C | 0.1595 | 0.0163 | 0.2818 | 0.2598 | 0.3174 | 0.4616 |
| **130** | Pro | Ca | | 0.6047 | 0.8891 | 0.3572 | 0.5189 | 0.3524 |
| **130** | Pro | Cb | | 0.1822 | 0.2160 | 0.1742 | 0.1320 | 0.3385 |
| **130** | Pro | Cd | | 0.5014 | 0.2356 | 0.0269 | 0.1999 | 0.2532 |
| **130** | Pro | Cg | | 0.2572 | 0.3743 | | 0.0467 | 0.0382 |
| **130** | Pro | N | | 0.8877 | 0.1744 | 0.0158 | 0.0492 | 0.1142 |
| **131** | Ala | C | 0.9179 | 0.5554 | 0.5611 | 0.7415 | 0.4303 | 0.1598 |
| **131** | Ala | Ca | 0.6378 | 0.5738 | 0.2277 | 0.5030 | 0.1353 | 0.0964 |
| **131** | Ala | Cb | 0.0842 | 0.1282 | 0.3145 | 0.4385 | 0.3253 | 0.2586 |
| **131** | Ala | N | 0.7963 | 0.5207 | 0.3711 | 0.4533 | 0.4034 | 0.3053 |
| **133** | Arg | C | 1.9386 | | | 1.4195 | | |
| **133** | Arg | Ca | 0.6722 | | | 0.5475 | | |
| **133** | Arg | N | 0.1480 | | | 0.1950 | | |
| **135** | Pro | C | 0.0314 | 0.3130 | 0.1865 | 0.3451 | | |
| **135** | Pro | Ca | 0.1552 | 0.2896 | 0.2722 | 0.4515 | | |
| **135** | Pro | Cb | 0.0688 | 0.0223 | 0.1084 | 0.1234 | | |
| **135** | Pro | Cd | 0.1253 | 0.0888 | 0.1440 | 0.1970 | | |
| **135** | Pro | Cg | 0.1247 | 0.0797 | 0.1941 | 0.0829 | | |
| **135** | Pro | N | 0.0183 | 0.0488 | 0.3313 | 0.6021 | | |
| **136** | Asn | C | 0.2364 | 0.0523 | 0.1638 | 0.1418 | 0.1831 | 0.2114 |
| **136** | Asn | Ca | 0.1508 | 0.0947 | 0.0549 | 0.0073 | 0.0578 | 0.0828 |
| **136** | Asn | Cb | 0.7865 | 0.3867 | 0.5498 | 1.3522 | 1.2167 | 1.2026 |
| **136** | Asn | Cg | 0.0520 | 0.4654 | 0.0909 | 0.0756 | 0.0021 | 0.0915 |
| **136** | Asn | N | 0.1470 | 0.3364 | 0.2831 | 0.1474 | 0.1102 | 0.1484 |
| **137** | Ala | C | 0.0616 | 0.1363 | 0.0272 | 0.1911 | 0.4908 | 0.1835 |
| **137** | Ala | Ca | 0.0988 | 0.0409 | 0.3876 | 0.0047 | 0.1312 | 0.1952 |
| **137** | Ala | Cb | 0.2912 | 0.0266 | 0.1128 | 0.5501 | 0.6241 | 0.7467 |
| **137** | Ala | N | 1.0566 | 0.2732 | 0.7163 | 0.4808 | 0.5726 | 0.6336 |
| **138** | Pro | C | | | 0.9268 | | 1.9845 | 2.3270 |
| **138** | Pro | Ca | 0.1237 | 0.2059 | 0.0564 | 0.1675 | 0.1183 | 0.2417 |
| **138** | Pro | Cb | 0.2900 | 0.2671 | 0.1211 | 0.0857 | 0.8339 | 0.9269 |
| **138** | Pro | Cd | 0.1150 | 0.0550 | 0.0003 | | 0.0887 | 0.1148 |
| **138** | Pro | Cg | | 0.2696 | 0.0817 | 0.2445 | 0.0513 | 0.0009 |
| **138** | Pro | N | 0.1062 | 0.1295 | 0.2227 | 0.2777 | 0.0339 | 0.0549 |
| **139** | Ile | C | 1.8585 | | | | | |
| **139** | Ile | Ca | 1.0108 | | | 1.1348 | | |
| **139** | Ile | Cb | 2.1820 | | | 3.1049 | | |
| **139** | Ile | Cg2 | 2.0284 | | | 1.6995 | | |
| **139** | Ile | N | 0.2451 | | | 2.4079 | | |
| **140** | Leu | Ca | 2.6186 | | | | | |
| **140** | Leu | Cg | 0.0117 | | | | | |
| **140** | Leu | N | 0.3941 | | | | | |

(3) Fogeron, M.-L.; Lecoq, L.; Cole, L.; Harbers, M.; Böckmann, A. Easy Synthesis of Complex Biomolecular Assemblies: Wheat Germ Cell-Free Protein Expression in Structural Biology. *Front. Mol. Biosci.*

**2021**, *8*, 639587. https://doi.org/10.3389/fmolb.2021.639587.
